# DELLA proteins regulate spore germination and reproductive development in *Physcomitrium patens*

**DOI:** 10.1101/2022.09.07.506957

**Authors:** Alexandros Phokas, Rabea Meyberg, Asier Briones-Moreno, Jorge Hernandez-Garcia, Panida T. Wadsworth, Eleanor F. Vesty, Miguel A. Blazquez, Stefan A. Rensing, Juliet C. Coates

## Abstract

- Proteins of the DELLA family integrate environmental signals to regulate growth and development throughout the plant kingdom. Plants expressing non-degradable DELLA proteins underpinned the development of high-yielding ‘Green Revolution’ dwarf crop varieties in the 1960s. In vascular plants, DELLAs are regulated by gibberellins, diterpenoid plant hormones. How DELLA protein function has changed during land plant evolution is not fully understood.
- We have examined the function and interactions of DELLA proteins in the moss *Physcomitrium* (*Physcomitrella*) *patens*, in the sister group of vascular plants (Bryophytes).
- *Pp*DELLAs do not undergo the same regulation as flowering plant DELLAs. *Pp*DELLAs are not degraded by diterpenes, do not interact with GID1 gibberellin receptor proteins and do not participate in responses to abiotic stress. *Pp*DELLAs do share a function with vascular plant DELLAs during reproductive development. *Pp*DELLAs also regulate spore germination. *Pp*DELLAs interact with moss-specific photoreceptors although a function for *Pp*DELLAs in light responses was not detected. *Pp*DELLAs likely act as ‘hubs’ for transcriptional regulation similarly to their homologues across the plant kingdom.
- Taken together, these data demonstrate that *Pp*DELLA proteins share some biological functions with DELLAs in flowering plants, but other DELLA functions and regulation evolved independently in both plant lineages.

## Introduction

Proteins of the DELLA family, named for their N-terminal conserved amino acid motif in flowering plants, are widespread regulators of plant growth and development (Thomas *et al.*, 2016; Vera-Sirera *et al.*, 2016). In *Arabidopsis*, DELLA proteins restrain growth in response to a range of environmental stimuli including abiotic stresses (Achard *et al.*, 2008) and pathogen attack (Lan *et al.*, 2014). DELLAs function throughout the flowering plant life cycle as key regulators of seed germination, root growth, seedling development, stem elongation, vegetative development flowering and reproductive development (Gao *et al.*, 2017).

One key property of DELLA proteins in flowering plants is their degradation via action of the gibberellin signalling pathway (Fu *et al.*, 2002). Bioactive gibberellins activate GIBBERELLIN INSENSITIVE DWARF 1 (GID1) receptors, which interact with DELLAs to target them for degradation by the proteasome via an F-box protein SLEEPY/GIBBERELLIN INSENSITIVE DWARF 2 (SLY/GID2) (McGinnis *et al.*, 2003; Sasaki *et al.*, 2003; Ueguchi-Tanaka *et al.*, 2005; Ueguchi-Tanaka *et al.*, 2007). Gain-of-function mutations in genes encoding DELLA proteins underpin semi-dwarf ‘Green Revolution’ phenotypes in cereals (Peng *et al.*, 1999). These mutations remove the N-terminal DELLA region producing non-degradable DELLA proteins, which restrain plant stem growth (Peng *et al.*, 1999).

DELLAs act as hubs for a wide range of protein-protein interactions, enabling their multiple functions and their ability to mediate crosstalk between gibberellin signalling and other hormone signalling pathways (Gao *et al.*, 2017; Hernandez-Garcia *et al.*, 2021a; Phokas & Coates, 2021). DELLAs regulate expression of multiple genes via a number of different mechanisms including transcriptional coactivation, transcriptional repression and sequestration of transcriptional regulators (Marin-de la Rosa *et al.*, 2014; Lantzouni *et al.*, 2020; Phokas & Coates, 2021). For example, in their role repressing seed germination, *At*DELLAs interact with the ABSCISIC ACID (ABA)–INSENSITIVE transcription factors *At*ABI3 and *At*ABI5 enabling transcriptional coactivation of the *AtSOMNUS* gene (Lim *et al.*, 2013). During restraint of vegetative growth, *At*DELLAs integrate light- and gibberellin signalling by interacting with the PHYTOCHROME INTERACTING FACTOR (*At*PIF) transcription factors and preventing PIFs binding to the promoters of growth-promoting genes (de Lucas *et al.*, 2008; Feng *et al.*, 2008). In addition, during promotion of male reproductive development, the rice DELLA protein *Os*SLR1 interacts with the MYB protein *Os*MS188 (Jin *et al.*, 2022).

DELLAs have largely been studied in flowering plants. The question of how DELLA protein functions evolved remains largely unanswered. DELLAs are part of a larger family of GRAS proteins (named for two DELLA proteins, GIBBERELLIC ACID INSENSITIVE (GAI) and REPRESSOR OF GA1-3 (RGA), plus the transcription factor SCARECROW). GRAS proteins are present in the closest algal relatives of land plants (Hernandez-Garcia *et al.*, 2019) and may have arisen via horizontal gene transfer (HGT) of GRAS-like genes from bacteria (Zhang *et al.*, 2012). DELLAs are land plant specific and present in all land plant lineages (Hernandez-Garcia *et al.*, 2019). In non-vascular plants (bryophytes), N-terminal DELLA domains in two of the three major clades, liverworts and hornworts, are largely similar to those of flowering plants (Hernandez-Garcia *et al.*, 2019). However, the N-terminal DELLA domain of most mosses diverges considerably from the vascular plant consensus suggesting moss-specific changes as this lineage evolved (Yasumura *et al.*, 2007; Hernandez-Garcia *et al.*, 2019). Despite moss DELLA divergence, DELLA N-terminal protein domains from both *Marchantia polymorpha* (liverwort) and *Physcomitrium patens* (moss) can act as transcriptional activators, similarly to flowering plant DELLA N-termini (Hernandez-Garcia *et al.*, 2019). The single *Marchantia* DELLA, *Mp*DELLA, represses vegetative growth, and via interaction with *Mp*PIF promotes oxidative stress resistance via apical notch survival and flavonoid accumulation, relieves gemma cup dormancy (promoting vegetative reproduction) but restrains sexual reproductive development (conversely to flowering plant DELLAs) (Hernandez-Garcia *et al.*, 2021b). Thus, DELLA-PIF interaction occurred early in land plant evolution, despite *Marchantia* possessing no GID1 gibberellin receptor homologue (Hernandez-Garcia *et al.*, 2021b).

The gibberellin-dependent interaction between DELLAs and GID1 receptors (and subsequent degradation of DELLAs) is thought to have originated in vascular plants (Hirano *et al.*, 2007; Yasumura *et al.*, 2007) suggesting non-vascular plant DELLAs do not play a role in a gibberellin signalling pathway. *Pp*DELLA is not degraded in the presence of an array of gibberellin-like compounds (Hirano *et al.*, 2007; Yasumura *et al.*, 2007). Mutants in the two *Physcomitrium PpDELLA* genes do not suggest a role for *Pp*DELLAs in controlling vegetative growth, including in response to salt stress or the flowering plant gibberellin GA_3_. Overexpression of *Pp*DELLA in *Arabidopsis* but not rice inhibits vegetative growth (Hirano *et al.*, 2007; Yasumura *et al.*, 2007).

Gibberellin or gibberellin signalling components play roles in controlling plant reproduction throughout the plant lineage. In flowering plants, gibberellin-DELLA signalling promotes transition from the vegetative to the reproductive phase of the life cycle and coordination of male and female reproductive organ development (Plackett & Wilson, 2016). *Arabidopsis* and rice DELLAs are required for male reproductive development (Plackett *et al.*, 2014; Jin *et al.*, 2022) while overexpression of *Mp*DELLA or loss of *Mp*PIF causes a delay to reproductive development in *Marchantia* (Hernandez-Garcia *et al.*, 2021b), indicating potential changes to DELLA reproductive function during plant evolution.

Given the relative lack of previous experimentation on *Pp*DELLA proteins using non-flowering plant diterpenes, in this paper we sought to further understand potentially divergent moss DELLA protein functions and interactions in relation to hormone- and environmental cues.

## Methods

### *Physcomitrium patens* tissue culture for maintenance and spore germination analyses

*Physcomitrium patens* ‘Gransden UK’ / Birmingham (Gd-UK / Birmingham) strain (Haas *et al.*, 2020) wild type was used for comparison with *Ppdella* mutant strains provided by Professor Nicholas Harberd, University of Oxford (*Ppdellaa, Ppdellab, Ppdellaab,* (Yasumura *et al.*, 2007))*. Physcomitrium* tissue was cultured for phenotyping and spores germinated as described previously (Moody *et al.*, 2012; Vesty *et al.*, 2016). Sporulation was induced and sporophytes harvested as described in (Moody *et al.*, 2012). Further details are provided in the Supplemental Methods.

### *P. patens* tissue culture for gametangia/sporophyte and crossing analyses

For gametangia analyses, *P. patens* Gd-UK and *Ppdellaab* were cultivated as described in (Hiss *et al.*, 2017). Tissues for crossing analyses (Gd-UK, *Ppdellaab*, Re-mCherry, *Ppccdc39*) were cultivated as described in (Perroud *et al.*, 2018; Perroud *et al.*, 2019; Meyberg *et al.*, 2020). Further details are given in the Supplemental Methods.

### *Arabidopsis* growth

*Arabidopsis* ‘Landsberg *erecta* (L*er*)’ wild-type ecotype and *pRGA::GFP-AtRGA* overexpression line in L*er* background (Achard *et al*., 2006), kindly provided by Professor Nicholas Harberd, University of Oxford, were grown as described in (Nibau *et al.*, 2011).

### Protein sequence alignment and phylogeny

Putative full-length land plant DELLA protein sequences were obtained from Phytozome (https://phytozome-next.jgi.doe.gov/) (Goodstein *et al.*, 2012) using BLASTP or from NCBI ((Altschul *et al.*, 1990); www.ncbi.nlm.nih.gov/BLAST/) using standard protein BLAST, or from the OneKP database ((One Thousand Plant Transcriptomes, 2019); https://db.cngb.org/onekp/) using BLASTP. *At*RGA protein sequence was used as query for all BLAST searches. DELLA protein sequences of *Anthoceros punctatus and Anthoceros agrestis* were obtained from (Li *et al.*, 2020) and DELLA protein sequences of *Ceratopteris richardii* from (Marchant *et al.*, 2019). Sequence alignment was carried out using SeaView software (version 4.7) (Gouy *et al.*, 2010) and presented using BoxShade (version 3.2) (https://embnet.vital-it.ch/software/BOX_form.html) on default settings. Phylogenetic trees of DELLA land plant homologues were created in SeaView software on default settings, using the maximum likelihood algorithm with 100 bootstrap replicates or the BioNJ algorithm with 1000 bootstrap replicates, and displayed using iTOL (https://itol.embl.de/).

### *P. patens* protoplast isolation and transformation

*P. patens* protoplast isolation was carried out as described in (Schaefer *et al.*, 1991) and protoplast transformation as in (Moody *et al.*, 2021).

### *P. patens* genomic DNA extraction

Genomic DNA extraction was carried out as described in (Moody *et al.*, 2021). Further details are given in the Supplemental Methods.

### *P. patens* RNA extraction, cDNA synthesis and reverse transcription PCR

*P. patens* RNA was isolated as described in (Vesty *et al.*, 2016). *P. patens* cDNA was synthesised using the Tetro cDNA synthesis kit (Bioline) with OligodT primers as per manufacturer’s instructions. Reverse transcription PCR (RT-PCR) was performed as described in (Vesty *et al.*, 2016).

### PCR and sequencing primers

All primers are listed in Supplemental Table 5. PCR primers were designed using the NCBI primer designing tool (https://www.ncbi.nlm.nih.gov/tools/primer-blast/).

### Generation of *PpDELLA* inducible overexpression constructs

*PpDELLAa and PpDELLAb* coding sequences (excluding start codons) were PCR-amplified from *P. patens* genomic DNA using *XhoI-PpDELLAa_pHSP-F* with *SalI-PpDELLAa_pHSP-R* and *XhoI-PpDELLAb_pHSP-F* with *SalI-PpDELLAb_pHSP-R,* respectively. The PCR products were ligated into pCR-blunt using the Zero Blunt PCR cloning kit (Thermo Fisher Scientific) as per manufacturer’s instructions, and then sequenced (Eurofins, Wolverhampton, UK) using the primers *M13_F, XhoI-PpDELLAa_pHSP-F*, *PpDELLAa_internal_F* and *M13_R* for *PpDELLAa,* and *M13_F*, *XhoI-PpDELLAb_pHSP-F*, *PpDELLAb_internal_F* and *M13_R* for *PpDELLAb*. *PpDELLAa* and *PpDELLAb* were then digested out of pCR-blunt using *Xho*I and *Sal*I and ligated into *pHSP-MCS-GFP-108-35SNPT* (GenBank: KP893621.1; (Moody *et al.*, 2016)).

### Genotyping of *P. patens* transformants

PCR was used to screen transformants for the presence of *pHSP::PpDELLAa-GFP, pHSP::PpDELLAb-GFP* or *pHSP::GFP.* For *pHSP::PpDELLAa-GFP* and *pHSP::PpDELLAb-GFP,* the primers *pHSP_F* and *mGFP_R* were used. In addition, for *pHSP::PpDELLAa-GFP* the primer *mGFP_R* was also used in combination with the primer *XhoI-PpDELLAa_pHSP-F*. For *pHSP::GFP*, the primers *pHSP_F* and *35STer_R* were used. All transformants were also genotyped using primers *nptII_F* and *108locus5’_R* to confirm integration into the inert *108* genomic locus.

### Plant protein expression analysis

For *P. patens* protein expression analysis, 7-day or 14-day old protonemata transformed with *pHSP::PpDELLAa-GFP-108-35SNPT*, *pHSP::PpDELLAb-GFP-108-35SNPT* or *pHSP::GFP-108-35SNPT* were incubated in liquid BCD supplemented with 1 mM CaCl_2_ and 5 mM ammonium tartrate in 24-well plates for 1h at 37°C (protein induction) or at 22±1°C (control) with gentle agitation, followed by incubation at 22±1°C for at least 6h with gentle agitation. Normally, half a plate of *P. patens* tissue was transferred to each well, which contained 1ml of liquid BCD supplemented with 1 mM CaCl_2_ and 5 mM ammonium tartrate. For testing the effects of GA_3_, GA_9_-ME, *ent*-kaurenoic acid, paclobutrazol, uniconazole or methanol on protein stability, the chemical was added to the well following the 6-hour post-heat-shock incubation period. For *Arabidopsis pRGA::GFP-AtRGA*, 7-day old seedlings were incubated in 24-well plates in liquid ½ Murashige and Skoog (MS) medium (2.2g MS basal medium [Sigma-Aldrich, M0404] in 1L dH_2_O, pH 5.7) supplemented with GA_3_, GA_9_-ME or methanol. For protein extraction, *P. patens* or *Arabidopsis* tissue was collected in 50ml Falcon tubes or 1.5ml Eppendorf tubes, flash frozen in liquid nitrogen and stored at −80°C. For live-cell imaging analysis, confocal microscopy was used.

### Confocal microscopy

Confocal images or z-stacks of protein expression in *P. patens* protonemata or *Arabidopsis* roots were captured with the Zen 2012 software using the Zeiss LSM170 confocal microscope (20× objective). Wild-type protonemata and protonemata with uninduced GFP expression were compared with protonemata with induced *Pp*DELLA-GFP protein expression using identical laser power, gain and pinhole settings. For *Arabidopsis*, L*er* roots were compared to *pRGA::GFP-AtRGA* roots. Excitation and emission wavelengths for GFP fluorescence were 488nm and 530nm respectively and for chloroplast autofluorescence 634nm and 696nm respectively.

### Plant protein extraction

A mortar and a pestle (pretreated with 70% ethanol) were used to grind up frozen plant tissue in liquid nitrogen. As soon as the ground tissue had reached room temperature, protein extraction buffer (50mM Tris/HCl pH 7.5 or HEPES pH 7.5, 150mM NaCl, 5% glycerol, 0.5% NP-40, cOmplete™ EDTA-free protease inhibitor tablets [Roche] - one per 10ml buffer) was added to the tissue and grinding was continued until a homogenous suspension was formed. The lysate was then filtered through a single layer of miracloth (Millipore) into 50ml falcon tubes or 1.5ml Eppendorf tubes on ice. The lysate was centrifuged at 14000 g for 30-45 minutes at 4°C and the supernatant transferred into fresh tubes. For protein analysis by Western blotting, the protein extract was diluted in 5× Laemmli buffer (10% [w/v] SDS, 50% [w/v] glycerol, 5% β-mercaptoethanol, 0.005% [w/v] bromophenol blue) and boiled for 10 minutes at 95°C. For *P. patens* protein extraction destined for immunoprecipitation coupled to mass spectrometry, four plates of 17-day old protonemata carrying either *pHSP::PpDELLAa-GFP* or *pHSP::GFP* were incubated for 1h at 37°C to induce protein expression, followed by incubation at 22±1°C for 6h. Four plates of 17-day old protonemata carrying *pHSP::PpDELLAa-GFP* were also continuously incubated at 22±1°C without undergoing incubation at 37°C (control). *P. patens* tissue was then flash frozen in 50ml Falcon tubes in liquid nitrogen and stored at −80°C. Frozen tissue from the three treatment groups was ground in liquid nitrogen, and then mixed with 4ml protein extraction buffer (50mM Tris/HCl pH 7.5, 5% glycerol, 150mM NaCl, 0.2% triton X-100, cOmplete™ EDTA-free protease inhibitor tablets [Roche] - one per 10ml buffer) as soon as the ground tissue had reached room temperature. Grinding was continued until a homogenous suspension was formed, which was filtered successively through a double and a single layer of miracloth (Millipore) into 50ml Falcon tubes on ice. This was followed by two rounds of centrifugation in 2ml Eppendorf tubes at 12000 g for 30 minutes each at 4°C. Supernatants were collected into single 15ml Falcon tubes and total protein adjusted to 4.8mg in 3.7ml protein extraction buffer, made up to 7.4ml with dilution buffer (10mM Tris/HCl pH 7.5, 0.5mM EDTA, 150mM NaCl, cOmplete™ EDTA-free protease inhibitor tablets [Roche] - one per 10ml buffer) and stored in −80 °C until immunoprecipitation was performed.

### Yeast protein extraction

AH109 yeast was cultured overnight in liquid synthetic amino acid Drop out (DO) - leu-trp at 30°C. 1ml yeast culture with OD600 8 was centrifuged for 4 minutes at 12000 g. The pellet was resuspended in 50μl Buffer A (0.1M NaOH, 50mM EDTA, 2% SDS, 2% β-mercaptoethanol) and incubated at 90°C for 10 minutes. The suspension was supplemented with 0.67μl 3M acetic acid, vortexed for 30s on a Titrtek and for 1 minute on a vortex, and incubated for 10 minutes at 90°C. This was followed by addition of 12.5μl of Buffer B (250mM Tris pH6.8, 50% glycerol, 0.05% bromophenol blue), brief vortexing and centrifugation at 12000 g for 5 minutes. 55μl supernatant was transferred to a fresh tube and boiled for 1 minute at 98°C. Protein expression was analysed by SDS-PAGE and western blotting.

### SDS-PAGE and Western blotting

SDS-PAGE and Western blotting were performed as described in Gibbs *et al*. (2014). Further details are given in the Supplemental Methods.

### Generation of Yeast two-hybrid constructs

*PpDELLAs* or *AtRGA* cloned in pGADT7 and *PpGLP1* or *AtGID1c* cloned in pGBKT7 (Yasumura *et al.*, 2007) were kindly provided by Professor Nicholas Harberd, University of Oxford. *PpPHY5B (Pp3c12_9240), PpPHOTA2 (Pp3c21_21410)* and *PpPHOTB1 (Pp3c2_10380)* coding sequences (excluding the start codons) were PCR-amplified from *P. patens* cDNA (from gametophore tissue) using *NdeI-PpPHY5B_F* with *NotI-PpPHY5B_R*, *SalI-PpPHOTA2_F* (including an additional TT after the restriction site to enable in-frame cloning) with *NotI-PpPHOTA2_R*, and *NdeI-PpPHOTB1_F* with *NotI-PpPHOTB1_R*, respectively. The PCR products were ligated into pCR-blunt using the Zero Blunt PCR cloning kit (Thermo Fisher Scientific) as per manufacturer’s instructions, and then sequenced (Eurofins, Wolverhampton, UK) using the primers *M13_F* and *M13_R*. *PpPHY5B, PpPHOTA2* and *PpPHOTB1* cDNA (excluding the start codons) was PCR-amplified using pCR-blunt as template and *NdeI-PpPHY5B_F* with *NotI-PpPHY5B_R*, *SalI-PpPHOTA2_F* with *NotI-PpPHOTA2_R*, and *NdeI-PpPHOTB1_F* with *NotI-PpPHOTB1_R,* respectively. The PCR products were digested with *Nde*I and *Not*I for *PpPHY5B* and *PpPHOTB1* or *Sal*I and *Not*I for *PpPHOTA2*, and ligated into pGBKT7 (Clontech, USA), where they were sequenced (Eurofins, Wolverhampton, UK). For *PpPHY5B*, the following sequencing primers were used: *T7_F*, *NdeI-PpPHY5B_F, PpPHY5B-Seq_F, PpPHY5B-Seq2_F, PpPHY5B-Seq3_F*. For *PpPHOTA2,* the following sequencing primers were used: *T7_F*, *M13_R*, *SalI-PpPHOTA2_F, PpPHOTA2-Seq_F, PpPHOTA2-Seq2_F, PpPHOTA2-Seq3_F.* For *PpPHOTB1,* the following sequencing primers were used: *T7_F*, *M13_R*, *NdeI-PpPHOTB1_F, PpPHOTB1-Seq_F, PpPHOTB1-Seq2_F* and *PpPHOTB1-Seq3_F*.

### Yeast two-hybrid assays

Transformation of AH109 yeast with the appropriate plasmids was carried out as described in (Gibbs *et al.*, 2014). Further details are given in the Supplemental Methods.

### Co-Immunoprecipitation (Co-IP) in a cell-free system

MYC-tagged *At*GID1c or *Pp*GLP1 expressed from pGBKT7 and HA-tagged *At*RGA or *Pp*DELLAa or *Pp*DELLAb expressed from pGADT7 (Yasumura *et al.*, 2007) were translated *in vitro* using the TNT® T7 Coupled Reticulocyte Lysate System (Promega, Madison, WI, USA) according to the manufacturer’s instructions. Further details are given in the Supplemental Methods.

### *P. patens* spore culture and germination assays

Spores were cultured and germination assays were performed as described in (Vesty *et al.*, 2016).

### *P. patens* vegetative tissue growth assays under abiotic stress conditions

*P. patens* vegetative growth under salt and oxidative stress was assayed by measuring plant area after culture plates were photographed using a Nikon D40 SLR camera. Individual plants of similar size were transferred onto BCDATG agar (4-5 per plate) supplemented with NaCl or methyl viologen (Sigma-Aldrich, 856177) and incubated at 22±1°C with a 16h:8h light:dark cycle. Plant area measured at the start of the assay was subtracted from that measured at the end of the assay. For measurements, the polygon function in Fiji software was used. For desiccation stress, *P. patens* vegetative growth was assayed by transferring cellophanes carrying 7-day old protonemata growing on BCD agar supplemented with 1 mM CaCl_2_ and 5 mM ammonium tartrate (BCDAT) onto fresh BCDAT agar plates supplemented with ABA or methanol and incubating them overnight at 22±1°C with a 16h:8h light:dark cycle. Following this, cellophanes were transferred into empty petri dishes and incubated under the same conditions for 7 days, after which time they were transferred onto fresh BCDAT agar plates for another 7 days to allow tissue recovery. The effect of desiccation stress was assayed qualitatively by observing plant physical appearance at the end of the assay.

### Sporophyte development assays

For sporophyte development assays, *P. patens* protonemata were homogenised and cultured on BCDAT agar supplemented with 1 mM CaCl_2_ and 5 mM ammonium tartrate for 7-14 days. Individual moss plants of similar size were then transferred aseptically onto cellophane-overlaid BCDATG agar plates. Plates were incubated at 22±1°C with a 16h:8h light:dark cycle for 4 weeks, and plants transferred aseptically inside sterile Magenta™ vessels (Merck, CO542) containing BCD agar supplemented with 1 mM CaCl_2_ (one plant per vessel). Plants were incubated at 15°C with an 8h:16h light:dark cycle for 7 weeks to induce reproductive development. Photographs were taken using a Nikon D40 SLR camera and sporophyte density was quantified by counting the number of sporophytes per plant tissue area, estimated using the polygon function in Fiji software.

### Gametangia, sporophyte and crossing analyses

For gametangia analyses, apices of adult gametophores (21d after SD transfer, timepoint of mature antheridia and archegonia) were uncovered and transferred on an objective slide with sterile tap water. Sporophytes were counted and analysed 2-3 weeks after watering for crossed sporophytes using a Leica MZ10F binocular to detect mCherry fluorescence in Re-mCherry crossed sporophytes. Samples were covered with coverslips and subsequently analysed with a Leica DM6000 microscope equipped with a DFC295 camera (Leica, Wetzlar, Germany) using the Leica Application Suite 4.4 with the Multi-Stack Module. For sporophyte and crossing statistical analyses Microsoft Excel (Microsoft, USA) was used to determine mean and standard error as well as to visualize data. To determine sporophytes per gametophore and the number of crossed sporophytes per total sporophytes, at least 100 gametophores per strain were analysed. To arrange and modify brightness and contrast of microscopic images, Microsoft Power Point was used (Microsoft, USA).

### Immunoprecipitation coupled to mass spectrometry

Immunoprecipitation was carried out using the GFP-trap® magnetic agarose kit (Chromotek, Germany) as per manufacturer’s instructions. Further details are given in the Supplemental Methods. Following SDS-PAGE, gels were stained using the ProteoSilver™ silver staining kit (Sigma-Aldrich) as per manufacturer’s instructions. Following staining, each run sample was split into two gel pieces. A strong band present in the middle of all samples was used as a reference for excising. Gel pieces excised were submitted for trypsin digest and LC/MS analysis at the School of Biosciences Proteomics Facility, Birmingham. Orbitrap Elite and Q-Exactive HF mass spectrometers were used. Peptides were matched against Uniprot database and proteins were identified with cut-off protein false discovery rate (FDR) set to 1%.

### *P. patens* light assays

For spore or vegetative tissue growth assays under different light wavelengths, *P. patens* was incubated on BCDATG agar (for vegetative tissue) or BCD agar supplemented with 5 mM CaCl_2_ and 5 mM ammonium tartrate (for spores) plates inside cardboard boxes at 22±1°C under continuous illumination from the top by either white LED lights, filtered via neutral density or colour filters, or by far-red LED lights (LumitronixR, Germany). White light (63μmolm^−2^s^−1^) was filtered using overlaid single layers of neutral density filters No. 209 and No. 298 (LEE filters, UK), red light (640-695nM, 26μmolm^−2^s^−1^) was filtered using two layers of Deep golden amber filter No. 135 (LEE filters, UK) and blue light (445-490nM, 16μmolm^−2^s^−1^) using three layers of Moonlight blue filter No. 183 (LEE filters, UK). Far-red light (730nM) was adjusted to 16μmolm^−2^s^−1^. The effect of light wavelength on vegetative tissue growth was assayed qualitatively by observing physical appearance, e.g. gametophore development or wilting after culture plates were photographed using a Nikon D40 SLR camera.

### RNA preparation and RNAseq analysis

The samples consisted of *P. patens* gametophores grown in BCDAT for 3 weeks. Total RNA was extracted from triplicate samples with the NucleoSpin™ RNA Plant Kit (Macherey-Nagel), and the RNA concentration and integrity (RIN) were measured in an RNA nanochip (Bioanalyzer, Agilent Technologies 2100). The preparation of libraries and subsequent sequencing in an Illumina NextSeq 500 platform was carried out at Beijing Genomics Institute (BGI) yielding at least 20M 100-bp paired-end reads per sample. The read qualities were explored using FastQC version 0.11.9. The adaptors were removed from the reads processing the paired-end files together using bbduk version 38.42 with the default adapters file and the following parameters: “ktrim=r k=23 mink=11 hdist=1”. Next, the reads were quality filtered using Trimmomatic (Bolger *et al.*, 2014) version 0.39 with the following parameters: “-phred33 LEADING:3 TRAILING:3 SLIDINGWINDOW:4:15 MINLEN:35” and the quality of the filtered files was assessed with FastQC.

For the differential expression analysis, the *P*. *patens* annotation version 3.3 was downloaded from Phytozome (Goodstein *et al.*, 2012). An index file was created using the index command from *Salmon* version 1.1.0 (Patro *et al.*, 2017). The number of reads per transcript was determined with *salmon quant* using the – validateMappings parameter and the filtered reads file. Finally, the differential expression analysis was performed using DESeq2 (Love *et al.*, 2014).

GO enrichment analysis was carried out with the corresponding tool in PlantRegMap (Tian *et al.*, 2020) and the categories were organized and represented with ReviGO (Supek *et al.*, 2011). TF enrichment analysis was performed using the TF enrichment tool available at PlantRegMap (Tian *et al.*, 2020) A TF is considered enriched if the number of possible targets for it on the input list of genes is higher than expected; and a gene is considered a target if there is experimental evidence or it has *cis* regulatory elements or binding motifs for the TF. Log fold-change for DEGs and TF enrichment were visualised using Cytoscape (Shannon *et al.*, 2003).

## Results

### DELLA proteins in *Physcomitrium patens* are part of monophyletic bryophyte and moss groups but are divergent from other plant DELLAs

We placed bryophyte DELLAs in a phylogeny with a selection of full-length bryophyte, lycophyte, fern, gymnosperm and flowering plant DELLA protein sequences (Fig. 1a; Supplemental Fig. 1). Bryophyte DELLAs form a monophyletic group within which resides a moss DELLA clade. Hornwort DELLAs are basal within the bryophyte clade in this analysis. The two *Physcomitrium* DELLAs appear to be the result of a relatively recent genome duplication (Fig. 1a). Alignment of both the N-terminal DELLA-LEQLE domain and the VHYNP domain that are conserved in flowering plants (Yasumura *et al.*, 2007) demonstrates that all selected mosses show considerable divergence in the DELLA and VHYNP domains compared to other bryophytes and vascular plants, while *Sphagnum fallax* and *Marchantia* show the most divergence in the LEQLE region (Fig. 1b). In contrast, the GRAS domain of moss DELLA proteins (including the LHR1 domain; Fig.1c) does not appear to be more divergent than other bryophyte- or vascular plant DELLAs (Fig.1c; Supplemental Fig. 1). These data suggest that moss DELLAs may have some conserved roles (involving the GRAS domain) and some divergent functions (involving the N-terminal DELLA-LEQLE-VHYNP domains).

**Figure 1.**
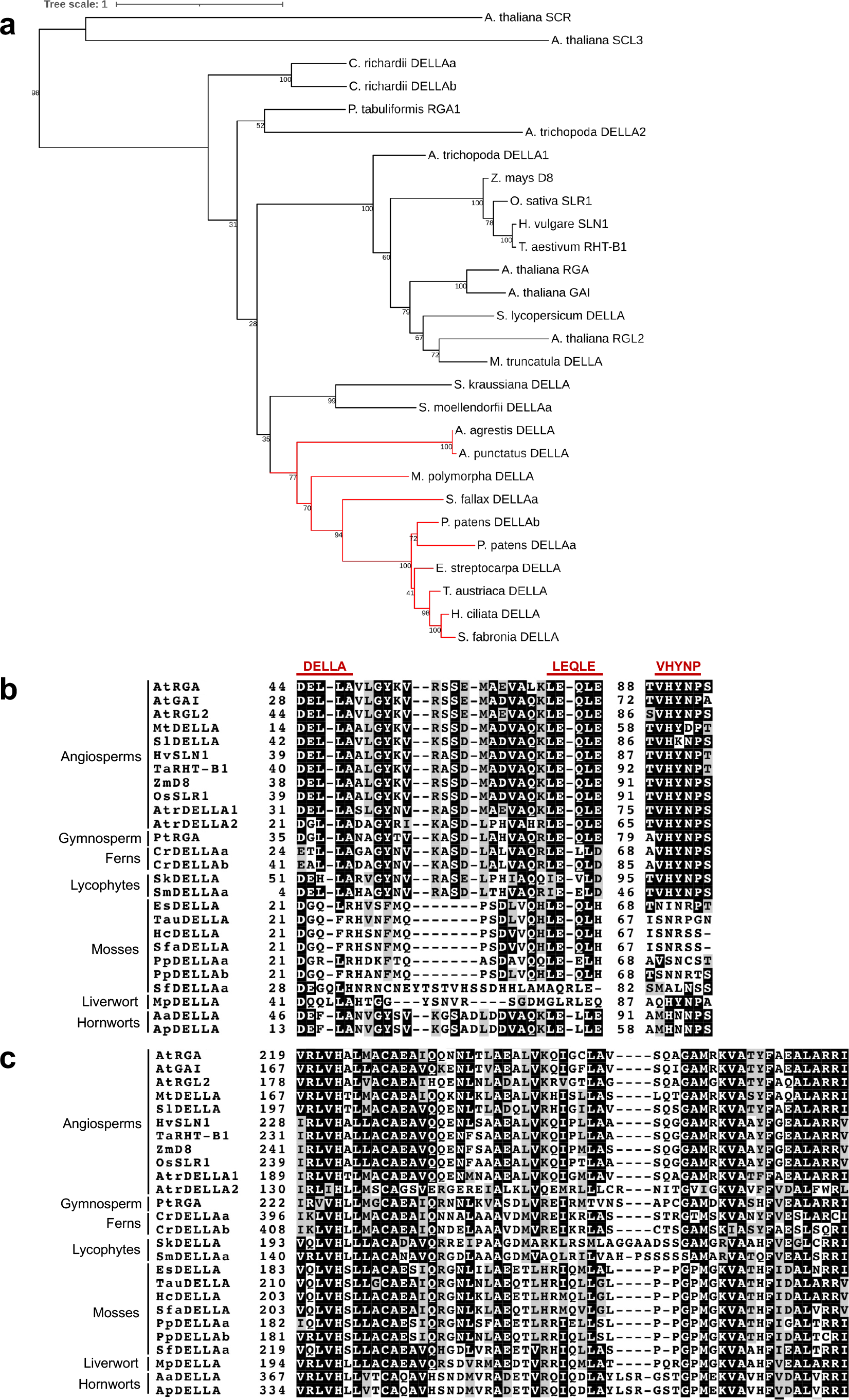
*Pp*DELLAs form a monophyletic group with other bryophyte DELLAs and have a divergent DELLA-LEQLE-VNHYP domain compared to vascular plants with a conserved LHR1 region in the GRAS domain. (A) Maximum Likelihood phylogenetic tree generated using the peptide sequences of selected DELLA homologues from bryophytes, lycophytes, ferns, gymnosperms and angiosperms. The monophyletic bryophyte group is shown in red. Scale bar: 0.1 substitutions per amino acid site. (B) Alignment of the N-terminal DELLA-LEQLE-VHNYP domain that is necessary for the interaction with GID1 receptors in angiosperms. (C) Alignment of the LHR1 region of the GRAS domain demonstrates similarity with GRAS domains from vascular plants and other bryophytes. In (B) and (C) black shading indicates that at least 50% of the amino acids in a particular column are identical. Amino acids that are similar to the column-consensus peptide are shaded grey. The peptide sequences used in this figure are as follows: *Arabidopsis thaliana, Medicago truncatula, Solanum lycopersicum, Hordeum vulgare, Triticum aestivum, Zea mays, Oryza sativa, Amborella trichopoda* (angiosperms)*, Pinus tabuliformis* (gymnosperm)*, Ceratopteris richardii* (fern)*, Selaginella kraussiana, Selaginella moellendorfii* (lycophytes)*, Encalypta streptocarpa, Timmia austriaca, Hedwigia ciliata, Schwetschkeopsis fabronia, Physcomitrium patens, Sphagnum fallax* (mosses)*, Marchantia polymorpha* (liverwort)*, Anthoceros agrestis* and *Anthoceros punctatus* (hornworts).

### *Pp*DELLAs are not degraded in response to diterpenes

Given the divergent N-terminal sequences of *Pp*DELLA proteins, we explored the functions of moss N-terminal DELLA domains further by analysing the stability of *Pp*DELLA proteins compared to flowering plant DELLAs. We generated transgenic *Physcomitrium* lines expressing either *Pp*DELLAa or *Pp*DELLAb as C-terminal fusion proteins with green fluorescent protein (GFP) under the control of an inducible heat shock promoter (Saidi *et al.*, 2005), named *pHSP*::*Pp*DELLAa-GFP and *pHSP*::*Pp*DELLAb-GFP, in addition to control lines expressing *pHSP*::GFP (Supplemental Fig. 2 and 3). Upon heat shock induction of *Pp*DELLAa-GFP and *Pp*DELLAb-GFP proteins both localise to the nucleus in *Physcomitrium* protonemal cells, with *Pp*DELLAb being expressed more strongly and *Pp*DELLAa showing additional punctate cytosolic localisation (Fig. 2a). Neither *Pp*DELLAa-GFP nor *Pp*DELLAb-GFP appear to be degraded upon treatment with 10μM GA_9_-methyl ester or 10μM GA_3_, whilst specific GA_3_-dependent degradation of an *Arabidopsis* GFP-*At*RGA fusion protein expressed from the RGA promoter (Achard *et al.*, 2006) is observed (Fig. 2a, Fig. 2b). Moreover, no differences in *Pp*DELLAa-GFP or *Pp*DELLAb-GFP stability are seen upon treatment with 10μM *ent*-kaurene, or the *ent*-kaurene oxidase inhibitors uniconazole (10μM) or paclobutrazol (10μM). This suggests that *Pp*DELLA protein levels are not affected by changes in diterpene levels caused by exogeneous application of diterpenes or diterpene biosynthesis inhibitors.

**Figure 2.**
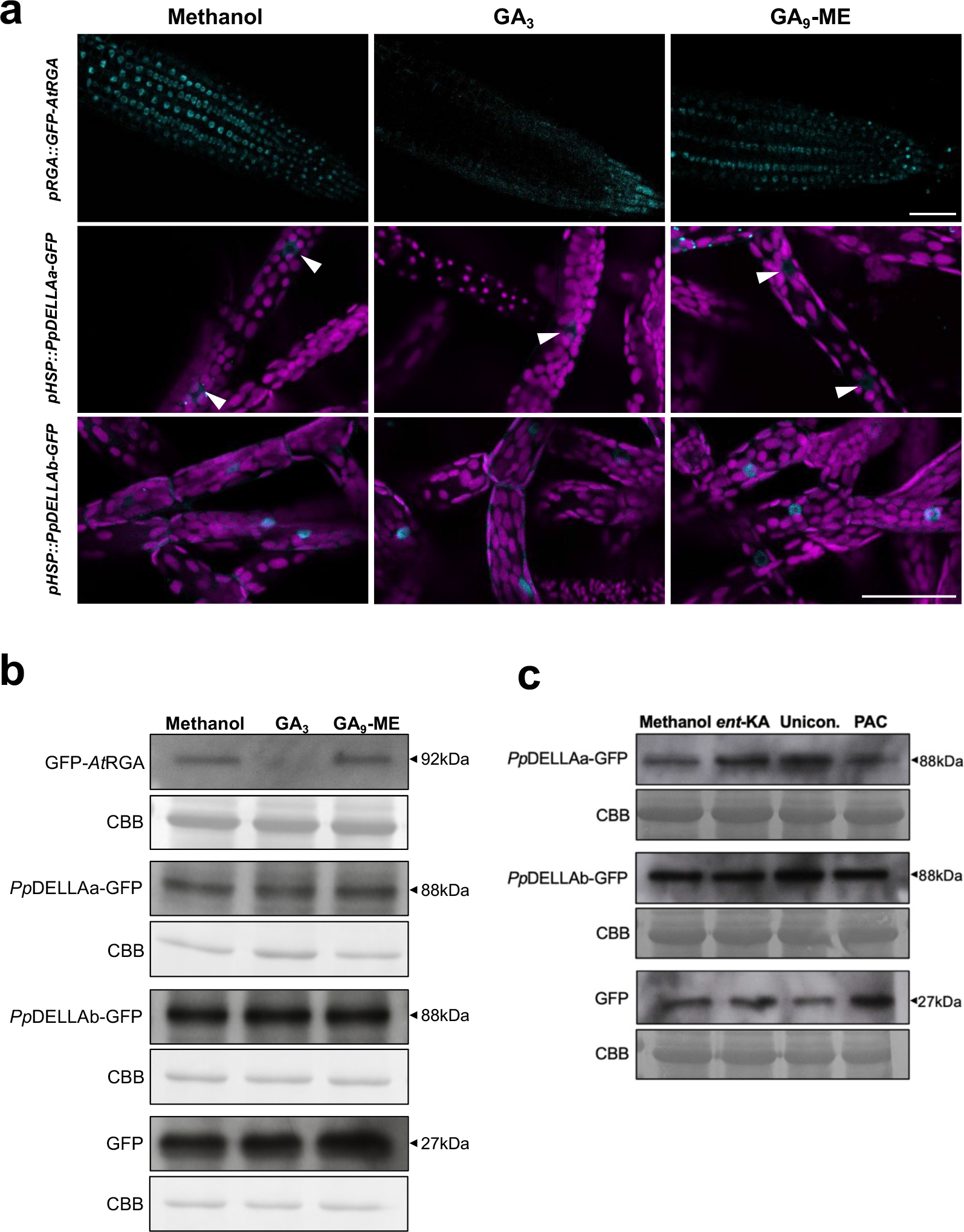
*Pp*DELLA proteins are neither degraded by diterpenes nor stabilised by gibberellin biosynthesis inhibitors. (A) Top panels: GFP-*At*RGA is degraded in 7-day old *Arabidopsis* (*pRGA::GFP-AtRGA*) roots following 2-hour incubation with 10μM gibberellin A_3_ (GA_3_). Middle panels: *Pp*DELLAa-GFP is not degraded in 7-day old *P. patens* protonemata following 2-hour incubation with either 10μM GA_3_ or 10μM GA_9_ methyl ester (GA_9_-ME). White arrowheads: nuclear *Pp*DELLAa-GFP. Bottom panels: *Pp*DELLAb-GFP is not degraded in 7-day old *P. patens* protonemata following 2-hour incubation with either 10μM GA_3_ or 10μM GA_9_-ME. Scale bar: 50μM. (B) GFP-*At*RGA is degraded in 7-day old *Arabidopsis* (*pRGA::GFP-AtRGA*) roots following 2-hour incubation with 10μM GA_3_ while *Pp*DELLAa-GFP and *Pp*DELLAb-GFP are not degraded in 7-day old *P. patens* protonema tissue following 2-hour incubation with either 10μM GA_3_ or 10μM GA_9_-ME. CBB, Coomassie brilliant blue staining. (C) *Pp*DELLAa-GFP and *Pp*DELLAb-GFP levels are not changed in 7-day old *P. patens* protonema tissue following 2-hour incubation with the diterpene *ent*-kaurenoic acid (ent-KA; 10μM) or the GA biosynthesis inhibitors uniconazole (Unicon.; 10μM) or paclobutrazol (PAC; 10μM). CBB, Coomassie brilliant blue staining.

### *Pp*DELLAs do not interact with GID1 receptor homologues from *Arabidopsis* or *Physcomitrium* in the presence of bryophyte-active diterpenes

As *Pp*DELLA protein stability is not affected by diterpenes and previous studies demonstrated that *Pp*DELLAs could not interact with GID1 homologues from *Physcomitrium* or *Arabidopsis* in the presence of gibberellins (Hirano *et al.*, 2007; Yasumura *et al.*, 2007), an obvious hypothesis is that no link exists between DELLA protein function and gibberellin-like signalling in mosses. To build on previous studies, we tested the possibility of an interaction between *Arabidopsis* and moss DELLA-receptor pairs both within- and between species (both *Pp*DELLAs with the representative putative receptor *Pp*GLP1, both *Pp*DELLAs with *At*GID1c, *At*RGA with *Pp*GLP1 and *At*RGA with *At*GID1c) in the presence of GA_3_, GA_9_ methyl ester and *ent*-kaurenoic acid (Fig. 2). The only interaction detected both via yeast two-hybrid assays (Fig. 3a) and via pull-down assays in a cell-free system (Fig. 3b,c) was the known interaction between *At*RGA and *At*GID1c in the presence of GA_3_. This strongly supports the hypothesis that DELLAs and GID1-like proteins do not interact in moss and that there is no link between diterpene function and DELLA protein function in moss.

**Figure 3.**
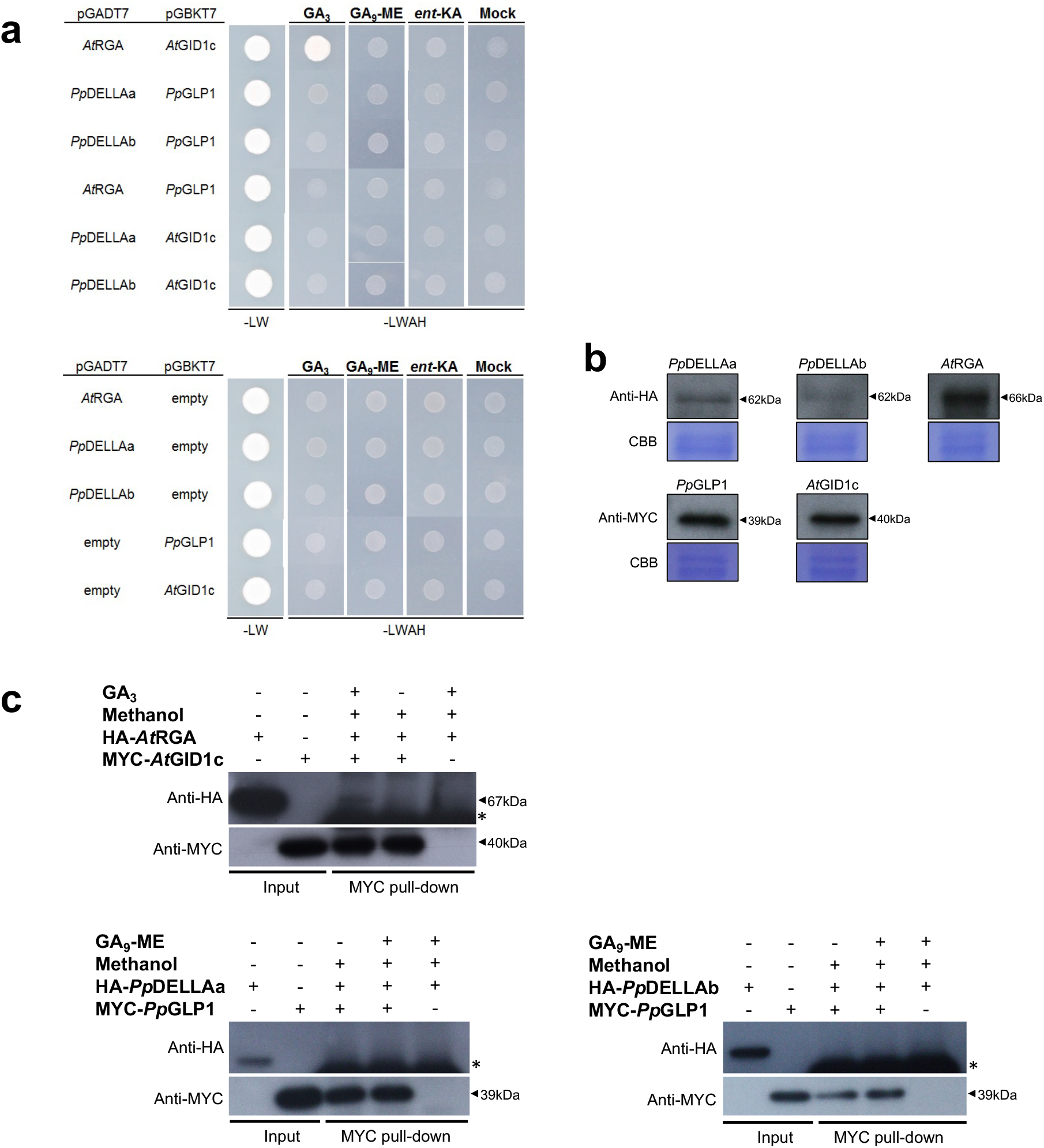
*Pp*DELLAs do not interact with a moss GID1-like protein or with an *Arabidopsis* GID1 gibberellin receptor and an *At*DELLA protein does not interact with a putative moss GID1-like protein. (A) Top panel: in a yeast two-hybrid assay*, Arabidopsis At*RGA and *At*GID1c interact with one another only in the presence of the gibberellin GA_3_, but not GA_9_ methyl ester (GA_9_-ME) or *ent*-kaurenoic acid (*ent*-KA) or a solvent control (Mock). In the same system, *Pp*DELLAa and *Pp*DELLAb do not interact with *Pp*GLP1 including in the presence of diterpenes, *At*RGA does not interact with *Pp*GLP1 and *Pp*DELLAa/*Pp*DELLAb do not interact with *At*GID1c under any conditions. All DELLA proteins were cloned into the yeast vector pGADT7 while GID1 proteins were cloned into the pGBKT7 vector. Bottom panel: No autoactivation is seen when each construct is transformed into yeast alongside the corresponding empty vector. In both panels, −LW is growth in the absence of leucine and tryptophan (to test for the presence of the plasmids) while –LWAH is growth in the absence of leucine, tryptophan, adenine and histidine which tests additionally for the protein-protein interaction. (B) Western blot of yeast cell extracts confirming that HA-tagged *Pp*DELLAs and *At*RGA and MYC-tagged *Pp*GLP1 and *At*GID1 are expressed in yeast: DELLA proteins detected using anti-HA and receptor proteins detected using anti-MYC. CBB: Coomassie Brilliant Blue staining. (C) Coimmunoprecipitation from an *in vitro* cell free system using α-MYC-coupled beads. HA-*At*RGA and MYC-*At*GID1c interacted in a GA_3_-dependent manner, whereas HA-*Pp*DELLAs and MYC-*Pp*GLP1 did not interact in the presence or absence of GA_9_-ME (*, antibody heavy chain).

### *Pp*DELLAs restrain spore germination, while diterpenes promote spore germination in a *Pp*DELLA-independent manner

Because of the function of diterpenes in promoting *Physcomitrium* spore germination and the existence of some parallels between seed- and spore germination (Vesty *et al.*, 2016), and the fact that DELLAs regulate seed germination (Tyler *et al.*, 2004), we investigated whether *Pp*DELLAs were involved in spore germination. We analysed *PpDELLAa* and *PpDELLAb* expression by RT-PCR during the spore germination process (dry spores, imbibed spores and germinating spores) compared to *PpDELLA* expression in protonemal tissue and leafy gametophores (Supplemental Fig. 4a). *PpDELLAa* is most strongly expressed in dry spores, with expression reducing considerably after spore imbibition and absent from germinating spores (Supplemental Fig. 4a). *PpDELLAa* expression is present in older protonemal tissue and then increases in leafy tissue but not to such a high level as in dry spores (Supplemental Fig. 4a). *PpDELLAb* expression is much weaker than *PpDELLAa* in the tissues tested (also seen in Supplemental fig. 4 b,c) and is barely detectable in leafy tissue by RT-PCR and undetectable in other tissues (Supplemental Fig. 4a). The expression pattern of *PpDELLAa* implies function(s) for *PpDELLA*s during spore germination. We found that *Ppdellaa*, *Ppdellab* and *Ppdellaab* mutant spores all germinate faster than wild type spores and that the increased germination rate is similar across all three mutant genotypes (Fig. 4a-c). This demonstrates that *PpDELLAa* and *PpDELLAb* have non-redundant functions restraining spore germination in *Physcomitrium*.

**Figure 4.**
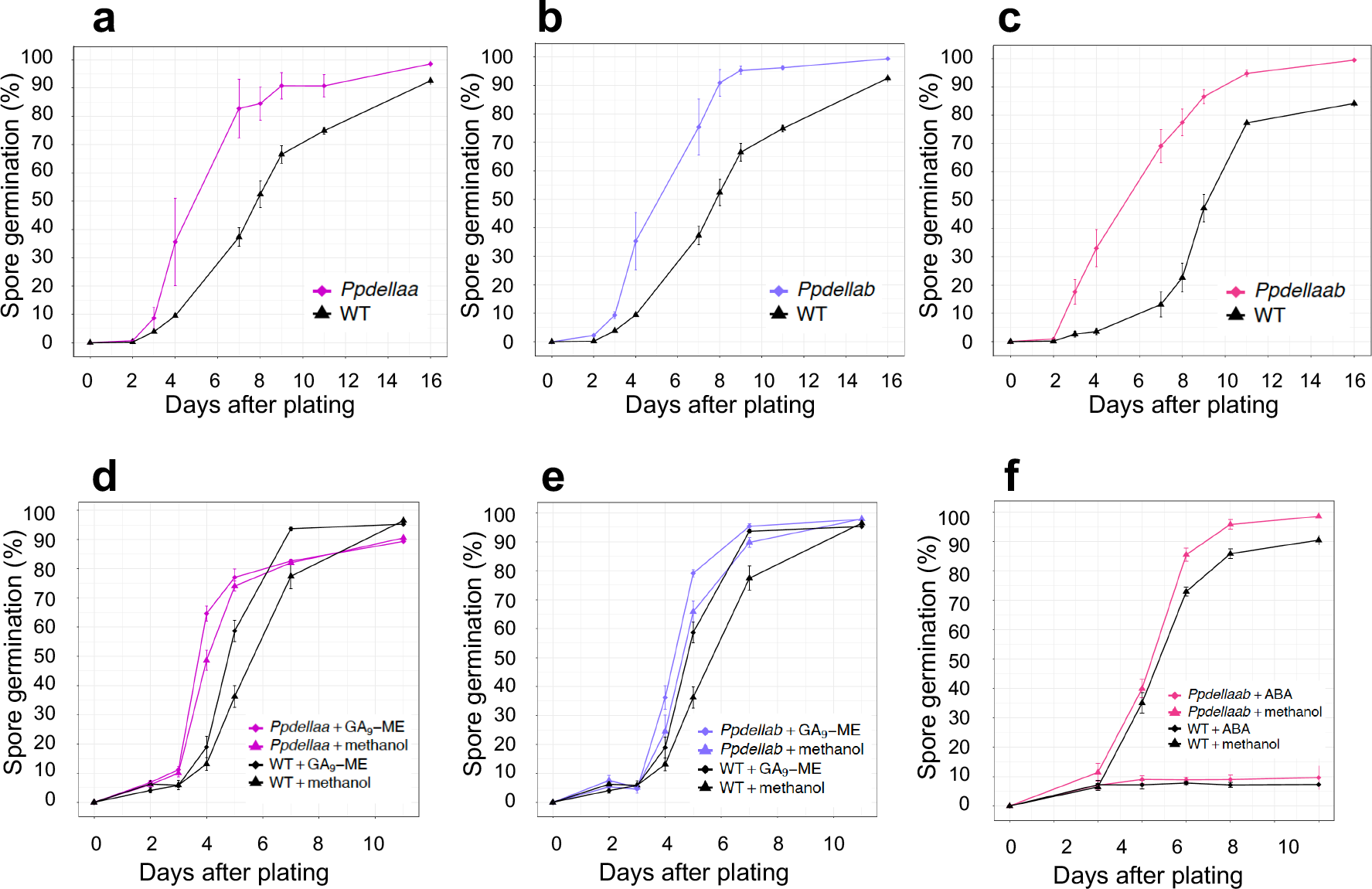
*Ppdella* mutants germinate faster than wild type but are sensitive to application of diterpenes. (A) *Ppdellaa* mutant spores germinate faster than wild type (WT). A Z-test indicates significant differences between *Ppdellaa* and WT on days 4, 7, 8, 8 and 11. (B) *Ppdellab* mutant spores germinate faster than WT. A Z-test indicates significant differences between *Ppdellaa* and WT on days 4, 7, 8, 8 and 11. (C) *Ppdellaab* mutant spores germinate faster than WT. A Z-test indicates significant differences between *Ppdellaa* and WT on days 3, 4, 7, 8, 8, 11 and 16. (D) Treatment with 5μM GA_9_-ME increases spore germination rate of *Ppdellaa* and WT to a similar extent. A Kruskal-Wallis test indicates significant differences between *Ppdellaa* + GA_9_-ME and WT + methanol on days 4 and 5 (p<0.01), between *Ppdellaa* + GA_9_-ME and WT + GA_9_-ME on day 4 (p<0.05), between *Ppdellaa* + methanol and WT + methanol on day 5 (p<0.05), between WT + GA_9_-ME and WT + methanol (p<0.05) on day 7, and between *Ppdellaa* + methanol and WT + GA_9_-ME on day 7 (p<0.05). (E) Treatment with GA_9_-ME increases spore germination rate of *Ppdellab* and WT to a similar extent. A Kruskal-Wallis test indicates significant differences between *Ppdellab* + GA_9_-ME and WT + methanol on days 4, 5 and 7 (p<0.05). (F) Wild type and *Ppdellaab* spores treated with 10μM ABA show a similar extent of germination suppression. A Kruskal-Wallis test indicates significant differences between *Ppdellaab* + ABA and *Ppdellaab* + methanol on days 3, 4, 5 and 7, and between *Ppdellaab* + methanol and WT + ABA on 3, 4, 5 and 7 (p<0.05). All germination assays are representative of 3 or more biological repeats. Error bars, ± SEM.

The absence of a change in DELLA stability in response to diterpenes (Supplemental Fig. 3) and the lack of interaction between *Pp*DELLAs and *Pp*GLP1 (Fig. 3) strongly suggests that diterpenes and *Pp*DELLAs regulate spore germination via independent pathways. To test this suggestion, *Ppdella* mutant spores were germinated in the presence of GA_9_-methyl ester (Fig. 4a, b). GA_9_ methyl ester was able to increase spore germination rate significantly in *Ppdellaa* and *Ppdellab* mutants as in wild-type spores (Fig. 4d, e), demonstrating that effects of GA_9_-methyl ester on spore germination are not dependent on the presence of *Pp*DELLAs. Futhermore, the *Ppdellaab* double mutant is similarly sensitive to spore germination inhibition by ABA as the wild-type, demonstrating that the inhibitory effect of ABA on *Physcomitrium* spore germination is also independent of *Pp*DELLAs (Fig. 4f).

### Loss of *Pp*DELLA function does not affect responses to oxidative stress and ABA

To examine whether *Pp*DELLA proteins have vegetative roles similar to the functions of *Marchantia Mp*DELLA and *Arabidopsis At*DELLAs, we analysed the effects of oxidative stress and the ‘stress hormone’ ABA on *Physcomitrium* vegetative tissue (Supplemental Fig. 5). No differences in response to the oxidative stress-inducing methyl viologen were seen between wild-type and *Ppdellaab* vegetative growth (Supplemental Fig. 5a). Both wild type and *Ppdellaab* vegetative tissue is similarly desiccation-sensitive, and this sensitivity can be rescued in each genotype by pretreatment with 10μM ABA (Supplemental Fig. 5b). This suggests that the vegetative roles of DELLA proteins seen in *Marchantia* and flowering plants are not conserved in the moss *Physcomitrium*.

### *Pp*DELLA proteins promote reproductive development

We observed that the *Ppdellaab* mutant consistently develops fewer sporophytes than wild type plants. Frequent monitoring of developmental processes after watering showed that *Ppdellaab* kept on developing reproductive organs, which usually happens when no fertilisation takes place (Meyberg *et al.*, 2020) but was largely unable to develop sporophytes (Fig. 5a-c).

**Figure 5.**
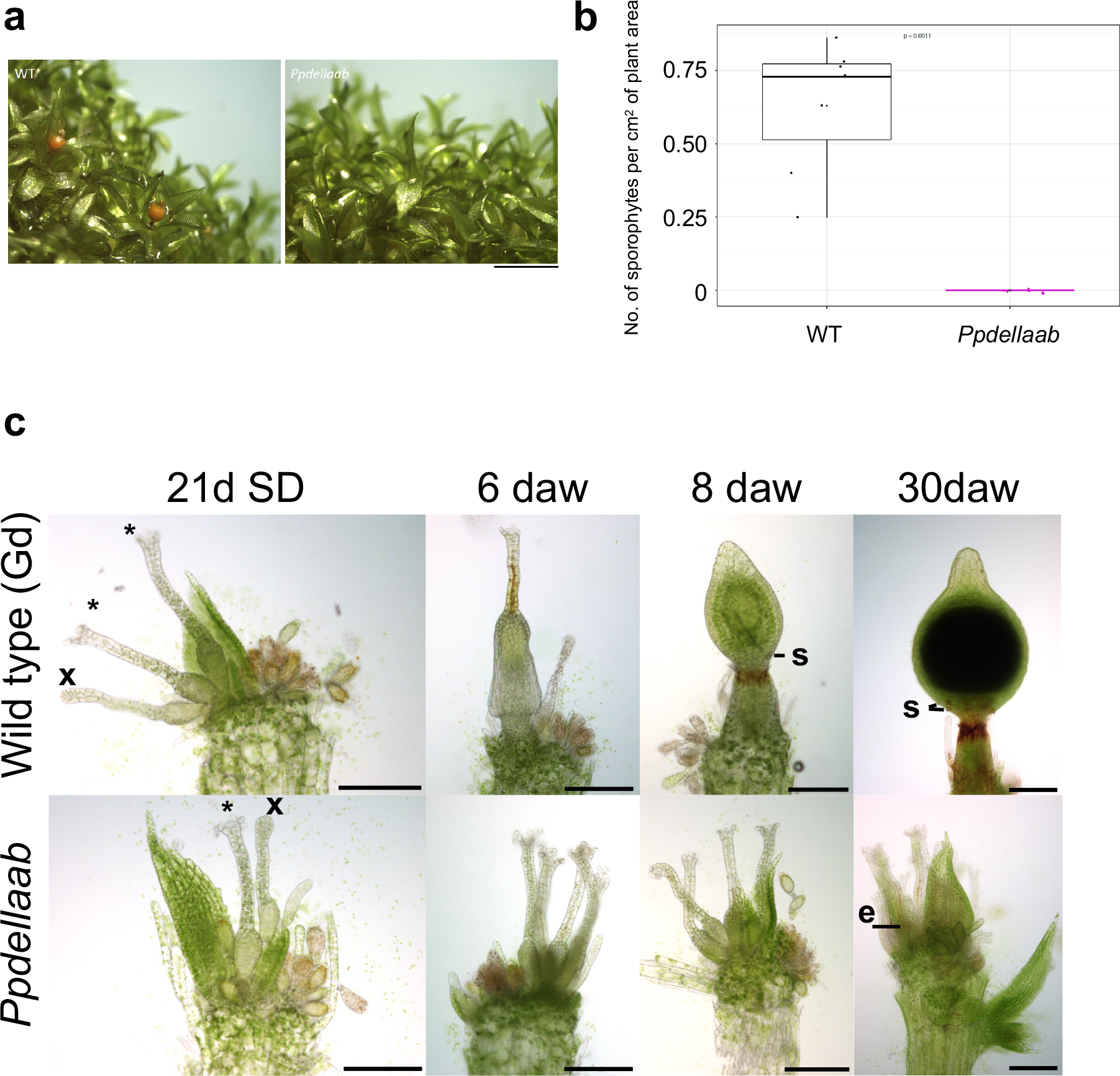
*Ppdellaab* mutants show defective sporophyte development. (A) *Ppdellaab* mutants (right) make fewer sporophytes than wild type (WT, left) plants under conditions that induce sex organ development, fertilisation and sporophyte development (7 weeks at 15°C with 8h light). Scale bar, 2mm. Representative of 6 biological repeats across 2 laboratories. (B) WT plants have higher sporophyte density than *Ppdellaab* plants. The number of sporophytes per cm^2^ plant area is shown. WT and *Ppdellaab* show a statistically significant difference (p=0.0011) in sporophyte density (Mann-Whitney U test; n=7-8). Black asterisks indicate the mean. Representative of 3 biological repeats from 2 independent laboratories. (C) Mature gametangia and sporophyte development of wildtype Gransden (Gd) and the mutant *Ppdellaab*. 21d short day (SD): Gd (wildtype) and *Ppdellaab* (mutant) both show similar amounts of immature (x, closed tip cells) and mature archegonia (*, open tips) and antheridia (swollen tip cell, yellowish color) as well as some older antheridia (brownish color). 6 days after watering (daw): Gd apices show embryos in stage E2 (no stomata present, (Fernandez-Pozo *et al.*, 2020)), *Ppdellaab* apices show several mature archegonia and mature/old antheridia. 8daw: Gd sporophyte development progressed and most sporophytes are in the ES1 stage showing developing stomata (s) and brownish color at the seta. *Ppdellaab* apices show old as well as some new developing gametangia. 30daw: Gd sporophytes are nearly mature, two rows of stomata (s) are present and the seta is dark brown. *Ppdellaab* apices show no sporophytes but multiple old and young gametangia with archegonia possessing darkened and shrunken egg cells (e). Bar: 200μm.

Moreover, *PpDELLAa* and *PpDELLAb* both show enriched expression in developing sporophytes, whereas the expression peak of *PpDELLAa* is in the developing embryo, and the expression peak of *PpDELLAb* is in the early sporophytic stages (Supplemental Fig. 4b, c). To examine the cause of this sporophyte-formation defect, we examined the ability of the *Ppdellaab* mutant to develop both male (antheridia) and female (archegonia) reproductive organs. The *Ppdellaab* mutant develops morphologically normal antheridia and archegonia similarly to wild type plants (Supplemental Fig. 6). However, crossing analysis showed that the *Ppdellaab* mutant likely has a male fertility defect, as *Ppdellaab* mutants can develop sporophytes when crossed with the male-fertile *Physcomitrium* Reute (Re)-mCherry marker strain (Perroud *et al.*, 2019) (Supplemental Fig. 7a). Interestingly, the stomata in crossed plants show aberrant distribution (Supplemental Fig. 7a). In addition, the male infertile strain *Ppccdc39* (Hiss *et al.*, 2017; Meyberg *et al.*, 2020) could not be fertilized by *Ppdellaab*, supporting the hypothesis that *Ppdellaab* has a fertility defect (Supplemental Fig. 7b). This ties in with detection of *PpDELLA* expression in antheridia (Supplemental Fig. 4d). Thus, a positive role for DELLA proteins in male reproductive development is present in the moss *Physcomitrium*, as in flowering plants.

### *Pp*DELLAs interact with *Physcomitrium* hybrid light receptors

Given that the function of *Pp*DELLAs in spore germination and reproductive development cannot be linked to regulation by diterpene/gibberellin signalling, we sought to identify protein interaction partners of *Pp*DELLAs that could represent alternative regulators of their activity. We carried out immunoprecipitation (IP) of *Pp*DELLAa-GFP protein from induced *pHSP*::*Pp*DELLAa-GFP moss tissue using an anti-GFP antibody, followed by mass spectrometry (MS). The proteins obtained by IP-MS from induced *pHSP*::*Pp*DELLAa-GFP were compared to those obtained from uninduced *pHSP*::*Pp*DELLAa-GFP tissue and induced *pHSP*::GFP tissue, to define proteins pulled down specifically in the presence of *Pp*DELLAa (Fig. 6a; Supplemental Table 1). From the 408 proteins specifically immunoprecipitated in the presence of *Pp*DELLAa, 8 proteins came under the Gene Ontology (GO) ‘Biological Function’ category of “chromophore-protein linkage” (GO:0018298) (Fig. 6b). Of these 8 proteins, 3 were putative photoreceptors, namely PHOTOTROPIN A2 (*Pp*PHOTA2), PHOTOTROPIN B1 (*Pp*PHOTB1) and PHYTOCHROME 5B (*Pp*PHY5B).

**Figure 6.**
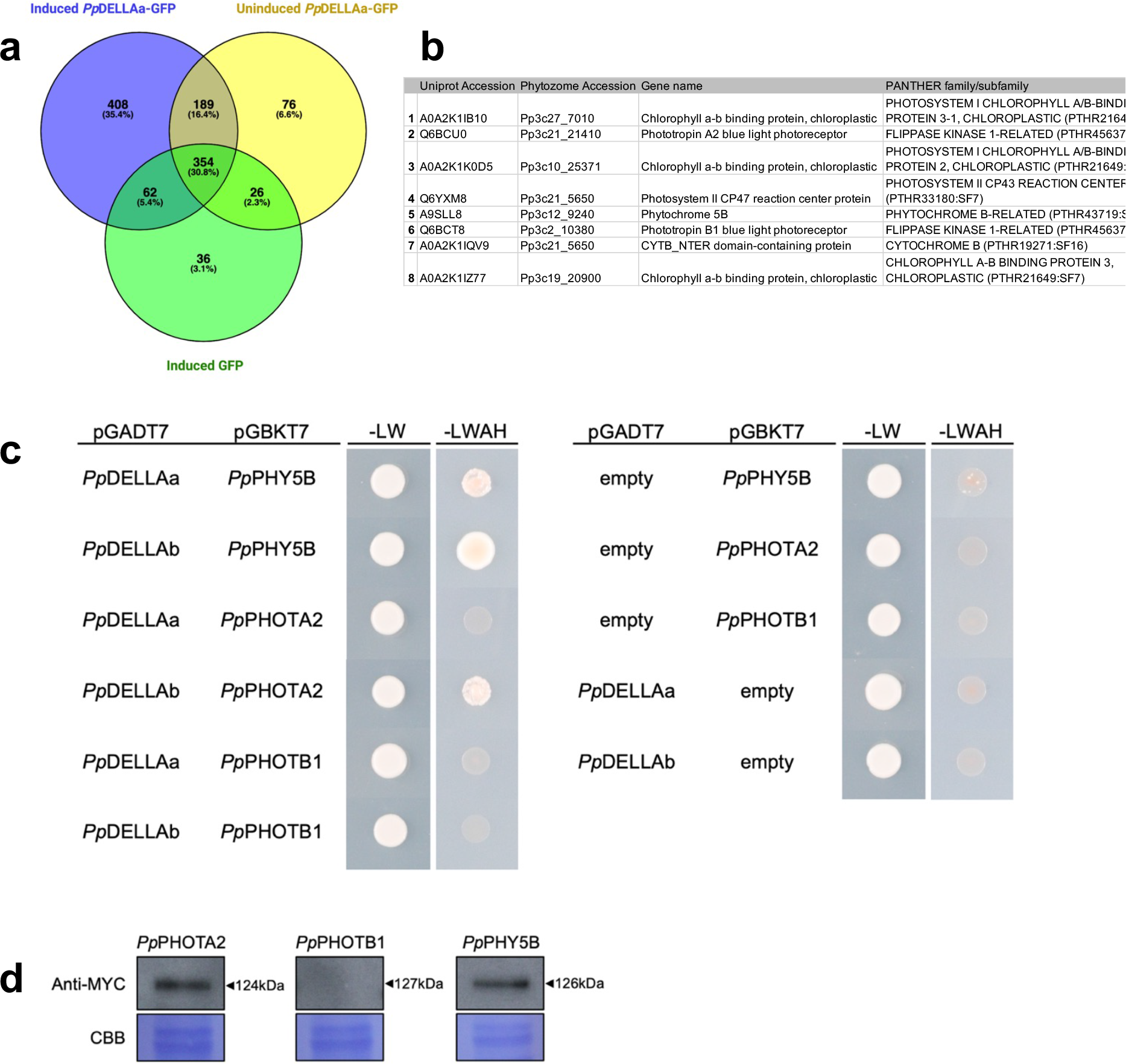
*Pp*DELLA proteins interact with putative light receptors. (A) Common (743) and unique (408) proteins in anti-GFP immunoprecipitations from samples expressing induced *Pp*DELLAa-GFP (blue), uninduced *Pp*DELLAa-GFP (yellow) or induced GFP (green). Venn diagram created with Venny 2.1 (Oliveros, 2007-2015) and edited with BioRender.com. (B) Proteins immunoprecipitated with GO biological function ‘chromophore-protein linkage’ (GO:0018298) from *Pp*DELLAa-expressing plants include three photoreceptors: PHYTOCHROME5B (*Pp*PHY5B), PHOTOTROPINA2 (*Pp*PHOTA2) and PHOTOTROPINB1 (*Pp*PHOTB1). (C) Yeast two-hybrid assay between *Pp*DELLAs fused to the GAL4 activation domain (AD) in pGADT7 and the photoreceptors: *Pp*PHOTA2, *Pp*PHOTB1 and *Pp*PHY5B, fused to the GAL4 DNA-binding (DBD) domain in pGBKT7. *Pp*DELLAa interacted with *Pp*PHY5B; and *Pp*DELLAb interacted with both *Pp*PHY5B and *Pp*PHOTA2. No interaction between *Pp*PHOTB1 was seen with either *Pp*DELLAa or *Pp*DELLAb in this system. (D) Anti-MYC western blot showing that MYC-tagged *Pp*PHOTA2 (124kDa) and *Pp*PHY5B (126kDa) are expressed in yeast but *Pp*PHOTB1 (127kDa) is not. CBB, Coomassie brilliant blue staining.

To confirm the association between *Pp*DELLAs and *Pp*PHOTA2, *Pp*PHOTB1 and *Pp*PHY5B, their interactions were tested in the yeast two-hybrid system (Fig. 6c). *Pp*PHY5B was able to interact with both *Pp*DELLAa and *Pp*DELLAb in yeast, while *Pp*PHOTA2 interacted with *Pp*DELLAb only (Fig. 6c). No interaction was detected between *Pp*PHOTB1 and *Pp*DELLAs in the yeast system, but this is likely due to its lack of detectable expression in yeast compared to *Pp*PHY5B and *Pp*PHOTA2 (Fig. 6d). The interactions between *Pp*DELLAb and *Pp*PHY5B/PpPHOTA2 were not changed in the presence of red, far-red or blue light (Supplemental Fig. 8a). This suggests that *Pp*DELLAs can interact with certain moss photoreceptor proteins independently of a particular wavelength of light.

Since flowering plant phytochromes function as temperature sensors as well as red/far-red light receptors (Jung *et al.*, 2016; Legris *et al.*, 2016) and since moss spore germination is reversibly inhibited by a far-red light pulse or a temperature of 35°C (Vesty *et al.*, 2016), we tested the sensitivity of *Ppdellaab* mutant spores to transient 35°C exposure. *Ppdellaab* mutant spore germination was reversibly inhibited by incubation at 35°C similarly to wild type (Supplemental Fig. 8b). To investigate if *Pp*DELLAs were involved in photoreceptor-mediated regulation of development, we examined the responses of *Ppdellaab* mutant spores to red and blue light. Both wild-type and *Ppdellaab* spores germinated faster under red light than white light and germination of both genotypes was inhibited under blue light (Supplemental Fig. 9a). Similarly, no differences in the vegetative growth of wild-type and *Ppdellaab* mutant plants were observed under red, far-red or blue light (Supplemental Fig. 9b). Taken together, these data suggest that *Pp*DELLA-photoreceptor interactions are not required for spore germination or vegetative growth and that *Pp*DELLAs do not mediate photoreceptor action via physical interaction or any other mechanism.

### *Pp*DELLAs likely function as transcriptional regulators of metabolism

To discover putative transcriptional targets of *Pp*DELLA proteins, we compared the gametophore transcriptomes of wild-type and *Ppdellaab* mutant plants. 782 genes were downregulated in the *Ppdellaab* mutant compared to wild type (Fig. 7a; Supplemental Table 2). 907 genes were upregulated in the *Ppdellaab* mutant (Fig. 7b; Supplemental Table 3). When examined for GO term enrichment of biological processes, genes downregulated in the *Ppdellaab* mutant (therefore likely to be *Pp*DELLA-induced, either directly or indirectly) were largely chloroplast- or photosynthesis-related (Fig. 7c) with some primary metabolic functions (terpenoid/isoprenoid (including carotenoid- and pigment-) biosynthesis and metabolism) and some functions in responses to light and hormones (Fig. 7d). Genes upregulated in the *Ppdellaab* mutant, therefore likely to be directly or indirectly DELLA-repressed, fell largely into the GO biological process category of primary metabolism in addition to secondary metabolism and stress response (Fig. 7c). Metabolic genes included those involved in phenylpropanoid biosynthesis and cell wall metabolism (Fig. 7e).

**Figure 7.**
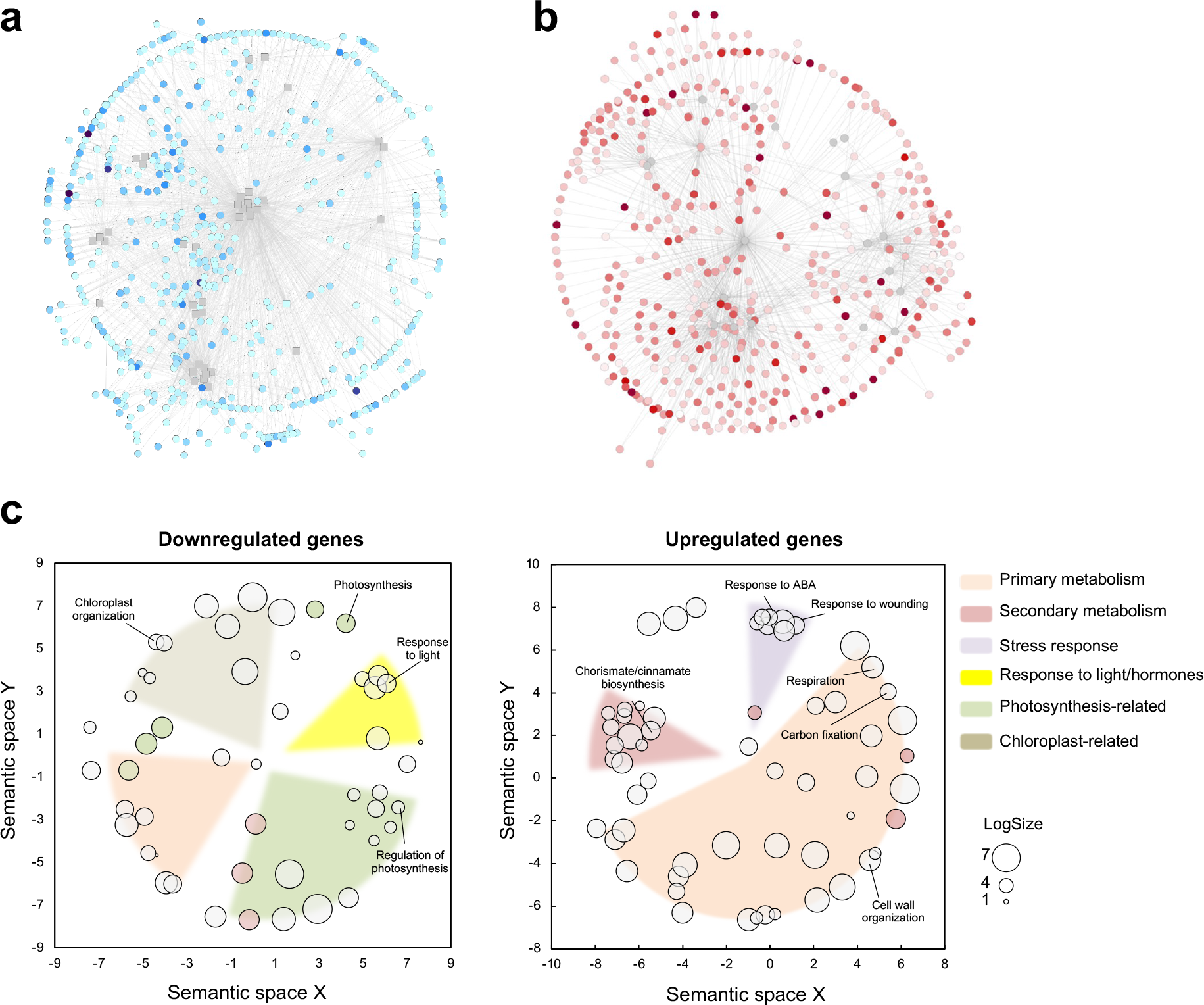
GO term enrichment (Biological Process) for differentially expressed genes in the *Ppdellaab* mutant includes photosynthetic, metabolic and cell wall functions. (A) Genes downregulated in the *Ppdellaab* mutant. Colour intensity represents log fold-change. The network represents regulatory interactions between differentially expressed genes, predicted using PlantRegMap and visualized with Cytoscape. (B) Genes upregulated in the *Ppdellaab* mutant. Colour intensity represents log fold-change. The network represents regulatory interactions between differentially expressed genes, predicted using PlantRegMap and visualized with Cytoscape. (C) Gene Ontology (GO) categories for Biological Process enriched among genes misregulated in *Ppdellaab*. GO term enrichment was calculated using PlantRegMap, and represented with ReviGO, based on the semantic distances between GO terms. *Pp*DELLA-induced biological process GO terms (from genes downregulated in the *Ppdellaab* mutant) include genes involved in photosynthesis and chloroplast function along with primary metabolism and responses to light and hormones. *Pp*DELLA-induced molecular function GO terms (from genes downregulated in the *Ppdellaab* mutant) are largely involved in metabolism (primary and secondary) with some genes involved in stress responses. LogSize represents the Log_10_(Number of annotations for GO Term ID in selected species in the EBI GOA database).

To understand more about how *Pp*DELLAs might carry out their metabolic functions, analysis was carried out using PlantRegMap to predict possible *Pp*DELLA-interacting transcription factors (DELLA-TFs) by identifying enriched transcription factor binding sites in the promoters of *Ppdella* up- and down-regulated genes. Following the assumption that *cis* elements for putative DELLA-TFs would be enriched in the promoters of *Pp*DELLA targets, 51 putative transcription factors were identified (Fig. 8a, b; Supplemental Table 4). Amongst the 21 transcription factors predicted to bind to the promoters of genes downregulated in *Ppdellaab* (therefore induced by *Pp*DELLA activity), binding sites for certain C2H2, DOF, LBD, bZIP, ERF, C3H, bHLH, SBP, WRKY, G2-like and Trihelix transcription factors were enriched (Fig. 8a). Amongst the 41 transcription factors predicted to bind to the promoters of *Ppdellaab*-upregulated genes (genes repressed by *Pp*DELLA activity), binding sites for specific transcription factors from the LBD, ERF, bZIP, WRKY, bHLH, C2H2, CAMTA, MYB, TCP, NAC, HSF, AP2 and BES1 families were enriched (Fig. 8b). There are 11 DELLA-TFs that are predicted to bind to both *Pp*DELLA-induced and DELLA-repressed gene promoters (Supplemental Table 4). TFs belonging to each of these families have been previously shown to act as DELLA interactors in *Arabidopsis* (Marin-de la Rosa *et al.*, 2014; Lantzouni *et al.*, 2020). Moreover, one of the transcription factor-encoding genes identified as a putative *Pp*DELLAa-interactor using PlantRegMap is the MYB domain transcription factor *Pp3c3_17580V3.1*. *Pp*3c3_17580V3.1 was also detected as a *Pp*DELLAa-interacting protein in the IP-MS experiment (Supplemental Table 1). These data suggest that as in flowering plants (Hernandez-Garcia *et al.*, 2019; Hernandez-Garcia *et al.*, 2021b; Phokas & Coates, 2021), *Pp*DELLAs act as ‘hubs’ for interaction with transcription factors. There is no overlap between the putative *Pp*DELLA-interacting transcription factor genes and the 88 DEGs in moss tissue treated with GA_9_-methyl ester (Perroud *et al.*, 2019) (Fig.7c). In total, 29 out of the 88 GA_9_-methyl ester DEGs (33%) are present amongst the 1689 *Ppdellaab* DEGs (Fig. 7c). This supports the idea that *Pp*DELLA and diterpenes act via independent transcriptional mechanisms in *Physcomitrium* although they impinge on some of the same developmental processes.

**Figure 8.**
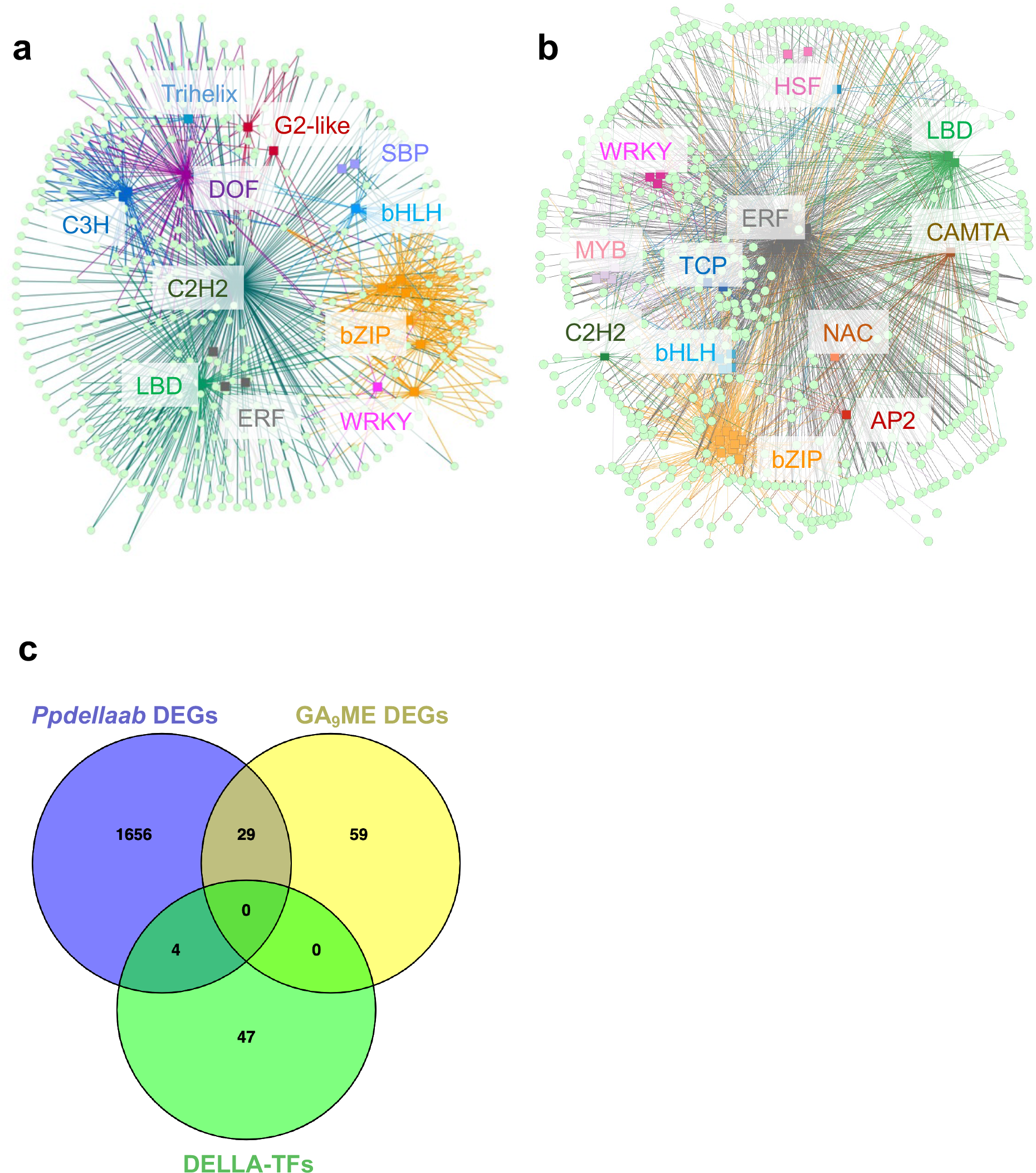
*Pp*DELLAs may act as transcriptional regulators. (A) Classes of transcription factors (TFs) most likely to be responsible for the regulation of *Pp*DELLA-induced genes (the TFs have putative binding sites in the promoters of genes downregulated in the *Ppdellaab* mutant), according to a TF-enrichment analysis performed with PlantRegMap, visualised using Cytoscape. (B) Classes of transcription factors (TFs) most likely to be responsible for the regulation of *Pp*DELLA-repressed genes (the TFs have putative binding sites in the promoters of genes upregulated in the *Ppdellaab* mutant), according to a TF-enrichment analysis performed with PlantRegMap, visualised using Cytoscape. (C) There is little overlap between the genes differentially expressed between wild type and *Ppdellaab* mutants, genes induced by GA_9_-methyl ester (Perroud et al 2018) and the transcription factors (DELLA-TFs) predicted to bind to the promoters of genes differentially expressed in the *Ppdellaab* mutant (shown in panels A and B). Venn diagram produced using Venny 2.1 (Oliveros, 2007-2015).

## Discussion

### *Physcomitrium* DELLAs have different biological functions to DELLAs in angiosperms and liverworts

Our data show that *Pp*DELLAs do not show diterpene-dependent regulation or interactions. Although *Pp*DELLAs restrain germination, this is independent of the germination-promoting functions of diterpenes. Moreover, *Ppdellaab*mutant spore germination is inhibited by ABA. This is in contrast to how the *Arabidopsis* DELLAs *At*RGL2, *At*RGA and *At*GAI affect seed germination, where loss of these three DELLAs in the absence of gibberellin biosynthesis (*Atga1-3* mutant background) renders seed coat rupture (potentially analogous to spore coat rupture in moss) insensitive to ABA (Piskurewicz *et al.*, 2009).

Roles for bioactive diterpenes related to gibberellin in spore germination and vegetative growth have been proposed in *Physcomitrium* (Hayashi *et al.*, 2010; Vesty *et al.*, 2016). *Physcomitrium* plants that make no diterpenes as they are mutant for the bifunctional *COPALYL DIPHOSPHATE SYNTHASE/KAURENE SYNTHASE* (*PpCPS/KS*) gene show slower spore germination (Vesty *et al.*, 2016) and defects in differentiation from chloronema (a highly photosynthetic cell type) to caulonema (a faster-elongating, less photosynthetic cell type, which gives rise to the leafy gametophore that is required for sexual reproduction) (Harrison *et al.*, 2009) (Hayashi *et al.*, 2010; Miyazaki *et al.*, 2014; Hiss *et al.*, 2017). Application of *ent*-kaurene or GA_9_-methyl ester can rescue these phenotypes (Hayashi *et al.*, 2010; Miyazaki *et al.*, 2014; Vesty *et al.*, 2016) and speed up wild-type spore germination (Vesty *et al.*, 2016) whereas flowering plant gibberellins do not replicate these effects (Hayashi *et al.*, 2010; Vesty *et al.*, 2016). However, GA_9_-methyl ester has no effect on the inhibition of spore germination by far-red light (Vesty *et al.*, 2016), despite a proposed role for diterpenes (*ent*-kaurene and *ent*-kaurenoic acid) in protonemal (vegetative tissue) responses to blue light in *Physcomitrium* (Miyazaki *et al.*, 2014). Taken together, these data suggest that the signalling networks for DELLAs, diterpenes and light in *Physcomitrium* are wired differently from those in land plants and may be largely independent from one another.

In contrast to the situation in angiosperms (Thomas *et al.*, 2016; Vera-Sirera *et al.*, 2016) and *Marchantia* (Hernandez-Garcia *et al.*, 2021b), *Pp*DELLAs do not restrain vegetative growth or protect against oxidative stress, desiccation or salt stress (Yasumura *et al.*, 2007) in *Physcomitrium*. *Pp*DELLA functions could reflect their divergent N-terminal sequences and protein regulation. Moreover, *Physcomitrium* lacks the apical notch structure seen in *Marchantia* vegetative tissue.

*Pp*DELLAs do play a role in reproductive development as *Ppdellaab* mutants are impaired in sporophyte formation. DELLAs are required for male reproductive development in *Arabidopsis* and rice via effects on anther- and pollen development (Plackett *et al.*, 2014; Jin *et al.*, 2022). In angiosperms, gibberellin signalling controls pollen development via the GAMYB transcription factor (Murray *et al.*, 2003). Gibberellin-related substances (including GA_9_-methyl ester) have been implicated as ‘antheridiogens’ in reproductive development in ferns (Yamane, 1998). GAMYB transcription factor activity regulates spore formation in the lycophyte *Selaginella* and in *Physcomitrium*, suggesting that GAMYB may be part of an ancient “reproductive” module that later became linked to gibberellin signalling (Aya *et al.*, 2011).

In *Marchantia*, overexpression of *Mp*DELLA eliminates the formation of reproductive structures after induction by far-red light (Hernandez-Garcia *et al.*, 2021b). In *P. patens*, we found that sporophyte development in *Ppdellaab* mutants is nearly inhibited (Figure 4) but in contrast to *Marchantia*, antheridia (male) and archegonia (female) development takes place similarly to wild type (Supplemental Fig. 5). When crossed with a fertile moss marker strain, *Ppdellaab* mutants are able to develop sporophytes similar to wildtype level, but the crossing attempt with a male sterile moss line failed, showing that *Ppdellaab* mutants have a male sterility phenotype. Given that mosses possess biflagellated spermatozoids as opposed to pollen, and that *Ppdellaab* mutant antheridia appear normal, it is likely that the effect of *Pp*DELLA on male fertility is mechanistically distinct from the effect of DELLA function on male fertility in flowering plants and is a result of convergent evolution from rewiring of ancient protein modules.

*Pp*DELLAs show an interaction with photoreceptors (*Pp*PHOTA2, *Pp*PHOTB1, *Pp*PHY5B) that is independent of light wavelength. *PpPHOTA2* and *PpPHOTB1* are two of the four PHOTOTROPIN genes in *P. patens*, which encode blue light receptors that also play a role in chloroplast responses to red light (Kasahara *et al.*, 2004) by interaction with the phytochrome *Pp*PHY4 (Jaedicke *et al.*, 2012). *PpPHY5B* is one of seven *P. patens PHYTOCHROME* genes and is a moss phytochrome family member that has not been well characterised to date (Mittmann *et al.*, 2009; Possart & Hiltbrunner, 2013; Trogu *et al.*, 2021).

A direct interaction of DELLA proteins with light receptors has not previously been detected. In *Marchantia* and flowering plants DELLAs and red/far-red light signalling are linked via PIFs, which interact with both DELLAs and phytochromes (de Lucas *et al.*, 2008; Feng *et al.*, 2008; Hernandez-Garcia *et al.*, 2021b). The GRAS domain of DELLA mediates its interaction with PIF (de Lucas *et al.*, 2008). *Physcomitrium* also possesses PIFs, but they are not degraded by red light when expressed in *Arabidopsis* (Xu & Hiltbrunner, 2017).

In a potential link between red/far-red and blue light signalling in moss, a direct interaction is seen between a *Physcomitrium* PHYTOCHROME (*Pp*PHY4) and PHOTOTROPINs (*Pp*PHOTs A1, A2 and B1) specifically at the plasma membrane (Jaedicke *et al.*, 2012). It is tempting to speculate that *Pp*DELLAs could be co-ordinating *Pp*PHY and *Pp*PHOT functions in the nucleus or that *Pp*PHY and *Pp*PHOT activity could regulate an aspect of *Pp*DELLA function (although this seems less likely to be a direct effect given the lack of light-dependent phenotypes in *Ppdellaab* mutants). It is also interesting to note that blue light protonemal avoidance is disrupted in the *Ppcps*/*ks* mutant (Miyazaki *et al.*, 2014) implying a role for diterpene signalling in moss light responses, independently of DELLA functions. Taken together, our data suggest that the wiring of moss light-, DELLA- and diterpene signalling pathways is divergent from other land plants.

Similarly to the situation in other land plants, *Physcomitrium* DELLAs likely function as transcriptional regulators. However, the transcriptional targets of *Pp*DELLAs are largely distinct from those involved in diterpene-dependent pathways and there is no overlap between GA_9_ME-induced genes and putative *Pp*DELLA-interacting TFs and their putative target genes. This further supports our conclusions that although *Pp*DELLAs and moss bioactive diterpenes may regulate some of the same biological processes ((Vesty *et al.*, 2016); this work), they do so via independent mechanisms.

It remains to be seen whether *Physcomitrium* DELLA proteins have different functions from those in other plants because of their divergent N-terminus, because of the identity and functions of their putative interaction partners, or because moss DELLA proteins more broadly have acquired divergent functions due to unique characteristics of the moss lifestyle and life-cycle. Examples of unique characteristics include the leafy rather than thalloid gametophyte, robust and slow-developing sporophytes and unique mechanisms of sporophyte development, specific microclimates and shading environments in particular ecological niches (Glime, 2017b; Glime, 2017c; Glime, 2017a).

## Conclusions

In this paper, we have demonstrated that *Physcomitrium* DELLA proteins have divergent N-termini compared to other land plants. In concordance with this, *Pp*DELLA protein stability is not affected by diterpenes or gibberellin and *Pp*DELLA proteins do not interact with gibberellin receptor-like proteins from *Phycomitrium* or *Arabidopsis*. Moreover, the positive effects of diterpenes on spore germination are independent of the restraining effect of *Pp*DELLAs on the same process. *Pp*DELLAs show a so far unique interaction with red- and blue light receptors, however no light-dependent phenotypes were detected in the *Ppdellaab* mutant. *Pp*DELLAs promote male fertility, which is similar to land plant DELLAs, although the effects are probably via unrelated mechanisms. *Pp*DELLAs likely function via interaction with multiple transcription factors as seen in other land plants: we have identified areas for future research. We conclude that the wiring of DELLA-, light- and diterpene signalling networks in *Physcomitrium* is substantially different to that in other land plants.

## Acknowledgements

AP was supported by the UK Biotechnology and Biological Sciences Research Council (BBSRC) doctoral training grant BB/M01116X/1. EFV was funded by a UK Natural Environment Research Council (NERC) doctoral training scholarship. SAR and RM were supported by grant RE 1697/15-1 from the DFG (German Research Foundation).

MAB, AB-M and J H-G were supported by grant PID2019-110717GB-I00 funded by Spanish MCIN/ AEI/ 10.13039/501100011033.

We thank Yuki Yasumura, Eric Belfield and Nicholas Harberd (University of Oxford) for contributing *Pp*DELLA and *Pp*GLP yeast two-hybrid constructs and for *Ppdella* mutants. We thank Professor Peter Hedden, Rothamsted Research (UK) for generous provision of diterpene supplies. We thank Amber Spiteri and Alec Ballentyne for initiating the yeast two-hybrid system for *Pp*DELLAs and *Pp*GLPs. We thank Amy Whitbread, Sarah Needs, Sue Bradshaw and Dan Holloway (University of Birmingham) for early maintenance of *Ppdella* moss sporophyte stocks and for preliminary *Ppdella* germination phenotyping. We thank the University of Birmingham Advanced Light Microscopy (BALM) facility for training and technical assistance. We thank Jon Hughes for advice with the light- and phytochrome/phototropin experiments. We thank Evelyn Vollmeister (University of Marburg) for comments on the manuscript.

## Author contributions

AP, RM, MAB, SAR. designed research. AP, RM, AB-M, JH-G, PW, EFV performed research. All authors analysed data. AP, RM, AB-M, JH-G visualised data. MAB, SAR, JCC supervised parts of the project. AP, RM, MAB, JCC wrote the manuscript. All authors reviewed the manuscript.

## Data availability

RNAseq data are deposited at NCBI BioProject database under accession number PRJNA695244 (“DELLA-dependent transcriptomes in different plant species”).

## Supplemental material

**Supplemental methods**

These describe further details of published methods cited in the Materials and Methods section.

**Supplemental tables 1-5**

**Supplemental table 1:** list of proteins identified as interacting with *Pp*DELLAs. 408 proteins were identified. Their unique (PANTHER) identifier assigned by the FASTA database used, description, molecular weight (MW), number of distinct peptides detected and protein FDR confidence (1% cut-off) are shown.

**Supplemental Table 2:** genes downregulated in the *Ppdellaab* compared to wild type (*Pp*DELLA-induced genes) (p<0.01).

**Supplemental Table 3:** genes upregulated in the *Ppdellaab* compared to wild type (*Pp*DELLA-repressed genes) (p<0.01).

**Supplemental Table 4:** transcription factor binding sites enriched in the promoters of *Pp*DELLA-induced genes and *Pp*DELLA-repressed genes identified by PlantRegMap. Both the TF target genes and the identified putative TFs are listed.

**Supplemental Table 5:** Primers used in this paper.

**Supplemental Figures 1-9**

**Supplemental Figure 1:** full length DELLA protein sequence alignment.

**Supplemental Figure 2.** Generation of *pHSP::PpDELLA-GFP* and *pHSP::GFP* transgenic lines.

**Supplemental Figure 3.** Induction of *Pp*DELLA-GFP and GFP protein expression by heat shock is sustained for at least 26 hours.

**Supplemental Figure 4.** *PpDELLA*s are strongly expressed in dry spores and developing sporophytes.

**Supplemental Figure 5**. *Ppdellaab* mutants do not show altered responses to salt, oxidative or desiccation stress compared to wild type (WT).

**Supplemental Figure 6.** *Ppdellaab* mutants can develop antheridia and archegonia.

**Supplemental Figure 7.** *Ppdellaab* mutants develop sporophytes when fertilized by a Reute (Re)-mCherry wild type strain but not when crossed with the male sterile mutant *Ppccd39*.

**Supplemental Figure 8.** *Pp*DELLA proteins and show no differences in interaction with light receptors in yeast in response to light wavelength and *Ppdellaab* mutant spores show normal thermoinhibition.

**Supplemental Figure 9.** *Ppdellaab* mutants respond to different light wavelengths similarly to wild type (WT) during spore germination and vegetative growth.

