## Supporting information for "DELLA proteins regulate spore germination and reproductive development in *Physcomitrium patens*"

**The following supporting information is available for this article:**

### Supplemental Figures 1-9.

### Figure S1.

```
AtRGA1       1 -------MKRDHHQFQGRLSNH---GTS--SSSSSISKDKMMMVKKEEDGGGNM-D----
AtGAI1       1 ---------------------M---KRD--HHHHHHQDKKTMMMNEED--DGNG-M----
AtRGL2       1 -------MKRGYGETWDPPPKPLPASRS--GEGPSMADKK---KADDDNNNSNM-D----
MtDELLA      1 -----------------------------------------MWREE-KETNGGG-M----
SlDELLA      1 -------MKRDRDRDREREKRA---FSN--GA--VSSGKSKIWEEDEEEKPDAG-M----
HvSLN1       1 -------MKREYQDGGGSGGG----G----DEMGSSRDKMMVSSS--EAGEGEE-V----
TaRHT-B1     1 -------MKREYQDAGGSGGG----G----GGMGSSEDKMMVSGSA-AAGEGEE-V----
ZmD8         1 -------MKREYQDAGGSG-----------GDMGSSKDKMMAAAAGAGEQEEED-V----
OsSLR1       1 -------MKREYQEAGGSSGG----GSS--ADMGSCKDKVMAGAAG----EEED-V----
AtrDELLA1    1 -------MKRAQG--------D---SSS--GAYRASHGKSK-----MQEPQDAG-V----
AtrDELLA2    1 ------------------------MDYPFKTF-----------PSTPSPPSKPE-I----
PtRGA1       1 ---------------------MDPMERAAKVLGSSPGHKNMMGCSSSGVKVEPE-I----
CrDELLAa     1 ------------------------------MF-QSPSDSLLPQNQTMGLGD-AD-I----
CrDELLAb     1 -------MRPDSRLSQAS---V----TQAEML-CCPSDSTFSQRQSMGLGREAD-I----
SkDELLA      1 MESMLQAVYEGARSAAAR---RTTMEANESMAGGNGGGG--GGHEQWKSTTTTE-V----
SmDELLAa     1 ----------------------------------------------------MG-M----
EsDELLA      1 -----------------------------MAY------QYHPGNSRHDATGGTL-V----
TauDELLA     1 -----------------------------MAY------QYYPGNSRREATGGAI-V----
HcDELLA      1 -----------------------------MAY------QYYPGSSRHEATGGAI-V----
SfaDELLA     1 -----------------------------MAY------QYYPGSSRHEATGGAI-V----
PpDELLAa     1 -----------------------------MAY------QYSPGGSRWKPTGGTL-V----
PpDELLAb     1 -----------------------------MAY------QYYPGSTRYEATGGAL-V----
SfDELLAa     1 ----------------------------MMAFRNLSSRSWDRGGSGGGGGGGAM-V----
MpDELLA      1 MDSSADY----ARRV--R---ARPSSSSASDLTGVTSPQYRH------HSGSVG-V----
AaDELLA      1 MSTSAQL-KDSSRAALYHA------GNEGQGMVDRHRDG---AMLEGPSPGGAG-I----
ApDELLA      1 -------------------------------------------MLEGPSPGGAG-I----

AtRGA1      44 DEL-LAVLGYKV--RSSE-MAEVALKLE-QLETMMSN---------VQEDGLS-------
AtGAI1      28 DEL-LAVLGYKV--RSSE-MADVAQKLE-QLEVMMSN---------VQEDDLS-------
AtRGL2      44 DEL-LAVLGYKV--RSSE-MAEVAQKLE-QLEMVLSN---------DD--VGS-------
MtDELLA     14 DEL-LAALGYKV--RSSD-MADVAQKLE-QLEMVMGS---------AQEEGIN-------
SlDELLA     42 DEL-LAVLGYKV--KSSD-MADVAQKLE-QLEMAMGT---------TMEDGIT-------
HvSLN1      39 DEL-LAALGYKV--RASD-MADVAQKLE-QLEMAMGMGG-----PAPDDGFAT-------
TaRHT-B1    40 DEL-LAALGYKV--RASD-MADVAQKLE-QLEMAMGMGGVGA-GAAPDDSFAT-------
ZmD8        38 DEL-LAALGYKV--RSSD-MADVAQKLE-QLEMAMGMGGVGGAGATADDGFVS-------
OsSLR1      39 DEL-LAALGYKV--RSSD-MADVAQKLE-QLEMAMGMGGVSAPGA-ADDGFVS-------
AtrDELLA1   31 DEL-LASLGYNV--RASD-MAEVAQKLE-QLEMVMGT---------AQEDGIS-------
AtrDELLA2   21 DGL-LADAGYRI--KASD-LPHVAHRLE-QLETQMIN---------AQPTEMT-------
PtRGA1      35 DGL-LANAGYTV--KASD-LAHVAQRLE-QLESIMGT---------VQDPGIS-------
CrDELLAa    24 ETL-LAGAGYNV--KASD-LALVAQRLE-LLDSLCSS---------HDAGALS-------
CrDELLAb    41 EAL-LADAGYNV--KASD-LALVAQRLE-QLDSLCAS---------QDTGALS-------
SkDELLA     51 DEH-LARVGYNV--RASE-LPHIAQQIE-VLDSLIGA---------APESLLG-------
SmDELLAa     4 DEL-LAHAGYNV--RASD-LTHVAQRIE-ELDSLLGA---------AAPA--D-------
EsDELLA     21 DGQ-LRHVSFMQ-------PSDLVQHLE-QLHSVLGA---------ASQDSASIPAHDTT
TauDELLA    21 DGQ-FRHVNFMQ-------PSDLVQHLE-QLHGVLGA---------AAQE-TGIPAHHTS
HcDELLA     21 DGQ-FRHSNFMQ-------PSDVVQHLE-QLHSVLGT---------AAQD-PGIPVHHTS
SfaDELLA    21 DGQ-FRHSNFMQ-------PSDVVQHLE-QLHSVLGV---------AAHE-SGVPAHHTS
PpDELLAa    21 DGR-LRHDKFTQ-------ASDAVQQLE-ELHTSLGS---------VSQDSLNIPAYYTL
PpDELLAb    21 DGQ-FRHANFMQ-------PSDLVQHLE-QLHSVLGT---------VSQDSPNIPAHHTL
SfDELLAa    28 DEGQLHNRNCNEYTSTVHSSDHHLAMAQRLE-QLETVLTA------AAAQEASIAHSSS-
MpDELLA     41 DQQLLAHTGG---YSNVR----SGDMGLRLEQQLDTVLGV------S-QDG---PISHL-
AaDELLA     46 DEF-LANVGYSV--KGSADLDDVAQKLE-LLENVVGS---------APEGNIL-------
ApDELLA     13 DEF-LANVGYSV--KGSADLDDVAQKLE-LLENVVGS---------APEGNIL-------

AtRGA1      83 HLATDTVHYNPSELYSWLDNMLSELNPPPLPASSNGLDPV---LPSPEIC---------G
AtGAI1      67 QLATETVHYNPAELYTWLDSMLTDLNPPSS------------------------------
AtRGL2      81 TVLNDSVHYNPSDLSNWVESMLSELNNPASSDLDT----T--------------------
MtDELLA     53 HLSSDTVHYDPTDLYSWVQTMLTELNPDSSQINDP----LASLGSSSEIL---------N
SlDELLA     81 HLSTDTVHKNPSDMAGWVQSMLSSISTNFDMCNQENDVLVSGCGSSSSII---------D
HvSLN1      82 HLATDTVHYNPTDLSSWVESMLSELNAPPPPLPPAPPQLNA-STSSTV------------
TaRHT-B1    87 HLATDTVHYNPTDLSSWVESMLSELNAPPPPLPPAP-QLNA-STSSTV------------
ZmD8        86 HLATDTVHYNPSDLSSWVESMLSELNAPPAPLPPATPAPRLASTSSTVTS---------G
OsSLR1      86 HLATDTVHYNPSDLSSWVESMLSELNAPLPPIPPAPPAARHASTSSTVTG---------G
AtrDELLA1   70 HLASETVHYNPSDIATWIESMLTELNFPPNLGAPYAPNPSNNCWTAAELP---------P
AtrDELLA2   60 HLASETVHYNPSDLATWVESMLFELNPSEIPATSG-------------------------
PtRGA1      74 HLASEAVHYNPSDLAGWIESMFGELNPGADMPVPFGDRG---SLID--------------
CrDELLAa    63 YLSSEAVHYNPSDMASWLECMIGELAPSSAPTDICSFQG---VLE-GHFSQQTSGHYGID
CrDELLAb    80 YLSSEAVHYNPSDMAAWLECMIGELGPSSVPGDVGGTQR---PASENPLPPLSSTFYDFG
SkDELLA     90 GVSQDTVHYNPSDLASWVECLLDELGPLPASMATTTTTT---SMVR--------------
SmDELLAa    41 ILAQDTVHYNPSDLVSWIEGMLDELVPQQPTATSSSDME---SVN---------------
EsDELLA     63 DSGPQTNINRPTDLAGWIDCMIEELSSNTAGPMAA-Q-QQRSPLTED-------SLRKND
TauDELLA    62 NSVPQISNRPGNDLAGWIDCMIEELSSNTAGPMAAPQ-QQRSSLTED-------SLHNND
HcDELLA     62 EAGPQISNRSS-NMTGWIDCMIEELSSNTAGPMAVAQ-QQRSSLSED-------SLHNND
SfaDELLA    62 EYGLQISNRSS-NMTGWIDCMIEELSSNTAGPMAVTQ-QQRSSLSED-------SLHNND
PpDELLAa    63 GSSSQAVSNCSTDLAGWIDCMIEELSSNTACPIMAPQ-QQH-GLLEG-------SFLKND
PpDELLAb    63 DAGAQTSNNRTSDLAGWIDGMIDELSFNNAGTMAAPQ-Q--RSLTED-------SLHQNE
SfDELLAa    80 ---DGSMALNSSDLSGWIEGMIEELTAANNVP----A-QRSSPFTAD-------SPYNNN
MpDELLA     83 -SAEAAQHYNPADLAGWIECMICDMQPSSSQAQLHTSQQQQQQHSP-------TPTHS-
AaDELLA     86 QLLNEAMHNNPSEIAAWIETMIKELSGPANVSGAAGVYA------GGVATPGLSSVPGSN
ApDELLA     53 QLLNEAMHNNPSEIAAWIETMIKELSGPANVSGAAGVYA------GGVATPGLSSVPGSN
```

```
AtRGA1      131 ----------------------FPAS----------------------------------
AtGAI1       97 ------------------------NA----------------------------------
AtRGL2      117 -------------------RSCVDRS----------------------------------
MtDELLA     100 --------------NTF-----NDDS----------------------------------
SlDELLA     132 --------------FSQ-NHRTSTIS----------------------------------
HvSLN1      129 --------TGGGGYFDLPPSVDSSSS----------------------------------
TaRHT-B1    133 --------T-GGGYFDLPPSVDSSCS----------------------------------
ZmD8        137 A-------AAGAGYFDLPPAVDSSSS----------------------------------
OsSLR1      137 ---------GGSGFFELPAAADSSSS----------------------------------
AtrDELLA1   121 --------------PPL-PDSNA--A----------------------------------
AtrDELLA2    95 ------------------------------------------------------------
PtRGA1      117 ------------------SSQ-FHKPLQDD---------------------PSLSAMDLA
CrDELLAa    119 DVYGPFGCTRGTDYQLNKPNTFLQDSFPNPQPKQGAL--PSVLLQTPVECVTSIPQLIRD
CrDELLAb    137 NVNSSVPCSSVVKNSFIDQKSSVHSPFVDCPPKQAVPQPALGILDPTAEGLPSISQLIKD
SkDELLA     133 ------------------AES-E------S---------------------SS-------
SmDELLAa     83 ----------------------E------V---------------------G--------
EsDELLA     114 L-E-A--------------TSSRDSSLDTNSS-Q--------------------------
TauDELLA    114 L-E-V--------------SSSRDSSLETGSP-Q--------------------------
HcDELLA     113 L-E-V--------------TSSRDSSLETGSP-Q--------------------------
SfaDELLA    113 L-E-V--------------TSSRDSSLETGSP-Q--------------------------
PpDELLAa    114 H-D-A--------------SSCRDSLLETGSH-R--------------------------
PpDELLAb    113 L-E-A--------------SSSHDSSLDTGSS-R--------------------------
SfDELLAa    125 TVEGS--------------STSLDSSLDTDSPSQ--------------------------
MpDELLA     134 ----S--------------FVSMESSLDSMDS-Q--------------------------
AaDELLA     140 ----------MMGDTNS-PSTSM-RNISASSPMNMALD-PVVSMAESQNCLPYATKLQQG
ApDELLA     107 ----------MMGDTNS-PSTSM-RNISASSPMNMALD-PVVSMAESQNCLPYATKLQQG

AtRGA1      135 --------------DYDLKV------IPGNAIY----------QFP--------------
AtGAI1       99 --------------EYDLKA------IPGDAIL----------NQF--------------
AtRGL2      124 --------------EYDLRA------IPGLSAF----------PKE--------------
MtDELLA     107 --------------EYDLSA------IPGMAAY----------PPQ--------------
SlDELLA     143 --------------DDDLRA------IPGGAVF----------NSD--------------
HvSLN1      147 --------------TYALRP------IISPPVA----------PAD--------------
TaRHT-B1    150 --------------TYALRP------IPSPAVA----------PAD--------------
ZmD8        156 --------------TYALKP------IPSPVAA----------PSA--------------
OsSLR1      154 --------------TYALRP------ISLPVVA----------TAD--------------
AtrDELLA1   130 --------------ESCSSN------LHPPQLF----------DSS--------------
AtrDELLA2    95 ---------------------------------------------F-AGESQSPEIGFFA
PtRGA1      137 LIQEY---------------------------------GLQ----F-NGNQASNPQGFSP
CrDELLAa    177 AIGNQGGASATADRNESRSS------YPGVTLPKRDVGGLHHYKELEDQGSCNQAKGFCA
CrDELLAb    197 AIGHNGGAPAAS---ATLKG------YPGIALKDRTPGGLQQHKIIEDQGSSNQVGAFFP
SkDELLA     140 -VVTN---------------------------------SQH----F----------GFAP
SmDELLAa     86 -VVAS---------------------------------HSQ----I-A-------ASTTP
EsDELLA     131 --------------------------LP----------TLN-YQDT-----PAVRTNFAA
TauDELLA    131 --------------------------LP----------TLL-YRDT-----PAVGTNFAT
HcDELLA     130 --------------------------LL----------TLQ-YRDT-----PAVGTNFAA
SfaDELLA    130 --------------------------LP----------TLQ-YRDA-----SAAGTNFIA
PpDELLAa    131 --------------------------LS----------NVQ-FQDT-----SAARNKSST
PpDELLAb    130 --------------------------LP----------TLH-YQNT-----PAVGNNFLA
SfDELLAa    145 --------------------------VP----------PLH-YQEA-----LLDNGFSST
MpDELLA     149 --------------------------AP-----------LQ------------------P
AaDELLA     187 FQGQP---NMFMDKYDGSTGMVGSTGLHGPHDASVSETVDQQWQPSANQL----SHNYYH
ApDELLA     154 FQGQP---NMFLDKYDGSTGMVGSTGLHGPHDASVSETVDQQWQPSANQL----SHSYYH

AtRGA1      151 ------------------------------------------------------------
AtGAI1      115 ------------------------------------------------------------
AtRGL2      140 ------------------------------------------------------------
MtDELLA     123 ------------------------------------------------------------
SlDELLA     159 ------------------------------------------------------------
HvSLN1      163 ------------------------------------------------------------
TaRHT-B1    166 ------------------------------------------------------------
ZmD8        172 ------------------------------------------------------------
OsSLR1      170 ------------------------------------------------------------
AtrDELLA1   146 ------------------------------------------------------------
AtrDELLA2   109 GNQ---------------------------------------------------------
PtRGA1      159 DSDPSV----------RCNIFS------GLPL----RSGDS---TRHTNFQA--------
CrDELLAa    231 GNSTQPCLISHVSLQKSCSMPSLHQLQQAGHISATQARGSFSFHTQHQTQGQSFSSPAAS
CrDELLAb    248 RSSAGD----------PPQLSNMSTLQQAVPIPSPKMHGNPSLSMQHQMQSQSLFSSVSI
SkDELLA     152 QPQQQQ---------------------------------------------Q--------
SmDELLAa    100 RPASGS---------------------------------------------S--------
EsDELLA     149 AAPCA-------------------------------------------------------
TauDELLA    149 AQYSGA---------------------------------------------QVN--AN--
HcDELLA     148 AQYNGA---------------------------------------------QVN--AN--
SfaDELLA    148 AQYNGS---------------------------------------------RVN--AN--
PpDELLAa    149 APHN--------------------------------------------------------
PpDELLAb    148 TPQN--------------------------------------------------------
SfDELLAa    163 GLPCAT---------------------------------------------TSY--PA--
MpDELLA     154 ALPS-------------------------------------------------A--AA--
AaDELLA     240 GNSHP------------VSSYGVVVTSGGGPMGIPSLQDARGLP--QQMMGA--------
ApDELLA     207 GNSHP------------VSSYGVVVTSGGGPMGVPSLQDARGLP--QQMMGA--------
```

```
AtRGA1      151 ---AIDSSSSSNNQNKR---------------------------------------LKSC
AtGAI1      115 ---AIDSASSSNQGGGG---------------------------------------DTY-
AtRGL2      140 ---EEVFDEEASSKRIR---------------------------------------LGSW
MtDELLA     123 ---EENT-AAK-----R---------------------------------------MKTW
SlDELLA     159 ---SNKR-HRS-----T---------------------------------------TSSF
HvSLN1      163 ---LSADS-VRDPKRMR---------------------------------------TGGS
TaRHT-B1    166 ---LSADSVVRDPKRMR---------------------------------------TGGS
ZmD8        172 ---DPSTDSAREPKRMR---------------------------------------TGGG
OsSLR1      170 ---PSAADSARDTKRMR---------------------------------------TGGG
AtrDELLA1   146 ---DFGTSSQI-----S---------------------------------------SLVY
AtrDELLA2   112 ------------------------------------------------------------
PtRGA1      188 ------------------------------------------------------------
CrDELLAa    291 --PATTSSQNSNNKATYHEAP-SVRFQQQLHRK-VNQEE---------VKITEPEVTADL
CrDELLAb    298 PPPNPASSQSSSNKVPRTGSPSPVHVQRQCHRPPQNQGT---------VRTSTAMVMASV
SkDELLA     159 ------------------------------------------------------------
SmDELLAa    107 ------------------------------------------------------------
EsDELLA     154 ------------------------------------------------------------
TauDELLA    160 -----------------------------------------------------GPTTPVF
HcDELLA     159 -----------------------------------------------------RPITPAF
SfaDELLA    159 -----------------------------------------------------RPITPAF
PpDELLAa    153 ------------------------------------------------------------
PpDELLAb    152 ------------------------------------------------------------
SfDELLAa    174 -----------------------------------------------------SSKSCSM
MpDELLA     161 -----------------------------------------------------AAVMPDM
AaDELLA     278 -------S--------------DSSQQQILHRTHLSDAGMSRSLKNAGVNSADSQIL--S
ApDELLA     245 -------S--------------DSSQQQILHRTHLSDAGMSRSLKNAGVNSADSQIL--S

AtRGA1      169 SSPDSMVTS-TSTGTQI-----GGVIGT---TVTTTTTTTTAAGESTRSVILVDSQE-NG
AtGAI1      132 ------TTN-----KRL-----K---------CSNGVVETTTATAESTRHVVLVDSQE-NG
AtRGL2      158 ---------------------------------------CESSDESTRSVVLVDSQE-TG
MtDELLA     135 SEPE---------------------------SEPAVVMSPPPAVENTRPVVLVDTQE-TG
SlDELLA     171 STTS---------------------------------SSMVTDSSATRPVVLVDSQE-TG
HvSLN1      180 STSSSSSSSSSLGGGAA-----RSSVVE---AAPPVAA---AAAAPALPVVVVDTQE-AG
TaRHT-B1    184 STSSS-SSSSSLGGGGA-----RSSVVE---AAPPVAA---AAGAPALPVVVVDTQE-AG
ZmD8        190 STSSSSSSSSSMDGGRT-----RSSVVE---AAPPATQASAAANGPAVPVVVVDTQE-AG
OsSLR1      188 STSSSSSSSSSLGGGAS-----RGSVVE---AAPPATQGAAAANAPAVPVVVVDTQE-AG
AtrDELLA1   159 QSPP---------------------------REKKRIK--AAPPPEPRPVVVVDSQE-TG
AtrDELLA2   112 -----------------------------------------NRGSFSGAAILTHEED-SG
PtRGA1      188 ---------RSFSAQSSDEGSSLS----------------STRLGTAQQSIDNGAQE-SG
CrDELLAa    338 SPSSSSPMSVSYQEHCSPQDKDSIYHMR-YAPSKHANSQTMQTCPYTEVVDYENVQE-SG
CrDELLAb    349 SPSNSSPVSISYQDHSSPHDKEASYVHIQSPSAKRTRSQTVHECPYDDISNDENAQE-SG
SkDELLA     159 ---------VLYNDLQSPPSSSSA----------------VLQSMPSMAMPPTTTEE-LG
SmDELLAa    107 ---------SSTSPHGIPPHAA-G----------------GMTSAAAMPTIQESDEL-SG
EsDELLA     154 --------------GASQ---------------VHSSRP-TGAIVQQQQLQMGEDEN-NG
TauDELLA    167 QPSLTTIDTAQYANGGNV---------------LRGRNS-TGAVAEQQQLSMGEDEN-SG
HcDELLA     166 QPGLTTTNAPPYVG---------------------GQTS-AGAVAEQQQIPMGDDET-SG
SfaDELLA    166 QTGLTNTDTVPYGE---------------------DHAS-TGAVAEQQQIQMGDDET-SG
PpDELLAa    153 --------------GTSQ---------------VNAIRT-TAAGLEQQLNKMGEDEN-NG
PpDELLAb    152 --------------DASQ---------------LNANRA-TGAVLEQQPSPMGEDED-NG
SfDELLAa    181 LPHQQSTDSSS---ETHV---------------LPM--M-ESRNHQRPQVNEDEQED-NG
MpDELLA     168 YPADDIS-----------------------------DSM-AGALCPQ---EDTGVEE-SG
AaDELLA     315 DPSLNSHMSMMMEEAANPM-------VNRLPQHEQRDLGPSPADSQQVDAAPHQQEE-SG
ApDELLA     282 DPSLNSHMSMMMEEAANPM-------VNRLPQHEQQDLGPSPADSQQVDAAPHQQEE-SG

AtRGA1      219 VRLVHALMACAEAIQQNNLTLAEALVKQIGCLAV----SQAGAMRKVATYFAEALARRIY
AtGAI1      167 VRLVHALLACAEAVQKENLTVAEALVKQIGFLAV----SQIGAMRKVATYFAEALARRIY
AtRGL2      178 VRLVHALVACAEAIHQENLNLADALVKRVGTLAG----SQAGAMGKVATYFAQALARRIY
MtDELLA     167 VRLVHTLMACAEAIQQKNLKLAEALVKHISLLAS----LQTGAMRKVASYFAQALARRIY
SlDELLA     197 VRLVHTLMACAEAVQQENLTLADQLVRHIGILAV----SQSGAMRKVATYFAEALARRIY
HvSLN1      228 IRLVHALLACAEAVQQENLSAAEALVKQIPLLAA----SQGGAMRKVAAYFGEALARRVF
TaRHT-B1    231 IRLVHALLACAEAVQQENFSAAEALVKQIPLLAA----SQGGAMRKVAAYFGEALARRVF
ZmD8        241 IRLVHALLACAEAVQQENFSAAEALVKQIPMLAS----SQGGAMRKVAAYFGEALARRVY
OsSLR1      239 IRLVHALLACAEAVQQENFAAAEALVKQIPTLAA----SQGGAMRKVAAYFGEALARRVY
AtrDELLA1   189 IRLVHTLMACAEAVQQENMNAAEALVKQIGMLAV----SQAGAMRKVATFFAEALARRIF
AtrDELLA2   130 IRLIHLLMSCAGSVERGEREIALKLVQEMRLLLCR---NITGVIGKVAVFFVDALFWRLS
PtRGA1      222 IRVVHLLMGCAEAIQRNNLKVASDLVREIRMTVNS---APCGAMDKVASHFVEALARRIC
CrDELLAa    396 IKLVHLLMACAEAIQNNALAAAVDMVREIKRLAS----STRGTMSKVANYFVESLARCIY
CrDELLAb    408 IKLVHLLMACAEAIQNDELAAAVDMVREIKRLAS----CTSGAMSKIASYFAESLSQRIY
SkDELLA     193 VQLVHLLLACADAVQRREIPAAGDMARKLRSMLAGGAADSSGAMGRVAAHFVEGLCRRIF
SmDELLAa    140 VRLVHLLLACANAVQRGDLAAAGDMVAQLRILVAH-PSSSSSAMARVATQFVEALSRRIQ
EsDELLA     183 VQLVHSLLACAESIQRGNLILAEETLHRIQMLAL----P-PGPMGKVATHFIDALNRRIY
TauDELLA    210 VQLVHSLLGCAEAIQRGNLNLAEQTLHRIQLLGL----P-PGPMGKVATHFIDALARRVY
HcDELLA     203 VQLVHSLLACAEAIQRGNLKLAEETLHRMQLLGL----P-PGPMGKVATHFIDALARRVY
SfaDELLA    203 VQLVHSLLACAEAIQRGNLKLAEQTLHRMQVLGL----P-PGPMGKVATHFIDALVRRVY
PpDELLAa    182 IQLVHSLLACAESIQRGNLSFAEETLRRIELLSL----P-PGPMGKVATHFIGALTRRIY
PpDELLAb    181 VRLVHSLLACAESIQRGNLNLAEQTLRRIQLLSL----P-PGPMGKVATHFIDALTCRIY
SfDELLAa    219 VQLVHSLLACAEAVQHGDLVRAEETVRHIQLLAS----P-PGPMGKVAAHFIEALTRRIY
MpDELLA     194 VRLVHLLLACAEAVQRSDVRMAEDTVRRIQMLAT----PQRGPMGKVAAHFVEALARRIF
AaDELLA     367 VRLVHLLVTCAQAVHSNDMVRADETVRQIQDLAYLSR-GSTGPMEKVAVHFVDALVRRIY
ApDELLA     334 VRLVHLLVTCAQAVHSNDMVRADETVRQIQDLAYLSR-GSTGPMEKVAVHFVDALVRRIY
```

```
AtRGA1      275 RLSPPQN----------QIDHCLSDTLQMHFYETCPYLKFAHFTANQAILEAFEGKKRVH
AtGAI1      223 RLSPSQS----------PIDHSLSDTLQMHFYETCPYLKFAHFTANQAILEAFQGKKRVH
AtRGL2      234 RDYTAETD------VCAAVNPSFEEVLEMHFYESCPYLKFAHFTANQAILEAVTTARRVH
MtDELLA     223 G-NP-EE----------TIDSSFSEILHMHFYESSPYLKFAHFTANQAILEAFAGAGRVH
SlDELLA     253 KIYP-QD----------SMESSYTDVLQMHFYETCPYLKFAHFTANQAILEAFTGCNKVH
HvSLN1      284 RFRPQPDS--------SLLDAAFADLLHAHFYESCPYLKFAHFTANQAILEAFAGCRRVH
TaRHT-B1    287 RFRPQPDS--------SLLDAAFADPIHAHFYESCPYLKFAHFTANQAILEAFAGCRRVH
ZmD8        297 RFRPPPDS--------SLLDAAFADLLHAHFYESCPYLKFAHFTANQAILEAFAGCRRVH
OsSLR1      295 RFRPA-DS--------TLLDAAFADLLHAHFYESCPYLKFAHFTANQAILEAFAGCHRVH
AtrDELLA1   245 RFHP-QD----------TVD-LFSDILQMHFYETCPYLKFAHFTANQAILEAFAGKKRVH
AtrDELLA2   187 GHPSNR------------VDSGESEFLYHHFYEGCPYLKFAHFTCNQAILEAFDGCDEVH
PtRGA1      279 GLNGAE--------SNMSQVDAQSEILYHHFYEVCPYLKFAHFTANQAILEAFEGHGSVH
CrDELLAa    452 PGNKCDWA-------YLCQADALSELLYANFYEALPYLKFAHFTANQAILEAFQGHKFVH
CrDELLAb    464 PASKDNWA-------RIYEAEAVSEMLYASFYEACPYLKFAHFTANQAILEAFQGHKVVH
SkDELLA     253 GGGGVGLGGIPGLDITGVSSATVDEILHFHYYETCPYLKFAHFTANQAILEAFEGQSQVH
SmDELLAa    199 NSCYNE-SS----DPGNTNNGAMDEILHFHFYETCPYLKFAHFTANQAILEALEGHKSVH
EsDELLA     238 GGASFSGN-----NVCSNQSDSLSELLHFHFYETCPYLKFAHFTANQAILEAFAGHRQVH
TauDELLA    265 GVASS-CG-----NNSSNHSDSLSELLHFHFYETCPYLKFAHFTANQAILEAFAGQKQVH
HcDELLA     258 GVASSSNG-----HNSSSQSDSLAELLHFHFYETCPYLKFAHFTANQAILEAFAGHKQVH
SfaDELLA    258 GAASSNG-------NNSNQSDSLSELLHFHFYETCPYLKFAHFTANQAILEAFAGHKQVH
PpDELLAa    237 GVASSSGN-----NSSSNQSDSLLGLLHFYFYESCPFLRFAHFTANQAILEAVTGLKEVH
PpDELLAb    236 GVAFSSGN-----NVGSNQSDSLSELLHFHFYETCPYLKFAHFTANQAILEAFAGQKQVH
SfDELLAa    274 GGTSSSQDSSSCNVVSYESNNYLSELLHFQYYETCPYLKFAHFTSNQAILEAFEGEKRVH
MpDELLA     250 GISAE------------PTSDPLTELLHFQFYETCPYLKFAHFTANQAILEAVQNHKRIH
AaDELLA     426 GFNNNDS-------VVGMQTDCLSELLHFQFYEICPYLKFAHFTANQAILEAFEGEQNVH
ApDELLA     393 GFNNNDS-------VGGMQTDCLSELLHFQFYEICPYLKFAHFTANQAILEAFEGEQNVH

AtRGA1      325 VIDFSMNQGLQWPALMQALALREGGPPTFRLTGIGPPAPDNSDHLHEVGCKLAQLAEAIH
AtGAI1      273 VIDFSMSQGLQWPALMQALALRPGGPPVFRLTGIGPPAPDNFDYLHEVGCKLAHLAEAIH
AtRGL2      288 VIDLGLNQGMQWPALMQALALRPGGPPSFRLTGIGPPQTENSDSLQQLGWKLAQFAQNMG
MtDELLA     271 VIDFGLKQGMQWPALMQALALRPGGPPTFRLTGIGPPQADNTDALQQVGWKLAQLAQTIG
SlDELLA     302 VIDFSLKQGMQWPALMQALALRPGGPPAFRLTGIGPPQPDNTDALQQVGWKLAQLAETIG
HvSLN1      336 VVDFGIKQGMQWPALLQALALRPGGPPSFRLTGVGPPQPDETDALQQVGWKLAQFAHTIR
TaRHT-B1    339 VVDFGIKQGMQWPALLQALALRPGGPPSFRLTGVGPPQPDETDALQQVGWKLAQFAHTIR
ZmD8        349 VVDFGIKQGMQWPALLQALALRPGGPPSFRLTGVGPPQPDETDALQQVGWKLAQFAHTIR
OsSLR1      346 VVDFGIKQGMQWPALLQALALRPGGPPSFRLTGVGPPQPDETDALQQVGWKLAQFAHTIR
AtrDELLA1   293 VIDFSMKQGMQWPALMQALALRPGGPPAFRLTGIGPPQPDNTDPLQQVGWKLAQLAETIH
AtrDELLA2   235 VIDFNLIHGLQWPALIQALALRPGGPPFLRLTGIGPPSPDGRDTIREVGIRLAELARSVN
PtRGA1      331 VIDLNLMHGLQWPALIQALALRPGGPPLLRLTAIGPRQPDGRDVLQEIGMKLAQFAESVN
CrDELLAa    505 IIDFNLMQGSQWPALIQALADREEGPPYLRMTGIGLPHQDNKDVLQEVGKELAELAHSVN
CrDELLAb    517 IIDFNLMQGSQWPELIKALAVRSEGPPHLRMTGIGPPRPDNKDVLQEVGVKLAELAGSVN
SkDELLA     313 VVDFNLEYGLQWPALIQALALRPGGPPQLRLTGIGPPQPGGKDLLQEIGLKLAQMAESVN
SmDELLAa    254 VVDLDLQYGLQWPALIQALALRPGGPPTLRLTGIGPPQPHRHDLLHEIGLKLAQLADSVN
EsDELLA     293 VIDFNLMHGLQWPALIQALALRPGGPPRLRLTGVGPPQPGGNDVLQEIGSKLRQLADTVK
TauDELLA    319 VIDFNLMHGLQWPALIQALALRSGGPPRLRLTGIGPPQPGGNDVLQEIGKKLRQLADTVK
HcDELLA     313 VIDFNLMHGLQWPALIQALALRPGGPPRLRLTGIGPPQAGGNDVLQEIGKKLRQLADTVK
SfaDELLA    311 VIDFNLMHGLQWPALIQALALRPGGPPRLRLTGIGPPQPGGIDVLQEIGKKLRQLADTVK
PpDELLAa    292 VIDFNLMQGLQWPALIQALSLRQGGPPRLRLTGIGPPQPSGSDTLQEIGTKLAELAKTVR
PpDELLAb    291 VIDFNLMHGLQWPALIQALALRPGGPPRLRLTGIGPPQSGGSDVLQEIGMKLAQLAETVK
SfDELLAa    334 VIDFNLMHGLQWPALIQALALRPGGPPSLRLTGIGPPQAGGNNGLQEIGMKLAQLAASVN
MpDELLA     298 VIDCNLMHGLQWPALIQALALRPGGPPILRLTGIGPPHQSGNDVLQEIGMKLAQLADLVS
AaDELLA     479 IIDFNLMHGLQWPALIQALALRPGGPPFLRLTGIGLPQPGGTDVLQEIGTKLAQLAGSVN
ApDELLA     446 IIDFNLMHGLQWPALIQALALRPGGPPFLRLTGIGLPQPGGTDVLQEIGTKLAQLAGSVN

AtRGA1      385 VEFEYRGFVANSLADLDASMLELR-----PSDTEAVAVNSVFELHKLLGR----------
AtGAI1      333 VEFEYRGFVANTLADLDASMLELR-----PSEIESVAVNSVFELHKLLGR----------
AtRGL2      348 VEFEFKGLAAESLSDLEPEMFETR-----PES-ETLVVNSVFELHRLLAR----------
MtDELLA     331 VQFEFRGFVCNSIADLDPNMLEIR-----PGE--AVAVNSVFELHTMLAR----------
SlDELLA     362 VEFEFRGFVANSLADLDATILDIR-----PSETEAVAINSVFELHRLLSR----------
HvSLN1      396 VDFQYRGLVAATLADLEPFMLQPEGEEDPNEEPEVIAVNSVFEMHRLLAQ----------
TaRHT-B1    399 VDFQYRGLVAATLADLEPFMLQPEGEEDPNEEPEVIAVNSVFEMHRLLAQ----------
ZmD8        409 VDFQYRGLVAATLADLEPFMLQPEGDD-TDDEPEVIAVNSVFELHRLLAQ----------
OsSLR1      406 VDFQYRGLVAATLADLEPFMLQPEGEADANEEPEVIAVNSVFELHRLLAQ----------
AtrDELLA1   353 VEFEYRGFVARSLADLEAYMLDVR-----PSDVEVVAVNSIFELHNLLAQ----------
AtrDELLA2   295 VRFAFRGVATQRLEDLKPWMIHVR-----ST--ETVAVNSVFVLHRLLYT----------
PtRGA1      391 VEFDFRGVMADKLEDIKPWMFQVK-----PD--EVVAVNSVLQLHRLLYID---------
CrDELLAa    565 VKFSFRGMVATKLEDVKPWYFEVN-----PG--EAIAVNSILQMHRLLYGCVG-------
CrDELLAb    577 VEFSFRGMVAAKLDDVKPWYFEVK-----PG--EAIAVNSILQMHRLLYGHVA-------
SkDELLA     373 VEFTFHGVVAARLEDVRPWMLTCR-----SG--EAVAVNSVFQLHATLLDGEG-------
SmDELLAa    314 VDFAFHGVVAARLNDVQPWMLTVR-----RG--EAVAVNSVFQMHKALVE----------
EsDELLA     353 VEFEFRGVIAVKLDDIKPWMLHVR-----HG--EAVAVNSVLQLHKLLYHA-G-------
TauDELLA    379 VEFEFRGVVAVKLDDIKPWMLQVR-----HG--EAVAVNSVLQLHKLLYSA-G-------
HcDELLA     373 VEFEFRGVVAVKLDDIKPWMLQGR-----HG--EAVAVNSVLQLHKLLYSD-G-------
SfaDELLA    371 VEFEFRGVVAVKLDDIKPWMLQVR-----HG--EAVAVNSVLQLHKLLFSD-G-------
PpDELLAa    352 VDFEFRGVIAVKLDDIKPWMLQIR-----HG--EAVAVNSVLQLHKLLYSA-G-------
PpDELLAb    351 VEFEFRGVVAVKLDDIKPWMLQIC-----HG--EAVAVNSVFQLHKLLYSA-G-------
SfDELLAa    394 IEFDFRGVVALKLNEVKPWMLQVL-----PG--EVVAVNSVLQLHRFLNSDGG-------
MpDELLA     358 VQFEFRGVYAQKLDDVKPYMLHVR-----QG--EAVAVNAVMQLHRLLHNDNN-------
AaDELLA     539 VKFDFRGVVATKLDDVKPWMLQVR-----QG--EAVAVNSILQLHRLLGPHPEELWQVGN
ApDELLA     506 VKFDFRGVVATKLDDVKPWMLQAT-----LA--TSLQ-----------------------
```

```
AtRGA1      430 ----------PGGIEKVLGVVKQIKPVIFTVVEQESNHNGPVFLDRFTESLHYYSTLFDS
AtGAI1      378 ----------PGAIDKVLGVVNQIKPEIFTVVEQESNHNSPIFLDRFTESLHYYSTLFDS
AtRGL2      392 ----------SGSIEKLLNTVKAIKPSIVTVVEQEANHNGIVFLDRFNEALHYYSSLFDS
MtDELLA     374 ----------PGSVEKVLNTVKKINPKIVTIVEQEANHNGPVFVDRFTEALHYYSSLFDS
SlDELLA     407 ----------PGAIEKVLNSIKQINPKIVTLVEQEANHNAGVFIDRFNEALHYYSTMFDS
HvSLN1      446 ----------PGALEKVLGTVRAVRPRIVTVVEQEANHNSGSFLDRFTESLHYYSTMFDS
TaRHT-B1    449 ----------PGALEKVLGTVRAVRPRIVTVVEQEANHNSGTFLDRFTESLHYYSTMFDS
ZmD8        458 ----------PGALEKVLGTVRAVRPRIVTVVEQEANHNSGTFLDRFTESLHYYSTMFDS
OsSLR1      456 ----------PGALEKVLGTVHAVRPRIVTVVEQEANHNSGSFLDRFTESLHYYSTMFDS
AtrDELLA1   398 ----------PSALDKVLASVRALRPKIVTIVEQEANHNGPVFMDRFTEALHYYSTLFDS
AtrDELLA2   338 ------TPDQTQPIKPVLNWVRELGPKILTVVEQEASHNGLGFVDRFTEALHYYSAMFDS
PtRGA1      435 ------APTGSSPIDVVLRSIGSLRPKIVTVVEHEANHNGPVFLDRFVEALHYYSTMFDS
CrDELLAa    611 --S----DPSKAPIDEVLSFIKSLKPKVVTLVEQEANHNGSIFLERFVEALHYYSTMFDS
CrDELLAb    623 --S----DPSKALIDEVLSSIKSLNPKVVTVVEQEANHNSNMFLERFVEALHYYSTMFDS
SkDELLA     419 --AAGSSPVAPSPVTEVLRWVRGLNPRIVTVVEQDADHNGVDFLDRFMAALHYYSTMFDS
SmDELLAa    357 ----------EPPIDEVLRLVRNLKPKIVTLVEQDADHNSPVFMERFMAALHYYSTMFDS
EsDELLA     398 --S-------VPPIEEVLRSVRELKPKIFTIVEHEANHNNPSFLGRFTEALHYYSTMFDS
TauDELLA    424 --P-------VRAIDEVLRSVRALKPKIFTIVEHEANHNNPSFLGRFTEALHYYSTMFDS
HcDELLA     418 --P-------VRAIDEVLRSVRALKPKIFTIVEHEANHNNPSFLGRFTEALHYYSTMFDS
SfaDELLA    416 --P-------VRAIDEVLRSVRTLKPKIFTIVEHEANHNNPSFLGRFTEALHYYSTMFDS
PpDELLAa    397 --P-------EAPIDAVLLLVRELKPKIFTIVEHEANHNQPSFLGRFIEALHYYSTMFDA
PpDELLAb    396 --S-------VIPIDEVLRSARALKPKIFTIVEHEANHNQPSFLGRFTEALHYYSTMFDS
SfDELLAa    440 --P-------VLAIDEVLHSILGLKPKIVTVVEHEANHNVSGFLDRFTEALHYYSTMFDS
MpDELLA     404 --C-------VPAVHEVLQSMRSLNPKIVTLVEHEANHNSPVFLDRFMEALHYYSTMFDS
AaDELLA     592 PGDISAVKGPVPAIAEVLQSVRSLNPKIVTLVEHEANHNSPVFIDRFMEALHYYSTMFDS
ApDELLA         ------------------------------------------------------------

AtRGA1      480 LEGVP-----------------NSQDKVMSEVYLGKQICNLVACEGPDRVERHETLSQWG
AtGAI1      428 LEGVP-----------------SGQDKVMSEVYLGKQICNVVACDGPDRVERHETLSQWR
AtRGL2      442 LEDSY---------------SLPSQDRVMSEVYLGRQILNVVAAEGSDRVERHETAAQWR
MtDELLA     424 LEGSNSSSNNSNSNS--TGLGSPSQDLLMSEIYLGKQICNVVAYEGVDRVERHETLTQWR
SlDELLA     457 LESSGSSSSASPTGI-LPQPPVNNQDLVMSEVYLGRQICNVVACEGSDRVERHETLNQWR
HvSLN1      496 LEGGSSGGPSEVSSGGAAPAAAAGTDQVMSEVYLGRQICNVVACEGTERTERHETLGQWR
TaRHT-B1    499 LEGGSSGGPSEVSSGAAAAPAAAGTDQVMSEVYLGRQICNVVACEGAERTERHETLGQWR
ZmD8        508 LEGAGAGSGQS---TDASPAAAGGTDQVMSEVYLGRQICNVVACEGAERTERHETLGQWR
OsSLR1      506 LEGGSSGQAEL---SPPAAGGGGGTDQVMSEVYLGRQICNVVACEGAERTERHETLGQWR
AtrDELLA1   448 LEGCG--------------LPPGSNDQVMSEVYLGRQICNIVACEGADRVERHETLTQWR
AtrDELLA2   392 MEGSN--------------R----NNQAFAELYLEREIKNIVCCEGSERVERHEPLTRWR
PtRGA1      489 LEACN--------------VLPNSMEKLLAELYIQKEICNIVACEGRYRIERHETLSHWR
CrDELLAa    665 LEASS--------------LDPQSSEMACAEAYLAREITNVLACEGAERVERHEPLSQWR
CrDELLAb    677 LEASS--------------LDPLGPEMVCSEMYLGREIANIVAREGAERVERHEPLSAWR
SkDELLA     477 LEACN--------------LAAGSLEQVVAEAYLGREVVDIVAADGPERRERHETLEQWR
SmDELLAa    407 LEACN--------------LAPGSVEQMVAETYLGQEIGNIVACEGAARTERHETLTQWR
EsDELLA     449 LEACN--------------LPSESSEQVLAEMYLGREIYNIVACEDAARIERHENLVQWR
TauDELLA    475 LEACN--------------LPSESNEQVLAEMYLGREIYNIVACEDAARVERHENLLQWR
HcDELLA     469 LEASN--------------LPSESNEQVLAKMYLGREIYNIVACEDAARTERHENLVQWR
SfaDELLA    467 LEASN--------------FPSESNEQVLAKLYLGREIYNIVACEDAARTERHENLVQWR
PpDELLAa    448 LEACN--------------LPSENNEQVLIEMYLGREIYNIVACEDGARTERHENLFQWR
PpDELLAb    447 LEACS--------------LPSDSSEQVLAEMYLGREINNIVACEDAARVERHENLVQWQ
SfDELLAa    491 LEACN--------------LQPQSSEQLLAEMYLGQEICNIIACEGVARVERHENLEQWR
MpDELLA     455 LEACS--------------SSSQSSEQLLAEMYVGREICNIVACEGFDRVERHENLVQWR
AaDELLA     652 LEACN--------------HTPDSGTQVLAEMYLGREICNIVACEGPGRVERHENLSQWQ
ApDELLA         ------------------------------------------------------------

AtRGA1      523 NRFGSSGLAPAHLGSNAFKQASMLLSVFNSGQGYRVEESNGCLMLGWHTRPLITTSAWKL
AtGAI1      471 NRFGSAGFAAAHIGSNAFKQASMLLALFNGGEGYRVEESDGCLMLGWHTRPLIATSAWKL
AtRGL2      487 IRMKSAGFDPIHLGSSAFKQASMLLSLYATGDGYRVEENDGCLMIGWQTRPLITTSAWKL
MtDELLA     482 SRMGSAGFEPVHLGSNAFKQASTLLALFAGGDGYRVEENNGCLMLGWHTRSLIATSAWKL
SlDELLA     516 VRMNSSGFDPVHLGSNAFKQASMLLALFAGGDGYRVEENDGCLMLGWHTRPLIATSAWKL
HvSLN1      556 NRLGNAGFETVHLGSNAYKQASTLLALFAGGDGYKVEEKEGCLTLGWHTRPLIATSAWRL
TaRHT-B1    559 NRLGNAGFETVHLGSNAYKQASTLLALFAGGDGYKVEEKEGCLTLGWHTRPLIATSAWRL
ZmD8        565 SRLGGSGFAPVHLGSNAYKQASTLLALFAGGDGYRVEEKDGCLTLGWHTRPLIATSAWRV
OsSLR1      563 NRLGRAGFEPVHLGSNAYKQASTLLALFAGGDGYRVEEKEGCLTLGWHTRPLIATSAWRV
AtrDELLA1   494 ARMGAAGFAPVHLGSNAFRQASMLLTLFSGGDGYKVEENNGCLMLAWHTRPLIATSAWQI
AtrDELLA2   434 GRFDKQDSNR-------LDSVRMRLDRRV-C----CSLFFRKMDLGWRRGMDA-------
PtRGA1      535 VRLGRAGFRPSHLGSNAFKQARMLLTLFS-GEGYTVEENNGSLTLGWHSRPLIAASAWQG
CrDELLAa    711 KRMSNAGFKPLHLGSNAFNKVSVLLKVFS-GEGYTVEENKGCLTLGWHNRPLIASSAWQC
CrDELLAb    723 KRMSNAGFKQVHLGSNAFDQVSYMLKYFS-GEGYTVEENRGCLTLGWHNRPLIAASAWEC
SkDELLA     523 SRMISAGFQPLFLGSNAFRQASMLLTLFS-GDGYRVVENGGCLTLGWHSRSLIAASAWRC
SmDELLAa    453 IRMARSGFQPLYLGSNAFKQANMLLTLFS-GDGYRVEEKDGCLTLGWHSRPLVAASAWEC
EsDELLA     495 LRLLKAGYRPIQLGLNAFKQASMLLRMFS-GEGYRVEEKLGCLTLGWHTRPLIAASAWQC
TauDELLA    521 QRLLKAGYRPIQLGLNAFKQASMLLTMFS-GEGYRVEEKLGCLTLGWHTRPLIAASAWQC
HcDELLA     515 LRLFKAGFRPIQLGLNVFKQASMLLTMFS-GEGYRVEERLGCLTLGWHTRPLIAASAWQC
SfaDELLA    513 LRLLKAGFRPIQLGLNALKQASMLLTMFS-GEGYRIEERLDCLTLGWHARPLIAASAWQC
PpDELLAa    494 LRLLKAGYRPIQLGLNAFKQASMLLTMFS-GEGYRVEEKLGCLTLGWHSRPLIAASAWKC
PpDELLAb    493 MRMLKAGYRPIQLGLNAFKQASMLLTMFS-GDGYRVEEKLGCLTLGWHTRPLISASAWQC
SfDELLAa    537 QRIAKAGFRPLQLGSTALKQAKLLLSLFP-GDGYRVEENNGCLTLGWHTRPLIAFSAWQC
MpDELLA     501 RRMTDAGFQLRHLGSNAFKQASMLLTLFP-GDGYRVEENNGCLTLGWHSRPLIAASAWHC
AaDELLA     698 LRMCNAGFQTRHLGANAYRQASMLLSLFS-AEGYSVEETNGFLLLKWHDRPLMAASAWQC
ApDELLA         ------------------------------------------------------------
```

```
AtRGA1      583 STAAY-----------------NSQDKVMSEVYLGKQICNLVACEGPDRVERHETLSQWG
AtGAI1      531 STN-------------------SGQDKVMSEVYLGKQICNVVACDGPDRVERHETLSQWR
AtRGL2      547 A-------------------SLPSQDRVMSEVYLGRQILNVVAAEGSDRVERHETAAQWR
MtDELLA     542 PQNESK-------NS--TGLGSPSQDLLMSEIYLGKQICNVVAYEGVDRVERHETLTQWR
SlDELLA     576 LPDSGTGAGEVELGI-LPQPPVNNQDLVMSEVYLGRQICNVVACEGSDRVERHETLNQWR
HvSLN1      616 AAP----------SGGAAPAAAAGTDQVMSEVYLGRQICNVVACEGTERTERHETLGQWR
TaRHT-B1    619 AAP----------SGAAAAPAAAGTDQVMSEVYLGRQICNVVACEGAERTERHETLGQWR
ZmD8        625 AAAAAP--------TDASPAAAGGTDQVMSEVYLGRQICNVVACEGAERTERHETLGQWR
OsSLR1      623 AAA-----------SPPAAGGGGGTDQVMSEVYLGRQICNVVACEGAERTERHETLGQWR
AtrDELLA1   554 ALP----------------LPPGSNDQVMSEVYLGRQICNIVACEGADRVERHETLTQWR
AtrDELLA2       -------------------R----NNQAFAELYLEREIKNIVCCEGSERVERHEPLTRWR
PtRGA1      594 S------------------VLPNSMEKLLAELYIQKEICNIVACEGRYRIERHETLSHWR
CrDELLAa    770 G------------------LDPQSSEMACAEAYLAREITNVLACEGAERVERHEPLSQWR
CrDELLAb    782 G------------------LDPLGPEMVCSEMYLGREIANIVAREGAERVERHEPLSAWR
SkDELLA     582 S------------------LAAGSLEQVVAEAYLGREVVDIVAADGPERRERHETLEQWR
SmDELLAa    512 C------------------LAPGSVEQMVAETYLGQEIGNIVACEGAARTERHETLTQWR
EsDELLA     554 A------------------LPSESSEQVLAEMYLGREIYNIVACEDAARIERHENLVQWR
TauDELLA    580 A------------------LPSESNEQVLAEMYLGREIYNIVACEDAARVERHENLLQWR
HcDELLA     574 A------------------LPSESNEQVLAKMYLGREIYNIVACEDAARTERHENLVQWR
SfaDELLA    572 A------------------FPSESNEQVLAKLYLGREIYNIVACEDAARTERHENLVQWR
PpDELLAa    553 A------------------LPSENNEQVLIEMYLGREIYNIVACEDGARTERHENLFQWR
PpDELLAb    552 A------------------LPSDSSEQVLAEMYLGREINNIVACEDAARVERHENLVQWQ
SfDELLAa    596 A------------------LQPQSSEQLLAEMYLGQEICNIIACEGVARVERHENLEQWR
MpDELLA     560 S------------------SSSQSSEQLLAEMYVGREICNIVACEGFDRVERHENLVQWR
AaDELLA     757 SQ-----------------HTPDSGTQVLAEMYLGREICNIVACEGPGRVERHENLSQWQ
ApDELLA         ------------------------------------------------------------

AtRGA1      523 NRFGSSGLAPAHLGSNAFKQASMLLSVFNSGQGYRVEESNGCLMLGWHTRPLITTSAWKL
AtGAI1      471 NRFGSAGFAAAHIGSNAFKQASMLLALFNGGEGYRVEESDGCLMLGWHTRPLIATSAWKL
AtRGL2      487 IRMKSAGFDPIHLGSSAFKQASMLLSLYATGDGYRVEENDGCLMIGWQTRPLITTSAWKL
MtDELLA     482 SRMGSAGFEPVHLGSNAFKQASTLLALFAGGDGYRVEENNGCLMLGWHTRSLIATSAWKL
SlDELLA     516 VRMNSSGFDPVHLGSNAFKQASMLLALFAGGDGYRVEENDGCLMLGWHTRPLIATSAWKL
HvSLN1      556 NRLGNAGFETVHLGSNAYKQASTLLALFAGGDGYKVEEKEGCLTLGWHTRPLIATSAWRL
TaRHT-B1    559 NRLGNAGFETVHLGSNAYKQASTLLALFAGGDGYKVEEKEGCLTLGWHTRPLIATSAWRL
ZmD8        565 SRLGGSGFAPVHLGSNAYKQASTLLALFAGGDGYRVEEKDGCLTLGWHTRPLIATSAWRV
OsSLR1      563 NRLGRAGFEPVHLGSNAYKQASTLLALFAGGDGYRVEEKEGCLTLGWHTRPLIATSAWRV
AtrDELLA1   494 ARMGAAGFAPVHLGSNAFRQASMLLTLFSGGDGYKVEENNGCLMLAWHTRPLIATSAWQI
AtrDELLA2   434 GRFDKQDSNR-------LDSVRMRLDRRV-C----CSLFFRKMDLGWRRGMDA-------
PtRGA1      535 VRLGRAGFRPSHLGSNAFKQARMLLTLFS-GEGYTVEENNGSLTLGWHSRPLIAASAWQG
CrDELLAa    711 KRMSNAGFKPLHLGSNAFNKVSVLLKVFS-GEGYTVEENKGCLTLGWHNRPLIASSAWQC
CrDELLAb    723 KRMSNAGFKQVHLGSNAFDQVSYMLKYFS-GEGYTVEENRGCLTLGWHNRPLIAASAWEC
SkDELLA     523 SRMISAGFQPLFLGSNAFRQASMLLTLFS-GDGYRVVENGGCLTLGWHSRSLIAASAWRC
SmDELLAa    453 IRMARSGFQPLYLGSNAFKQANMLLTLFS-GDGYRVEEKDGCLTLGWHSRPLVAASAWEC
EsDELLA     495 LRLLKAGYRPIQLGLNAFKQASMLLRMFS-GEGYRVEEKLGCLTLGWHTRPLIAASAWQC
TauDELLA    521 QRLLKAGYRPIQLGLNAFKQASMLLTMFS-GEGYRVEEKLGCLTLGWHTRPLIAASAWQC
HcDELLA     515 LRLFKAGFRPIQLGLNVFKQASMLLTMFS-GEGYRVEERLGCLTLGWHTRPLIAASAWQC
SfaDELLA    513 LRLLKAGFRPIQLGLNALKQASMLLTMFS-GEGYRIEERLDCLTLGWHARPLIAASAWQC
PpDELLAa    494 LRLLKAGYRPIQLGLNAFKQASMLLTMFS-GEGYRVEEKLGCLTLGWHSRPLIAASAWKC
PpDELLAb    493 MRMLKAGYRPIQLGLNAFKQASMLLTMFS-GDGYRVEEKLGCLTLGWHTRPLISASAWQC
SfDELLAa    537 QRIAKAGFRPLQLGSTALKQAKLLLSLFP-GDGYRVEENNGCLTLGWHTRPLIAFSAWQC
MpDELLA     501 RRMTDAGFQLRHLGSNAFKQASMLLTLFP-GDGYRVEENNGCLTLGWHSRPLIAASAWHC
AaDELLA     698 LRMCNAGFQTRHLGANAYRQASMLLSLFS-AEGYSVEETNGFLLLKWHDRPLMAASAWQC
ApDELLA         ------------------------------------------------------------
```


```
AtRGA1      583 STAAY-----------------NSQDKVMSEVYLGKQICNLVACEGPDRVERHETLSQWG
AtGAI1      531 STN-------------------SGQDKVMSEVYLGKQICNVVACDGPDRVERHETLSQWR
AtRGL2      547 A-------------------SLPSQDRVMSEVYLGRQILNVVAAEGSDRVERHETAAQWR
MtDELLA     542 PQNESK-------NS--TGLGSPSQDLLMSEIYLGKQICNVVAYEGVDRVERHETLTQWR
SlDELLA     576 LPDSGTGAGEVELGI-LPQPPVNNQDLVMSEVYLGRQICNVVACEGSDRVERHETLNQWR
HvSLN1      616 AAP----------SGGAAPAAAAGTDQVMSEVYLGRQICNVVACEGTERTERHETLGQWR
TaRHT-B1    619 AAP----------SGAAAAPAAAGTDQVMSEVYLGRQICNVVACEGAERTERHETLGQWR
ZmD8        625 AAAAAP--------TDASPAAAGGTDQVMSEVYLGRQICNVVACEGAERTERHETLGQWR
OsSLR1      623 AAA-----------SPPAAGGGGGTDQVMSEVYLGRQICNVVACEGAERTERHETLGQWR
AtrDELLA1   554 ALP----------------LPPGSNDQVMSEVYLGRQICNIVACEGADRVERHETLTQWR
AtrDELLA2       -------------------R----NNQAFAELYLEREIKNIVCCEGSERVERHEPLTRWR
PtRGA1      594 S------------------VLPNSMEKLLAELYIQKEICNIVACEGRYRIERHETLSHWR
CrDELLAa    770 G------------------LDPQSSEMACAEAYLAREITNVLACEGAERVERHEPLSQWR
CrDELLAb    782 G------------------LDPLGPEMVCSEMYLGREIANIVAREGAERVERHEPLSAWR
SkDELLA     582 S------------------LAAGSLEQVVAEAYLGREVVDIVAADGPERRERHETLEQWR
SmDELLAa    512 C------------------LAPGSVEQMVAETYLGQEIGNIVACEGAARTERHETLTQWR
EsDELLA     554 A------------------LPSESSEQVLAEMYLGREIYNIVACEDAARIERHENLVQWR
TauDELLA    580 A------------------LPSESNEQVLAEMYLGREIYNIVACEDAARVERHENLLQWR
HcDELLA     574 A------------------LPSESNEQVLAKMYLGREIYNIVACEDAARTERHENLVQWR
SfaDELLA    572 A------------------FPSESNEQVLAKLYLGREIYNIVACEDAARTERHENLVQWR
PpDELLAa    553 A------------------LPSENNEQVLIEMYLGREIYNIVACEDGARTERHENLFQWR
PpDELLAb    552 A------------------LPSDSSEQVLAEMYLGREINNIVACEDAARVERHENLVQWQ
SfDELLAa    596 A------------------LQPQSSEQLLAEMYLGQEICNIIACEGVARVERHENLEQWR
MpDELLA     560 S------------------SSSQSSEQLLAEMYVGREICNIVACEGFDRVERHENLVQWR
AaDELLA     757 SQ-----------------HTPDSGTQVLAEMYLGREICNIVACEGPGRVERHENLSQWQ
ApDELLA         ------------------------------------------------------------
```


**Supplemental Figure 1:** full length DELLA protein sequence alignment.

Alignment of the full-length DELLA protein sequences from selected vascular plants and bryophytes. Black shading indicates that at least 50% of the amino acids in a particular column are identical. Amino acids that are similar to the column-consensus peptide are shaded grey.

The sequences used in this figure are as follows: *Arabidopsis thaliana, Medicago truncatula, Solanum lycopersicum, Hordeum vulgare, Triticum aestivum, Zea mays, Oryza sativa, Amborella trichopoda* (angiosperms)*, Pinus tabuliformis* (gymnosperm)*, Ceratopteris richardii* (fern)*, Selaginella kraussiana, Selaginella moellendorfii* (lycophytes)*, Encalypta streptocarpa, Timmia austriaca, Hedwigia ciliata, Schwetschkeopsis fabronia, Physcomitrium patens, Sphagnum fallax* (mosses)*, Marchantia polymorpha* (liverwort)*, Anthoceros agrestis* and *Anthoceros punctatus* (hornworts).

**Figure S2.**

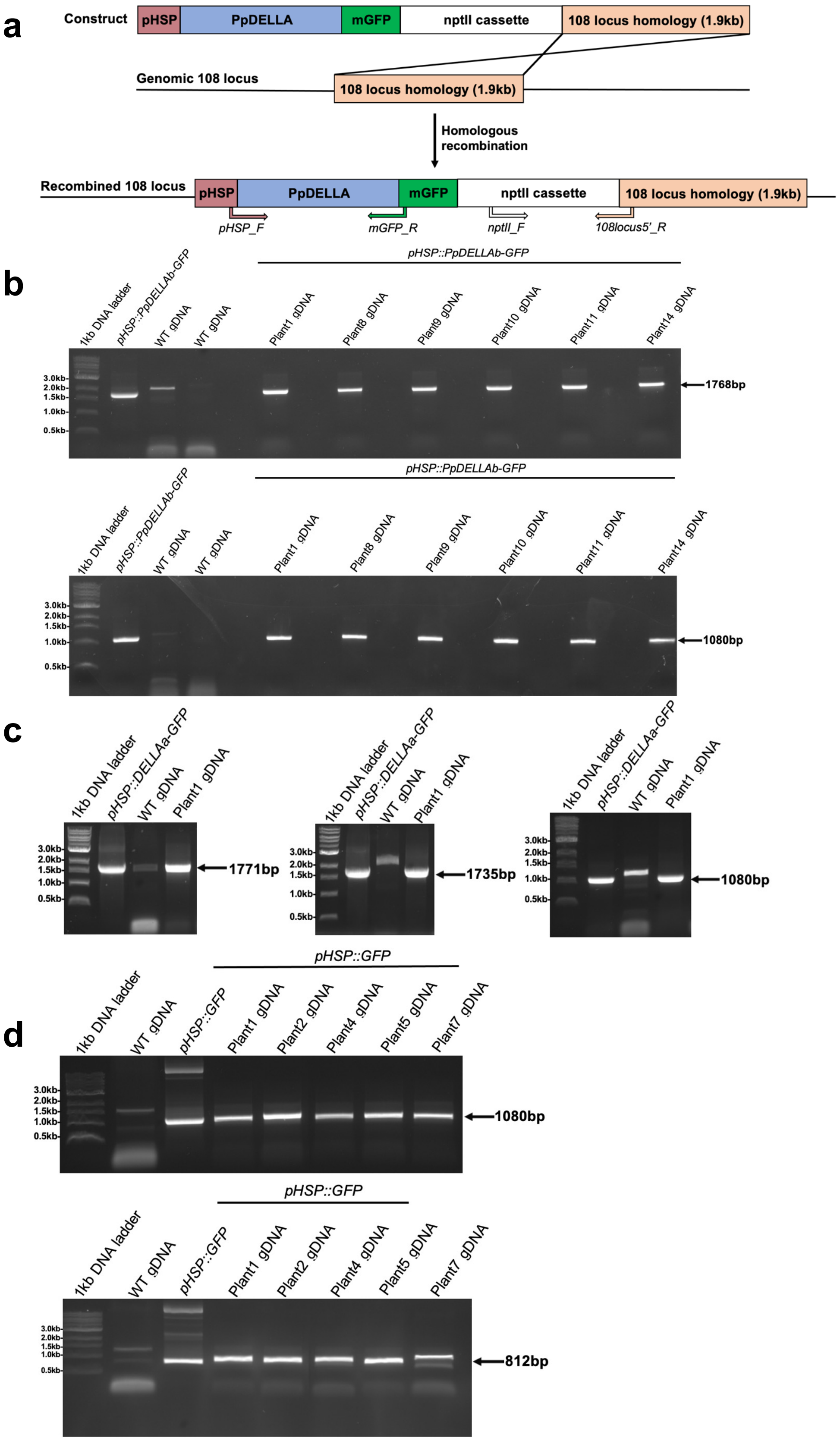

a
Construct
pHSP
PpDELLA
mGFP
nptII cassette
108 locus homology (1.9kb)
Genomic 108 locus
Homologous recombination
Recombined 108 locus
pHSP_F
mGFP_R
nptII_F
108locus5'_R
b
pHSP::PpDELLAb-GFP
1kb DNA ladder
WT gDNA
Plant1 gDNA
Plant8 gDNA
Plant9 gDNA
Plant10 gDNA
Plant11 gDNA
Plant14 gDNA
3.0kb
2.0kb
1.5kb
1.0kb
0.5kb
1768bp
1080bp
c
pHSP::DELLAa-GFP
1771bp
1735bp
1080bp
d
pHSP::GFP
Plant2 gDNA
Plant4 gDNA
Plant5 gDNA
Plant7 gDNA
812bp

**Supplemental Figure 2. Generation of *pHSP::PpDELLA-GFP* and *pHSP::GFP* transgenic lines.**

(A) Cloning strategy for inducibly overexpressing *Pp*DELLA-GFP proteins in *P. patens*. *PpDELLA*s were amplified by PCR from genomic DNA (as *PpDELLA*s are made up of one exon) and cloned in-frame with an *mGFP* gene in the pHSP::MCS::GFP-108-35SNPT moss transformation vector, which also contains a neomycin phosphotransferase (nptII) cassette for antibiotic selection in plant cells, and 1.9kb DNA of sequence homologous to the inert genomic 108 locus. The primers used for screening transformants for the presence of the construct are shown on the recombined 108 locus image.

(B) PCR-genotyping of *PpDELLAb* overexpression transformants to confirm *pHSP::DELLAb-GFP* integration in a *P. patens* wild-type (WT) background. Top panel: integration of the *pHSP::DELLAb-GFP* construct into the genome confirmed using the primers *pHSP_F* and *mGFP_R* in 6 transformants that survived two rounds of G418 selection. Genomic DNA from two wild-type plants (WT gDNA) and the transformation vector (*pHSP::PpDELLAb-GFP*) were included as negative and positive controls, respectively. Note the presence of a non-specific ~2kb amplification product in one WT gDNA sample. Bottom panel: integration of the *nptII* cassette into the *P. patens* genome was confirmed by PCR using the primers *nptII_F* and *108locus5'_R* in the same 6 transformants shown in the top panel. Controls as in the top panel. The PCR product from Plant1 in the top panel was also sequenced using the PCR primers to further confirm the presence of the construct.

(C) PCR-genotyping of *PpDELLAa* overexpression transformants to confirm integration of the *pHSP::DELLAa-GFP* construct in a *P. patens* wild-type (WT) background. From left to right: *pHSP_F* and *mGFP_R* primers, *XhoI-PpDELLAa_pHSP_F* and *mGFP_R* primers, *nptII_F* and *108locus5'_R* primers in the one plant that survived two rounds of G418 selection. In all cases, gDNA from a wild-type plant (gDNA) and the transformation vector (*pHSP::*DELLAa-GFP) were used as negative and positive controls, respectively. The PCR product (*pHSP_F* and *mGFP_R* primers) from Plant1 was also sequenced using the PCR primers to further confirm the presence of the construct. Note the presence of non-specific amplification products in WT gDNA in all three PCRs.

(D) PCR-genotyping of *GFP* overexpression transformants to confirm integration of the *pHSP::GFP* construct in a *P. patens* wild-type (WT) background. Top panel:

integration of the construct into the *P. patens* genomic locus 108 was confirmed using the primers *nptII_F* and *108locus5'_R* in 5 transformants that survived two rounds of G418 selection. Bottom panel: integration of the *pHSP::GFP* construct into the genome was confirmed using the primers *pHSP_F* and *35STer_R* in 4 out of 5 transformants (Plants 1, 2, 4 and 5) that survived two rounds of G418 selection. The PCR products from Plants 1, 2 and 4 in the bottom panel were also sequenced using the PCR primers to further confirm the presence of the construct. WT gDNA and the transformation vector (*pHSP::GFP*) were used as negative and positive controls, respectively, in each panel.

Figure S3.

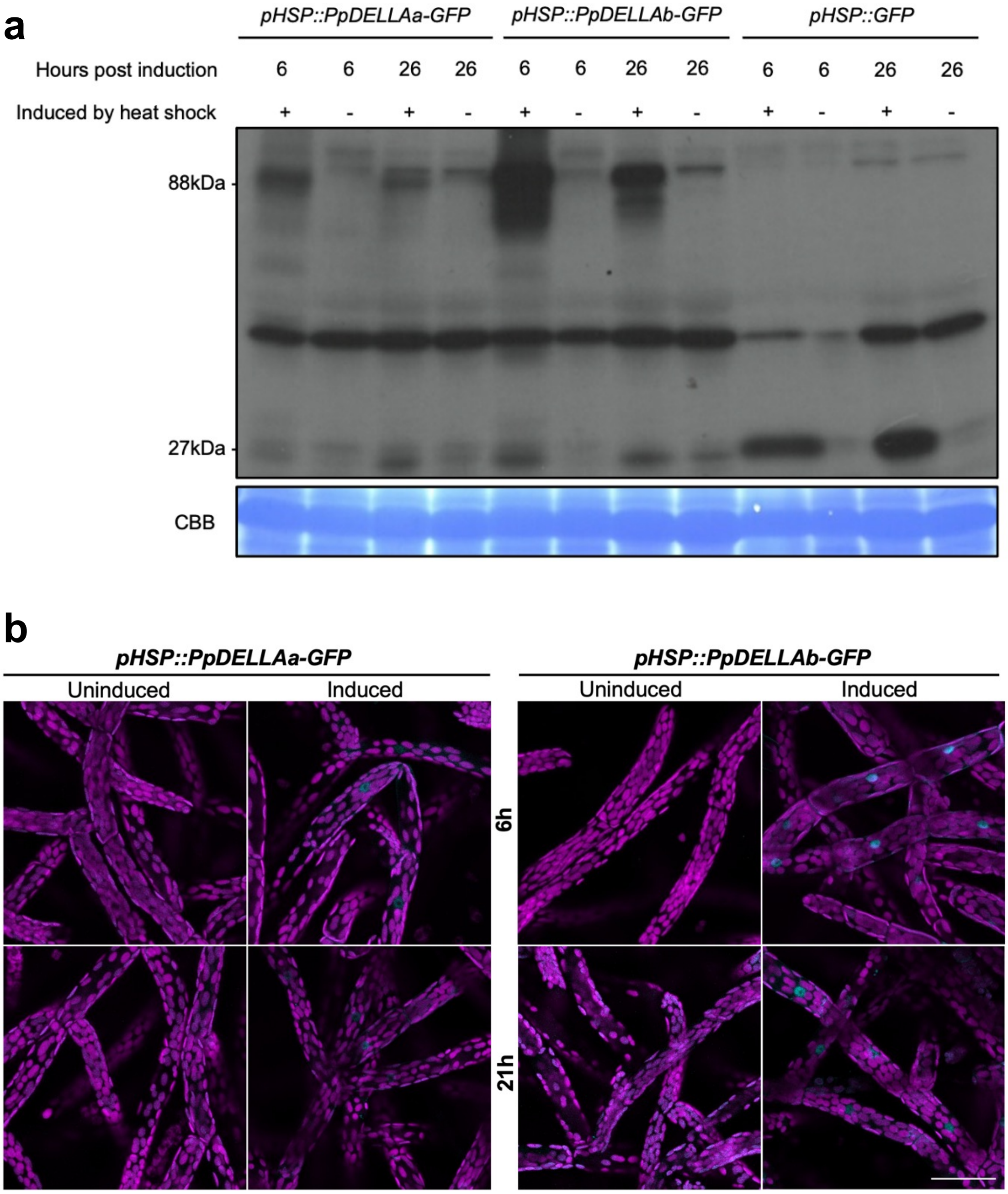


**Supplemental Figure 3. Induction of *Pp*DELLA-GFP and GFP protein expression by heat shock is sustained for at least 26 hours.**

*Pp*DELLAa-GFP (88kDa; from Plant1), *Pp*DELLAb-GFP (88kDa; from Plant1) and GFP (27kDa; from Plant2) protein expression was induced by a 1h heat shock at 37$^{o}$C.

(A) Detection using anti-GFP on a western blot 6 and 26 hours after induction. CBB, Coomassie brilliant blue staining.

(B) Confocal images showing *Pp*DELLAa-GFP and *Pp*DELLAb-GFP expression primarily in the nuclei of 7-day old *P. patens* protonemata 6h and 21h post-induction (induced). In the absence of heat shock, no *Pp*DELLA-GFP expression could be observed (uninduced). Cyan: GFP signal; Magenta: chloroplast auto-fluorescent signal. (Scale bar, 50μm).

**Figure S4.**

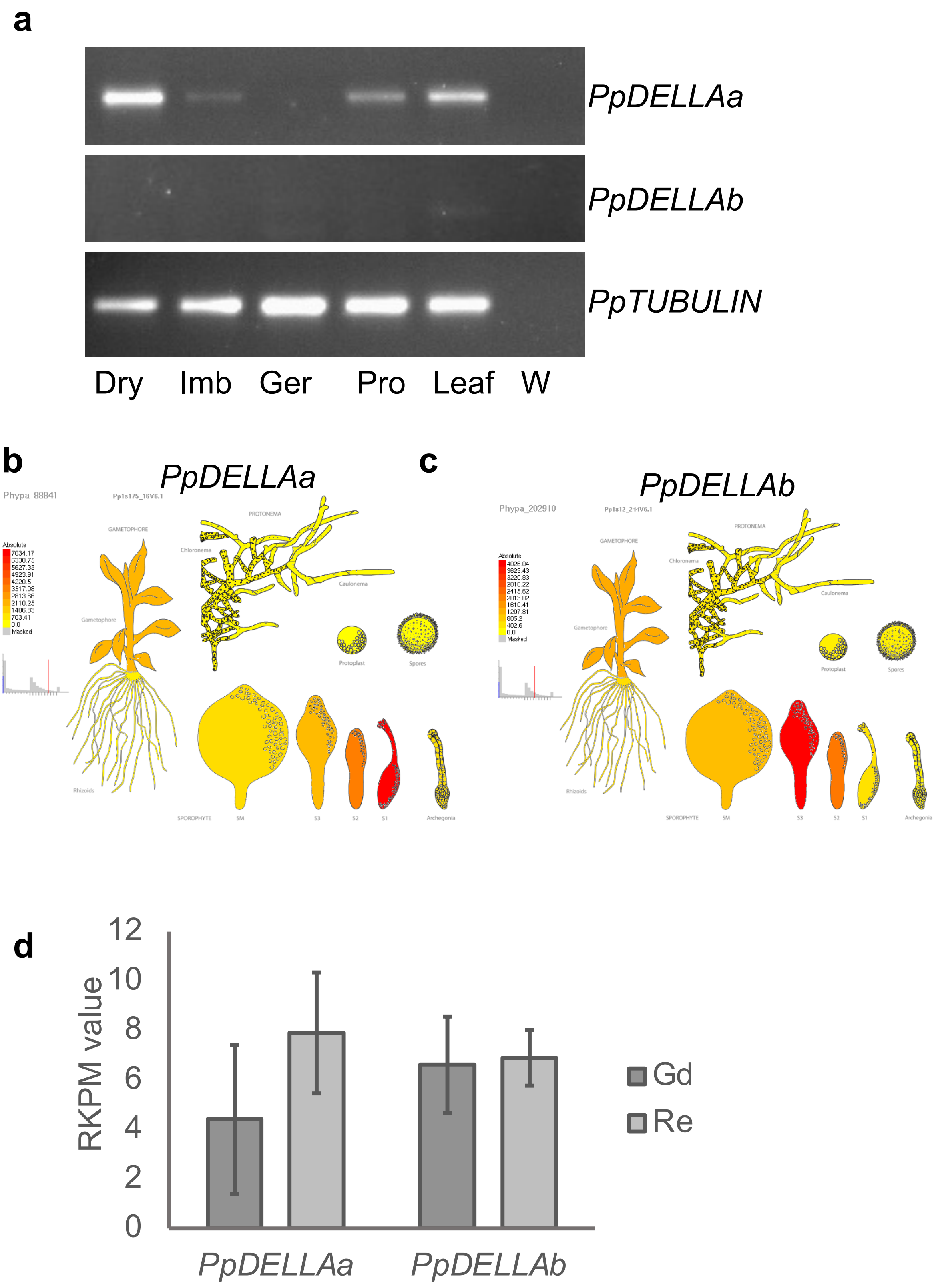
a
PpDELLAa
PpDELLAb
PpTUBULIN
Dry
Imb
Ger
Pro
Leaf
W
b
PpDELLAa
Phypa_88841
c
PpDELLAb
Phypa_202910
d
RKPM value
0
2
4
6
8
10
12
PpDELLAa
PpDELLAb
Gd
Re

**Supplemental Figure 4. *PpDELLA*s are strongly expressed in dry spores and developing sporophytes.**

(A) Using semi-quantitative RT-PCR analysis, *PpDELLAa* (639bp fragment) is detected most strongly in dry spores (Dry), with reduced expression in imbibed spores (Imb) and no expression in germinating spores (Ger). *PpDELLAa* expression is also present in protonema (Pro) and leafy tissue (Leaf), similarly to the data in panel C. *PpDELLAb* (588bp fragment) expression is much lower than *PpDELLAa* (as seen when comparing panel C and E) and is just detectable in leafy tissue, similarly to the data in panel E. Both *PpDELLA* expression profiles are compared to a *PpTUBULIN* (438bp) control. W, water negative control. Representative of 3 biological repeats.

(B) Using microarray data taken from the *Physcomitrella* eFP browser (Ortiz-Ramirez *et al.*, 2016), expression of *PpDELLAa* (*Pp3c19_8310V3.1*) is highest in the S1 stage of developing sporophytes, as shown in red (absolute value 7034.17). *PpDELLAa* expression is also present in spores from fully mature spores from the SM spore capsules (68.31) and mature sporophytes (944.73), archegonia (829.93), protonema (130.25-320.57) and leafy tissue (2040.29).

(C) Using microarray data taken from the *Physcomitrella* eFP browser (Ortiz-Ramirez *et al.*, 2016), expression of *PpDELLAb* (Pp3c22_7230V3.1) is highest in the S3 stage of developing sporophytes (4026.04), as shown in red. *PpDELLAa* expression is also present in spores from fully mature spores from the SM spore capsules (239.92) and mature sporophytes (1040.02), archegonia (196.78), protonema (208.79-376.53) and leafy tissue (1329.05).

(D) Using data from (Meyberg *et al.*, 2020), expression of *PpDELLAa* and *PpDELLAb* in antheridia bundles was plotted for both Gransden (Gd-UK) and Reute (Re) *Physcomitrium* wild type ecotypes.

**Figure S5.**

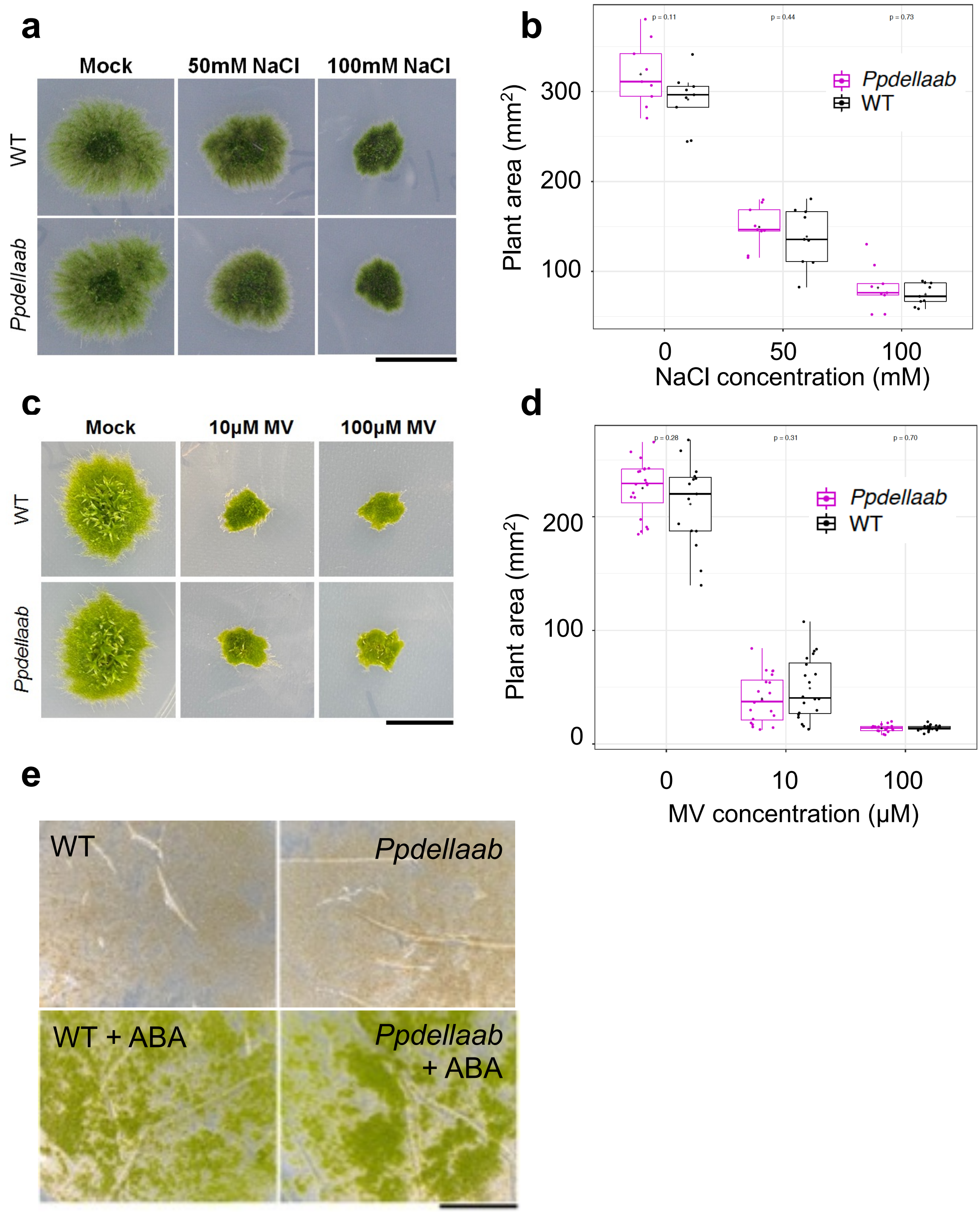


**Supplemental Figure 5. *Ppdellaab* mutants do not show altered responses to salt, oxidative or desiccation stress compared to wild type (WT).**

(A) *P. patens* protonemata grown for 15 days on different concentrations of salt (NaCl) reduces protonemal and gametophore development. Scale bar, 10mm.

(B) *Ppdellaab* plants do not have significantly different area to WT when treated with 0mM, 50mM or 100mM NaCl (n=9 per genotype). p-values were calculated via a Mann Whitney U-test: 0mM NaCl p=0.11, 50mM NaCl p=0.44, 100mM NaCl p=0.75. Black asterisks indicate the mean. To account for variability in plant area at the start of the experiment, this was subtracted from the plant area at the end of the experiment.
(C) *P. patens* protonemata treated with 1μM methyl viologen (MV) reduces gametophore differentiation and promotes protonema development, while treatment with 10μM or 100μM MV results in growth arrest. *Ppdellaab* and WT do not show obvious phenotypic differences. Scale bar, 10mm.
(D) *Ppdellaab* and wild-type plants do not have significantly different plant areas at 10μM and 100μM MV (n=20 per genotype). Differences were tested for significance ($p<0.05$) using the Mann-Whitney U test. p-values were calculated via a Mann Whitney U-test: 0mM MV p=0.20, 10μM MV p=0.31, 100μM MV p=0.70. Black asterisks indicate the mean. To account for variability in plant area at the start of the experiment, this was subtracted from the plant area at the end of the experiment.
(E) *P. patens* protonemata grown for 16 hours on cellophane-overlaid medium supplemented with 10μM abscisic acid (ABA) or methanol were then transferred (on the cellophane) into empty petri dishes for 7 days of desiccation stress. Plants were recovered by placing the desiccation-stressed tissue back under normal conditions for 7 days. Protonemata pre-treated with 10μM ABA (bottom panels) displayed desiccation stress tolerance, whereas protonemata pre-treated with methanol (top panels) did not. No difference was seen between WT and *Ppdellaab* responses. Scale bar, 20mm.

**Figure S6.**

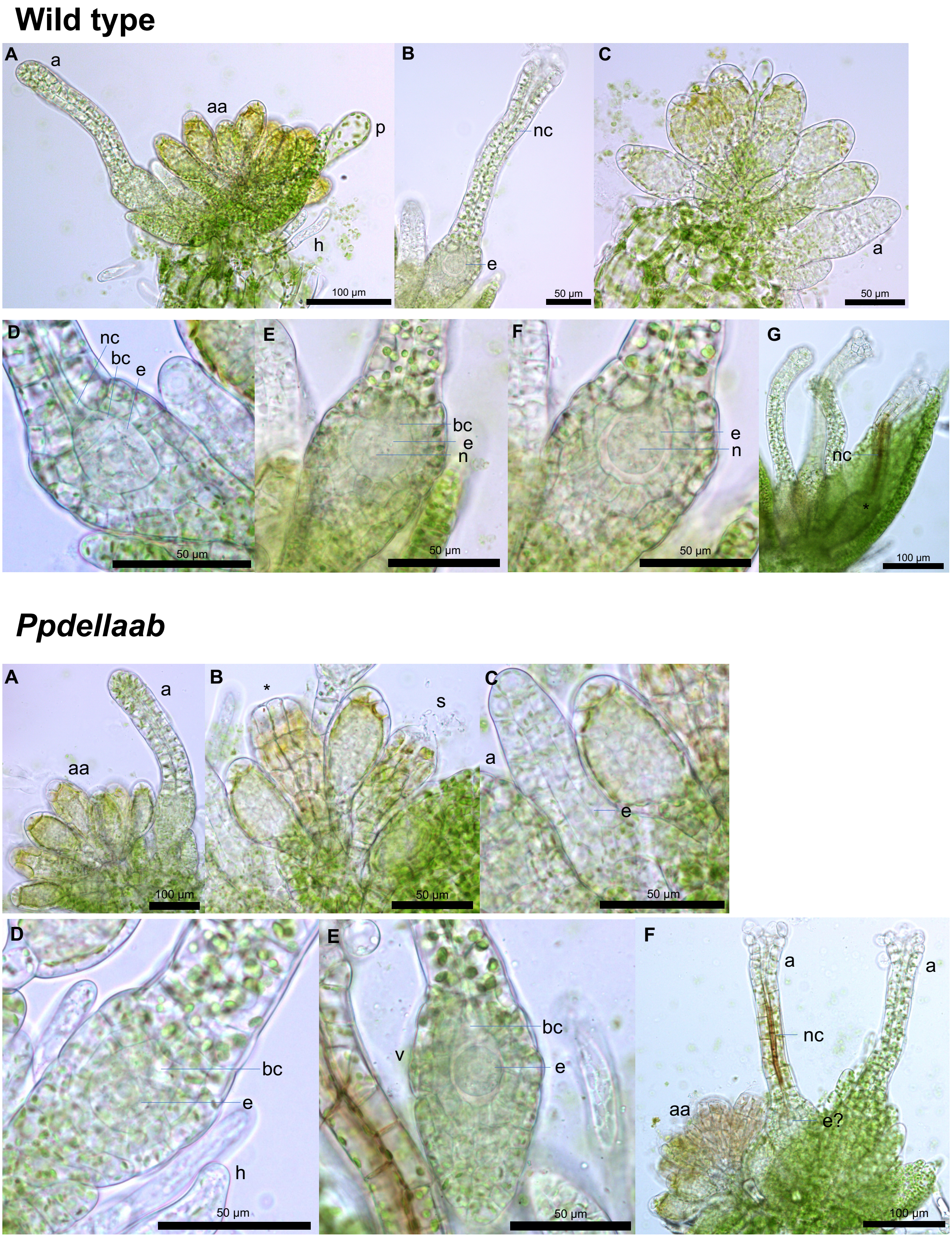


**Supplemental Figure 6. *Ppdellaab* mutants can develop antheridia and archegonia.**

**Top panel: Gametangia analysis of wild type (Gd).** A: Apex showing an immature archegonium with closed tip cells (a), mature antheridia (yellow color, swollen tip cell, aa), a paraphyse (p) and anaxilliary hair (h). B: Mature archegonium with open tip cells, dissolved neck canal cells (nc) and the mature egg cell (e). C: Antheridia bundle consisting of antheridia in different developmental stages and two newly arising archegonia (a). D: Archegonial venter showing the egg cavity with an immature egg (e) and the basal cell (bc), with visible vertical cell walls between neck canal (nc) cells, basal cell and egg. E: Archegonial venter showing the cavitiy including a maturing egg cell (e, with nucleus (n)) and basal cell (bc) (egg cell "swims“ already in the cavity and the basal cell has started to shrink. F: Mature egg cell as already shown in B with visible nucleus (n). G: Apex showing several archegonia in different developmental stages (development of gametangia typically occurs in *Physcomitrium* as long as no fertilization has taken place). Archegonium marked with * shows brown neck canal (nc), which in this time frame usually indicates the entrance of a spermatozoid into the nc; at later timepoints, this is also a sign of aging, but then the distribution of the colour is usually broader and not as distinct as is visible here.

**Bottom panel: Gametangia analysis of *Ppdellaab*.** A: Apex showing nearly mature antheridia (aa) and an immature archegonium (a). B: Bundle of antheridia including an empty antheridium (*, spermatozoids already released) and an antheridium just releasing separated spermatozoids (s). C: Early developmental stage of an archegonium (a) showing already the clear setup of the egg cell (e) in a cavity. D: Maturing archegonial venter with egg cell (e) and basal cell (e) and axilliary hairs (h). E: Nearly mature egg cell (e) with visible nucleus (n) in the archegonial venter (v) with a dissolving basal cell. F: Apex with two mature archegonia (a) showing open tip cells. Left sided archegonium shows brownish neck canal cells (nc), which indicates the entry of spermatozoids (as in the top panel). The egg cell (e) is not clearly visible, and may be shrunken. Bundle of antheridia present (aa) showing many antheridia, which have already released their spermatozoids.

**Figure S7.**

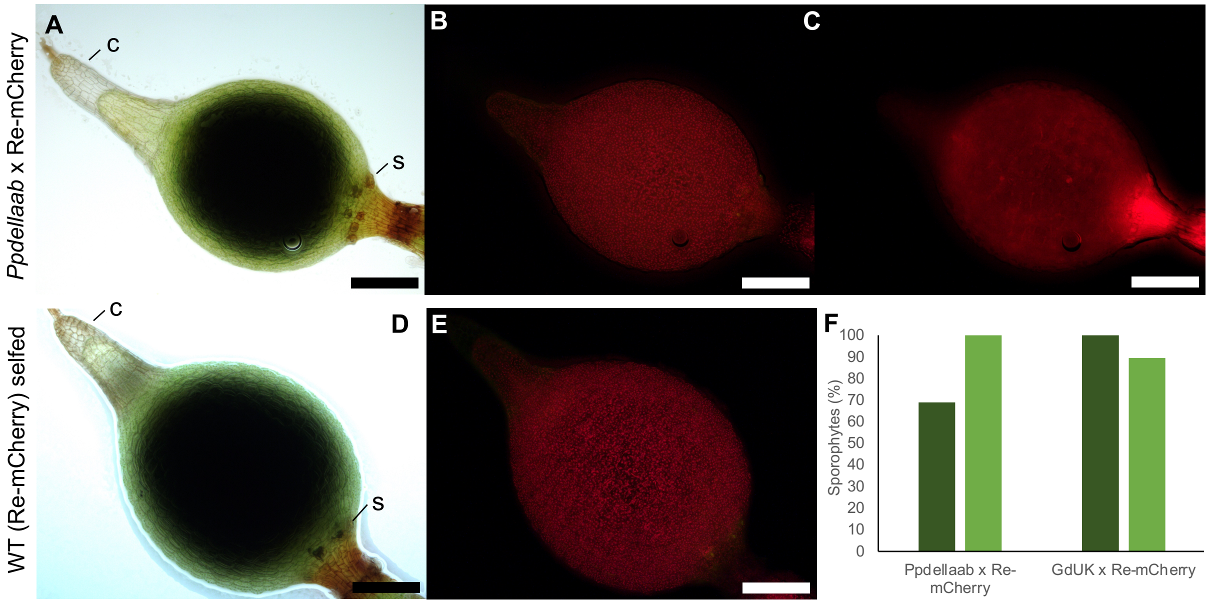


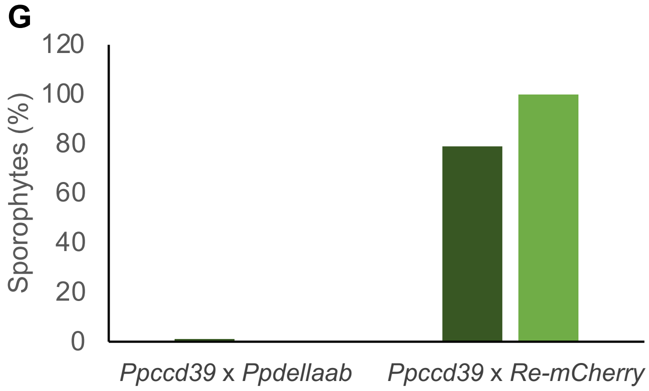


**Supplemental Figure 7. *Ppdellaab* mutants develop sporophytes when fertilized by a Reute (Re)-mCherry wild type strain but not when crossed with the male sterile mutant *Ppccd39*.**

(A) Brightfield image of crossed sporophyte of *Ppdellaab* x Re-mCherry between light brown to brown stage (LB-B, according to (Hiss *et al.*, 2017)) with the calyptra attached to the tip of the sporophyte **c**. Spores are developing in the dark area in the centre of the sporophyte. Stomata are located at the bottom of the sporophyte **s**. Stomata are distributed irregularly.

(B) Chlorophyll autofluorescence of the sporophyte in (A).

(C) mCherry fluorescence imaging of the sporophyte in (A).

(D) Re wild type sporophyte in LB-B stage. Calyptra **c** attached to the tip of the sporophyte. Stomata developed at the bottom of the sporophyte **s**.

(E) Chlorophyll autofluorescence of the sporophyte in (D).

(F) Sporophyte development under crossing conditions showed 69% sporophytes per gametophore for *Ppdellaab* (n=100; dark green bar). All developed sporophytes are products of a cross (mCherry fluorescence present; light green bar). By contrast, the background strain of the *Ppdellaab* mutant, Gd, develops 100% sporophytes per gametophore (n=104; dark green bar) with 89% of them being the product of a cross (mCherry fluorescence present; light green bar).

(G) Crossing of the male sterile mutant *Ppccdc39* with *Ppdellaab* resulted in one developed sporophyte. Resulting spores were not able to germinate, thus the genotype could not be identified. Under crossing with Re-mCherry, *Ppccdc39* developed 79% sporophytes per gametophyte (dark green har) of which 100% were crosses (light green bar).

Scale bars, 200µm.

**Figure S8.**

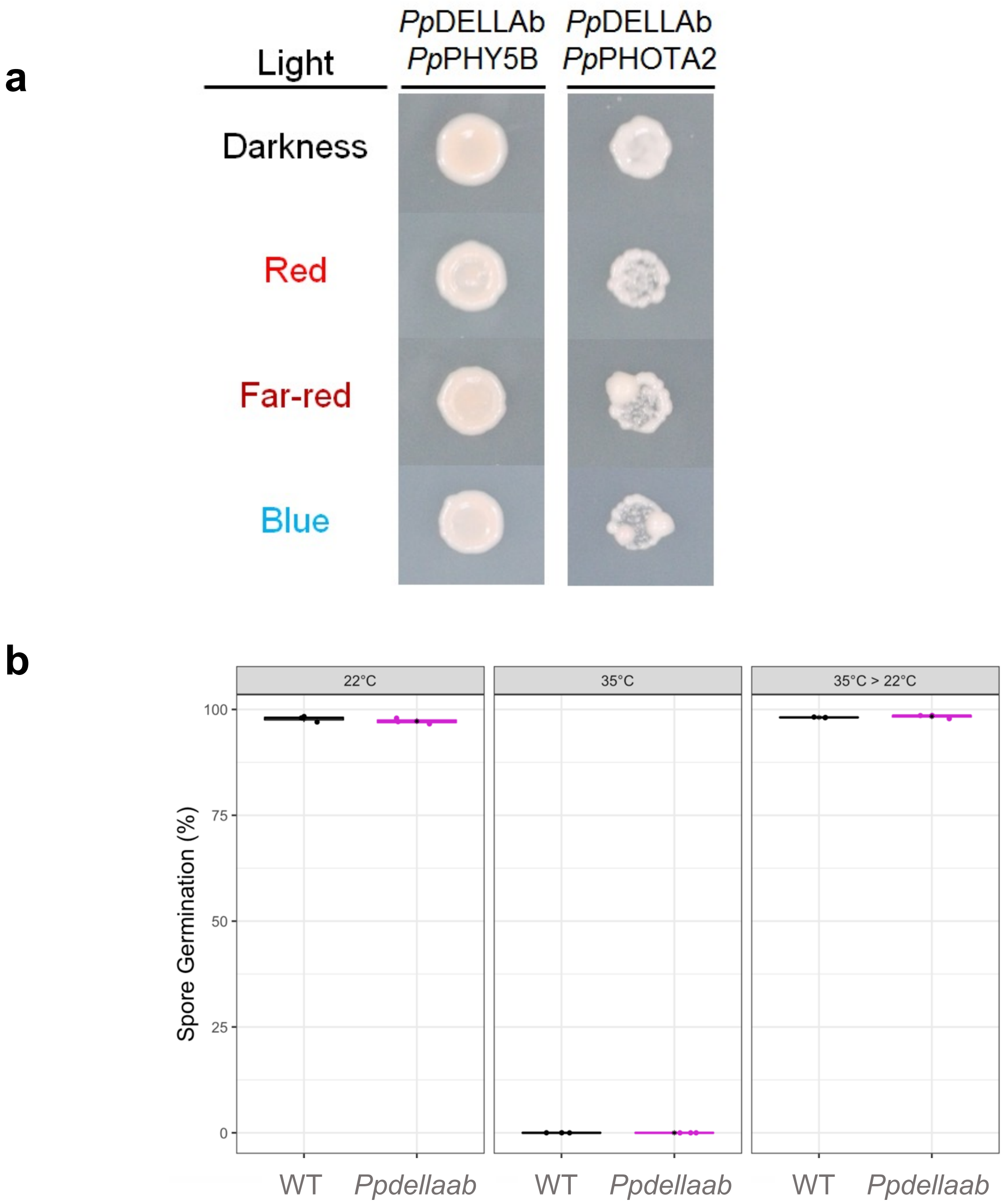


**Supplemental Figure 8. *Pp*DELLA proteins and show no differences in interaction with light receptors in yeast in response to light wavelength and *Ppdellaab* mutant spores show normal thermoinhibition.**

(A) Yeast two-hybrid assay between *Pp*DELLAb fused with the GAL4 activation domain (AD) in pGADT7 and the photoreceptors *Pp*PHY5B and *Pp*PHOTA2, fused with the DNA-binding (DBD) domain of pGBKT7 illuminated with different light wavelengths. *Pp*DELLAb interacted with both *Pp*PHY5B and *Pp*PHOTA2 in a light-independent manner. Red, 640-695nM, 5μmolm$^{-2}$s$^{-1}$; Far-red, 730nM, 3μmolm$^{-2}$s$^{-1}$; Blue, 445-490nM, 5μmolm$^{-2}$s$^{-1}$.

(B) Wild type (WT) and *Ppdellaab* mutant spores were incubated at 22˚C or at the thermoinhibitory temperature of 35˚C for 7 days. Both *Ppdellaab* and WT spores germinate fully at 22°C and do not germinate at 35˚C. Spores incubated at 35°C were then transferred to 22°C for a further 7-day period (35°C > 22°C). Both *Ppdellaab* and WT spores germinated fully upon transfer from 35˚C to 22°C. Black asterisks indicate the mean.

**Figure S9.**

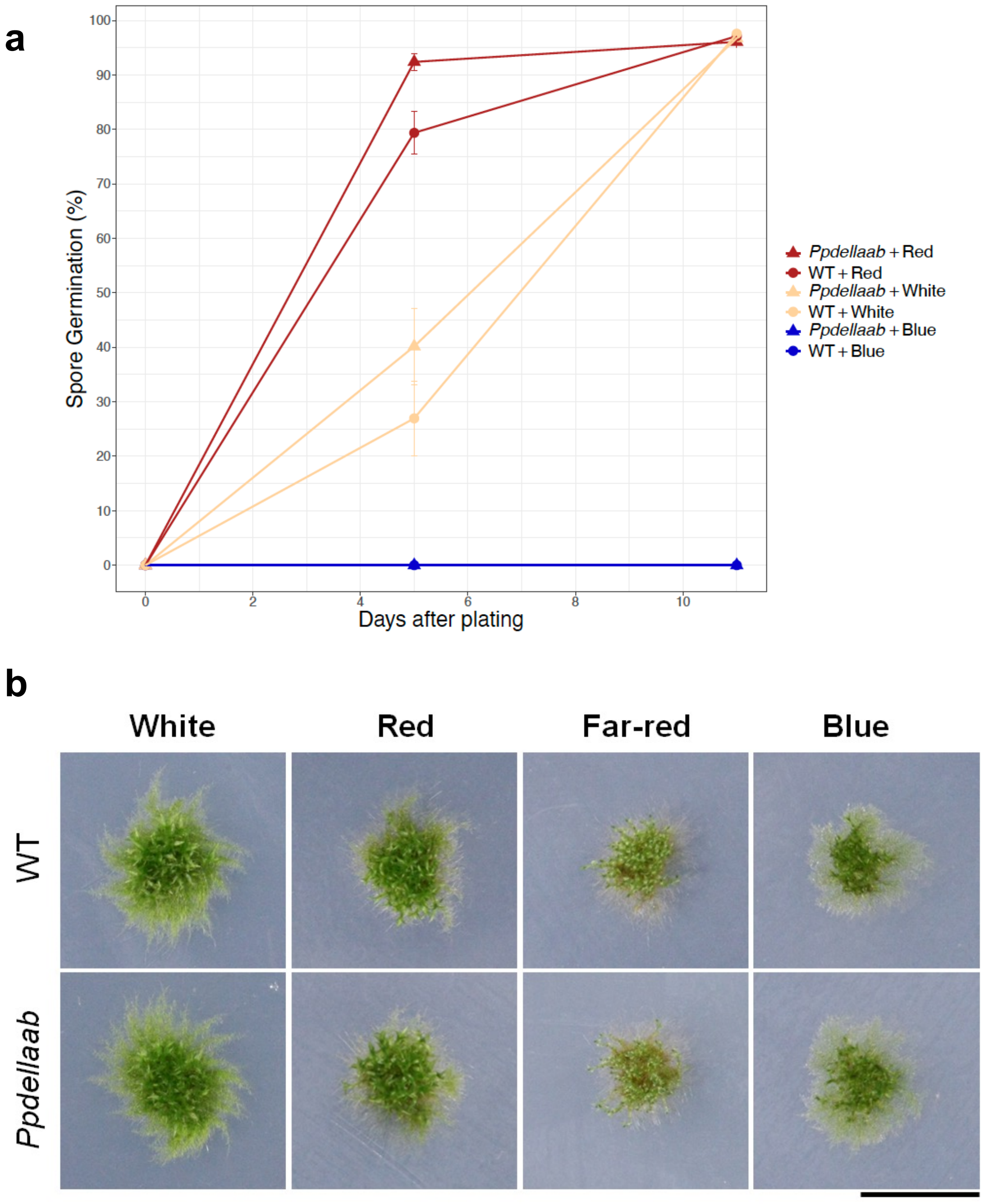


**Supplemental Figure 9. *Ppdellaab* mutants respond to different light wavelengths similarly to wild type (WT) during spore germination and vegetative growth.**

(A) Spore germination under continuous illumination with white light (63μmolm$^{-2}$s$^{-1}$), red light (640-695nM; 26μmolm$^{-2}$s$^{-1}$ intensity) or blue light (445-490nM; 16μmolm$^{-2}$s$^{-1}$ intensity) at 22$^{o}$C. The spore germination rate increases under red light compared to white light, while blue light inhibits germination in both genotypes. A Kruskal-Wallis test indicates significant differences between *Ppdellaab* + blue light and *Ppdellaab* + red light on day 5 ($p<0.01$), between *Ppdellaab* + blue light and WT + red light on days 5 and 11 ($p<0.05$), between WT + blue light and *Ppdellaab* + red light on day 5 ($p<0.01$), between WT + blue light and WT + red light on days 5 and 11 ($p<0.05$), between WT + blue light and WT + white light on day 11 ($p<0.05$), and between *Ppdellaab* + blue light and WT + white light on day 11 ($p<0.05$). Error bars, ± SEM.
(B) Moss vegetative tissue incubated for 11 days at 22$^{o}$C under continuous illumination from above with either white light (63μmolm$^{-2}$s$^{-1}$), red light (640-695nM; 26μmolm$^{-2}$s$^{-1}$), blue light (445-490nM; 16μmolm$^{-2}$s$^{-1}$) or far-red light (730nM; 16μmolm$^{-2}$s$^{-1}$). Far-red light induces etiolated growth in gametophores, which display a ‘slender’ phenotype, growing towards the light source. No differences were observed between WT and *Ppdellaab*. Scale bar, 10mm.

**Supplemental Tables 1-5**

**Table S1:** list of proteins identified as interacting with *Pp*DELLAs. 408 proteins were identified. Their unique (PANTHER) identifier assigned by the FASTA database used, description, molecular weight (MW), number of distinct peptides detected and protein FDR confidence (1% cut-off) are shown.
**Table S2:** genes downregulated in the *Ppdellaab* compared to wild type (*Pp*DELLA-induced genes) ($p<0.01$).
**Table S3:** genes upregulated in the *Ppdellaab* compared to wild type (*Pp*DELLA-repressed genes) ($p<0.01$).
**Table S4:** transcription factor binding sites enriched in the promoters of *Pp*DELLA-induced genes and *Pp*DELLA-repressed genes identified by PlantRegMap. Both the TF target genes and the identified putative TFs are listed.
**Table S5:** Primers used in this paper.

**Methods S1.**

***Physcomitrium patens* tissue culture for maintenance and spore germination analyses**
For phenotyping, BCD agar medium was supplemented with 1 mM $CaCl_2$, 5 mM ammonium tartrate and 0.5% glucose (BCDATG). For spore germination, BCD agar medium was supplemented with 5 mM $CaCl_2$ and 5 mM ammonium tartrate. For selection plates, BCD agar medium was supplemented with 1 mM $CaCl_2$, 5 mM ammonium tartrate and 50µg/ml G418 (Sigma-Aldrich, A1720).

***P. patens* tissue culture for gametangia/sporophyte and crossing analyses**
Single gametophores were inoculated for 6 weeks under long day conditions (LD, 70 µmol $m^{-2}$ $s^{-1}$ 16h light, 8h dark, 22°C) on solid KNOP medium (Knop, 1868) in 9cm petri-dishes with vents enclosed with parafilm. For gametangia induction, plates were transferred to short day (SD, 20µmol $m^{-2}$ $s^{-1}$, 8h light, 16h dark, 15°C) for three weeks. Gametangia analyses were performed at 21d after SD transfer.
For crossing analysis, gametophores were inoculated in Weck jars (Weck, Wehr-Öflingen, Germany) with 100ml of KNOP medium (Knop, 1868) and sealed with 3M

tape (3M). Plants were grown for 6 weeks under LD conditions and afterwards transferred to SD. Upon gametangia development, 14 days after SD transfer, cultures were flooded with sterile tap water for 24h. This was repeated after 21d days. Crossing analyses were carried out 2-3 weeks after watering using green sporophytes to easily detect mCherry fluorescence of crosses.

***P. patens* genomic DNA extraction**

*P. patens* tissue was ground up in Eppendorf tubes using sterile micropestles, resuspended in 700µl cetyltrimethyl ammonium bromide (CTAB) buffer (100mM Tris-HCl (pH 8.0), 20mM EDTA (pH 8.0), 1.4M NaCl, 2% (w/v) CTAB, 1% polyvinyl pyrrolidone 40,000) and incubated at 65°C for 1h. 1 volume of chloroform was added and vigorous shaking was applied. This was followed by centrifugation for 10 minutes at 14,000 g, transfer of 500µl of the upper aqueous layer to a 2ml Eppendorf tube and addition of 0.8 volumes of isopropanol. The mixture was then incubated for at least 2h at -20°C, centrifuged at 14,000 g for 20 minutes and pellets washed twice with 70% ethanol for 10 minutes and air-dried. DNA was eluted in 30-50µl nuclease free water.

**SDS-PAGE and Western blotting**

Primary and secondary antibodies were diluted in 10 ml 5% (w/v) Marvel semi-skimmed milk in TBST (50mM Tris, 150mM NaCl, pH 7.5 with 1M HCl, 0.1% tween). Mouse monoclonal α-HA (Abcam, ab130275) and α-MYC (Abcam, ab18185) were used at 1:2000 dilution and incubations were performed for 3h and 1h respectively at room temperature or overnight at 4°C. Rabbit polyclonal anti-GFP (Chromotek, Germany, PABG1) was used at 1:1000 or 1:500 dilution and incubations were performed at 4°C overnight. Three 5-minute washes in TBST were performed before probing with secondary antibody. Goat anti-mouse immunoglobin (Abcam, ab6789) was used for α-MYC and α-HA, and goat anti-rabbit immunoglobin (Abcam, ab6721) for α-GFP. Secondary antibodies were used at a 1:2000 dilution in 5% (w/v) milk in TBST and incubations were carried out for 1.5h.

**Yeast two-hybrid assays**

2μg of pGBKT7 and pGADT7, empty or carrying the construct of interest, was used for yeast transformation in TB buffer. Yeast colonies growing on synthetic amino acid Drop out (DO) -leu-trp (Formedium, DSCK172) agar were resuspended in 150μl nuclease free water and 5μl of the mixture was transferred on both DO -leu-trp-his-ade (Formedium, DSCK272) and DO -leu-trp agar plates. For assays testing DELLA-GID1 homologue interactions, DO -leu-trp-his-ade agar media were left to cool down to 50°C after autoclaving and were then supplemented with $GA_3$ or $GA_9$-ME or *ent*-kaurenoic acid or methanol before being poured into plates. For assays testing *Pp*DELLA interactions with photoreceptor proteins, selective agar plates were incubated upright at 30°C in blue (5μmolm$^{-2}$s$^{-1}$) or red (5μmolm$^{-2}$s$^{-1}$) or far-red (3μmolm$^{-2}$s$^{-1}$) light or in darkness for 4 days. Three biological replicates of each yeast two-hybrid assay were performed and plates were photographed using a Nikon D40 SLR camera.

**Co-Immunoprecipitation (Co-IP) in a cell-free system**

40U RNaseOut (Invitrogen) per 50ml reaction was used to inhibit ribonucleases. For each Co-IP, 15μl protein-A sepharose magnetic beads (Amersham), pre-washed three times with 1ml IP Buffer A (50mM HEPES pH 7.5, 150mM NaCl, 5% [v/v] glycerol, 0.1% Tween 20, cOmplete™ EDTA-free protease inhibitor tablets [Roche] - one per 10ml buffer), were incubated with 4μg α-MYC (Abcam, ab18185) and 250μl IP Buffer A for 1h at room temperature on a turning wheel. This was followed by 3 three-minute washes with 1ml IP Buffer A. Co-IPs were performed by adding 9μl translated proteins to the MYC-coupled beads and mixing in a total volume of 500μl IP buffer A supplemented with $GA_3$ or $GA_9$-ME or methanol at 4°C for 3h on a turning wheel. This was followed by 4 three-minute washes with 1ml IP Buffer B (50mM HEPES pH 7.5, 300mM NaCl, 5% [v/v] glycerol, 0.1% Tween 20, cOmplete™ EDTA-free protease inhibitor tablets [Roche] - one per 10ml buffer), and a three-minute wash with 1ml IP Buffer A. Samples were resuspended in 50μl 1x Laemmli buffer (2% [w/v] SDS, 10% [w/v] glycerol, 1% β-mercaptoethanol, 0.001% [w/v] bromophenol blue), boiled for 10 minutes at 95°C and stored at -20°C. Samples were analysed by SDS-PAGE and Western blotting.

***P. patens* spore culture and germination assays**

For spore thermoinhibition assays, plates were incubated at 35°C with a 16h:8h light:dark cycle for 7 days and returned to 22±1°C with a 16h:8h light:dark cycle for 7 days. All assays were performed with a light intensity of 50-70μmolm$^{-2}$s$^{-1}$. For hormone treatment assays, BCD agar medium supplemented with 5 mM $CaCl_2$ and 5 mM ammonium tartrate and after autoclaving, was cooled down to 50°C, supplemented with the required hormone or solvent and poured into plates. Methanol was used as solvent for abscisic acid (ABA) (Sigma-Aldrich, A1049), gibberellin $A_3$ ($GA_3$) (Sigma-Aldrich, 48880), gibberellin $A_9$ methyl ester ($GA_9$-ME) and *ent*-kaurenoic acid. $GA_9$-ME and *ent*-kaurenoic acid supplies were kindly provided by Professor Peter Hedden (Rothamsted Research, UK).

**Immunoprecipitation coupled to mass spectrometry**

Each immunoprecipitation was performed in 15ml falcon tubes with 40μl GFP-trap® magnetic agarose beads for 90 minutes at 4°C on a rotating wheel (half speed). Following immunoprecipitation, working in a cold room, the beads were washed twice (5 minutes each) with a high salt buffer (10mM Tris/HCl pH 7.5, 0.5mM EDTA, 400mM NaCl, cOmplete™ EDTA-free protease inhibitor tablets [Roche] - one per 10ml buffer), followed by two washes (5 minutes each) with dilution buffer (kit's own). Beads were then resuspended in 50μl 2x Laemmli buffer (4% [w/v] SDS, 20% [w/v] glycerol, 2% β-mercaptoethanol, 0.002% [w/v] bromophenol blue), boiled at 95°C for 10 minutes and the elution was stored at -20°C.
